# Phenotypic CRISPR screening identifies ZBTB10 as a novel regulator of human trophoblast differentiation

**DOI:** 10.64898/2026.06.29.734890

**Authors:** Meagan N. Esbin, Rituja Bhowmik, Hanqin Li, Andrew Cearlock, Jiayu Chen, Tomy Tran, Stephen A. McCartney, Ian A. Glass, Birth Defects Research Laboratory (BDRL), Dirk Hockemeyer, Xavier Darzacq, Fyodor Urnov, Robert Tjian, Min Yang

**Affiliations:** Department of Obstetrics & Gynecology, University of Washington, Seattle, USA; Edward and Pearl Fein Memory and Aging Center, Weill Institute for Neurosciences, University of California San Francisco, San Francisco, USA; Innovative Genomics Institute, Berkeley, USA; Department of Molecular and Cell Biology, University of California Berkeley, Berkeley, USA; Department of Pediatrics, University of Washington, Seattle, WA, USA; Howard Hughes Medical Institute, Berkeley, USA; Brotman Baty Institute for Precision Medicine, Seattle, USA; Institute for Stem Cell and Regenerative Medicine, Seattle, USA; Washington National Primate Research Center, Seattle, USA; Department of Genome Sciences, University of Washington, Seattle, USA

## Abstract

The human placenta is built by trophoblast cells that fuse together, secrete hormones, and invade the uterus, and defects in these processes contribute to pregnancy disorders such as preeclampsia. Because cell-cell fusion and hormone secretion are inherently non-cell-autonomous processes, their regulators have remained inaccessible to conventional pooled CRISPR screens. Here, we developed an arrayed CRISPR screen in fusogenic BeWo trophoblasts that simultaneously quantifies fusion and hCG secretion across 412 gene perturbations. The screen revealed that these two hallmark functions of trophoblast differentiation are genetically separable. We characterized the strongest novel hit, ZBTB10, in trophoblast stem cells, organoids, and placental tissue and find that ZBTB10 is an essential regulator of human trophoblast differentiation. ZBTB10 is required for invasive extravillous trophoblast differentiation and supports syncytiotrophoblast maturation, establishing it as a cross-lineage regulator that both activates and represses distinct trophoblast fate programs. Together, these findings provide a genetic platform and phenotypic dissection of how regulatory networks control human placental development.

## INTRODUCTION

The placenta is a critical, temporary organ that sustains human pregnancy.^1,2^ Human placental development depends on the differentiation of cytotrophoblasts (CTBs, the progenitor stem cell population within the human placenta) into two downstream specialized lineages: extravillous trophoblasts (EVTs) invade the uterus to remodel spiral arteries to establish adequate blood flow to the placenta and syncytiotrophoblasts (STBs) form a multi-nucleated syncytium that covers the placental villi and mediate nutrient exchange, endocrine signaling, and barrier function at the maternal fetal interface.^3–5^ Defects in either lineage contribute to major pregnancy complications, including preeclampsia, fetal growth restriction, implantation failure, and miscarriage.^6–12^

Among trophoblast lineages, STBs undergo a highly unusual differentiation program in which CTBs fuse as they differentiate. Cell-cell fusion drives the formation of the placental syncytium, which expands throughout gestation ultimately resulting in a single syncytialized layer that contains ∼60 billion nuclei at term.^13,14^ In the human placenta, this process is driven primarily by Syncytin-2 (*ERVFRD-1*) which binds to its receptor MFSD2A on opposing cells to mediate fusion.^15–19^ In parallel with fusion, STBs acquire endocrine function centered on secretion of hCG, a hormone which stimulates the maternal corpus luteum to produce progesterone to sustain pregnancy, and is thought to reinforce trophoblast fusion by triggering Protein Kinase A (PKA) signaling to activate the transcription factor GCM1 upstream of the syncytin fusion machinery.^20–22^ Cell-cell fusion and hCG secretion are therefore two defining hallmarks of syncytiotrophoblast differentiation and while they are often assumed to be tightly coupled, studies using single genetic or drug-based perturbations in both BeWo cells, a fusogenic choriocarcinoma trophoblast cell line, and human trophoblast stem cells (hTSCs), have yet to produce a consensus on their relationship.^10,19,23–29^ Defining the separability and shared or distinct genetic programs driving fusion and endocrine differentiation could open avenues to detect and treat specific aspects of syncytiotrophoblast dysfunction in pregnancy. Furthermore, defining the factors that govern STB function may also reveal broader programs that coordinate and balance trophoblast lineage specification across placental development.

CRISPR screening has become a powerful tool for systematically identifying and characterizing regulators of cellular processes, both *in vitro* and *in vivo*.^30–32^ Recent CRISPR screen modifications using single-cell sequencing (e.g., CROP-seq, Perturb-seq) or imaging-based readouts are dramatically expanding genotype-phenotype profiling beyond simple survival and fluorescence-based sorting.^33–35^ However, despite this progress and a clear biological need, syncytializing trophoblasts presents a unique challenge to the traditional CRISPR screen. As both fusion and hormone secretion are inherently cell non–autonomous phenotypes, pooled approaches, which assume cell-intrinsic readouts, are poorly suited to these processes. Rare examples of fusion-compatible screening approaches have either required that fusion can occur rapidly in suspended microdroplets (for example with viral fusion), or have selectively depleted syncytialized cells (via filtration or toxic split-intein expression) from sequenced pools, eliminating gain-of-fusion phenotypes and dramatically reducing sensitivity because of the mixing of sgRNAs within a single syncytium.^36–38^ Finally, due to the inability of any of these screening methods to simultaneously measure hormone production and fusion, previous trophoblast studies have relied on candidate strategies that lack high-throughput capability.

Here, to address this limitation, we developed a high-throughput, arrayed CRISPR screen that leverages the scalability and short Forskolin (FSK)-induced differentiation window of BeWo cells to simultaneously quantify trophoblast fusion and hCG secretion across 412 candidate regulators. The screen revealed that fusion and hCG production are controlled by largely distinct regulatory programs. Importantly, we identified the transcription factor ZBTB10 as a novel regulator of STB differentiation. We systematically characterized ZBTB10’s role in physiologically relevant human trophoblast stem cells, organoids, and primary placental tissue, and found that it both supports STB maturation and is required for EVT differentiation. As a cross-lineage regulator of human trophoblast differentiation, we also find evidence that ZBTB10 placental expression is conserved across eutherian mammals, despite the evolutionary divergence of placentation across species, underscoring its importance in placental invasion.

## RESULTS

### Scalable BeWo-based arrayed CRISPR screen pairs cell–cell fusion with hCG secretion

Syncytiotrophoblast formation couples two defining outputs of human placental differentiation–cell–cell fusion and secretion of the pregnancy hormone hCG– both of which act non-cell-autonomously and are therefore inaccessible to pooled genetic screens. To dissect their regulation systematically, we established an arrayed CRISPR-knockout screen in fusing BeWo trophoblasts in which high-throughput live-cell imaging of syncytial fusion alongside ELISA measurements of secreted hCG from the same wells (Fig. 1A). Using RNA-Sequencing data from BeWo cells, we selected 412 functionally relevant gene candidates based on highly induced genes during fusion and expressed transcription factors (Fig. 1B, Fig. S1A). This library covers a range of transcriptional regulators, membrane proteins, and other relevant trophoblast genes (Fig. S1C), each targeted by 1-3 sgRNAs. Each plate carried negative (the non-essential TRAC gene and no sgRNA, Forskolin only wells) and positive (*ERVFRD-1*, essential for BeWo fusion, and the master TF regulator *GCM1*, required for both fusion and hCG) controls^10,16,39^, plus undifferentiated DMSO wells (Fig. S1B). To establish a CRISPR screening platform capable of measuring cell-cell fusion and hCG secretion at scale, we developed a co-culture assay in which GFP- and mScarlet-labeled BeWo cells were mixed at a 50:50 ratio, edited using Cas9 RNPs delivered via nucleofection, differentiated for 48 hours using Forskolin, and assayed by 4-channel live-cell confocal imaging and supernatant sampling for hCG ELISA (Fig. 1C). TIDE analysis confirmed an average knockout (KO) score of ∼80% in a scalable 96-well format (Fig. S1D).

**Figure 1.**
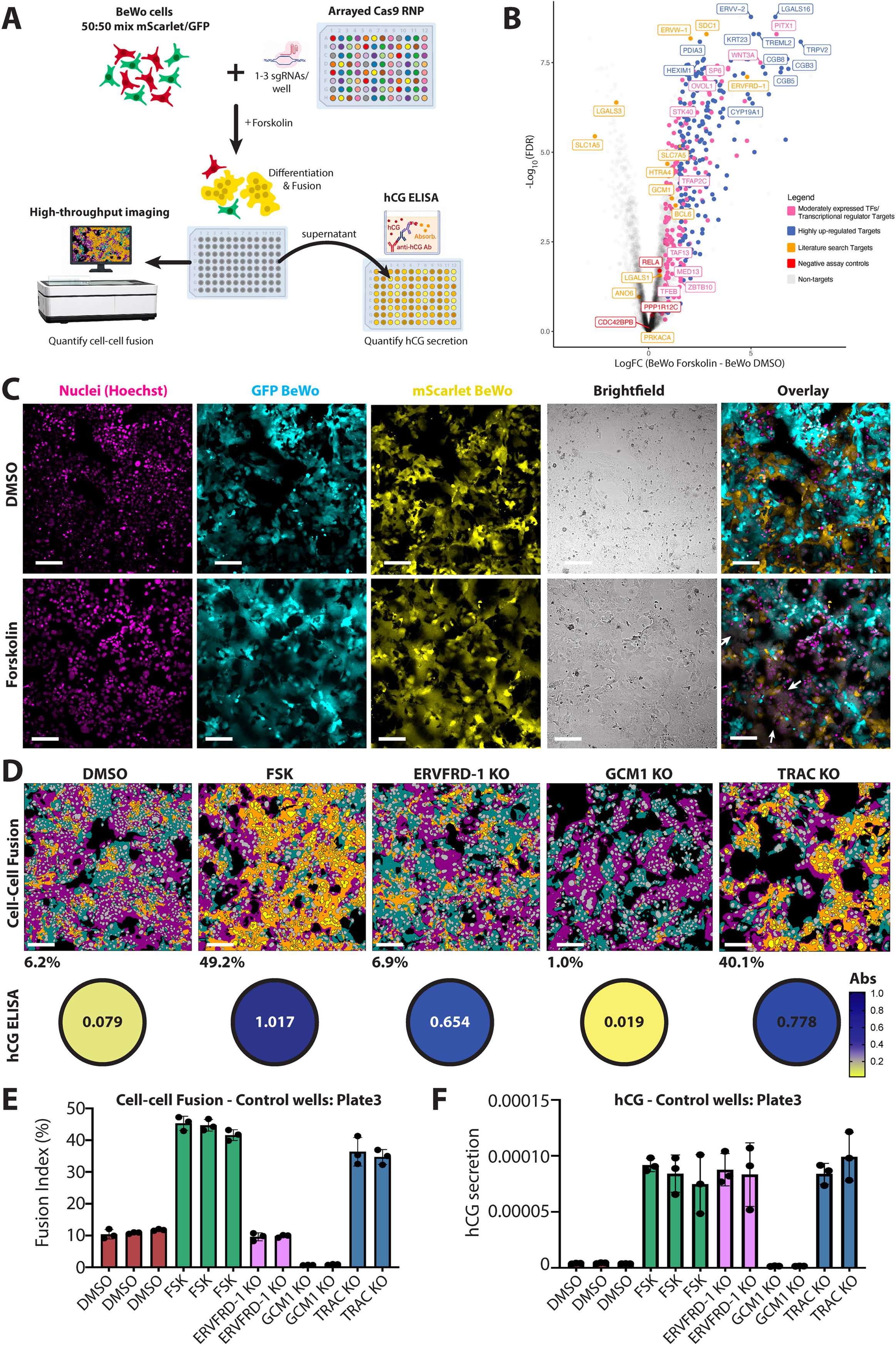
Scheme and controls for an arrayed CRISPR Screen for cell-cell fusion and hCG secretion. **A)** Overview of the arrayed, phenotypic CRISPR screen with paired readouts. **B)** The 412 candidate gene targets selected by RNA-Seq of differential gene expression in Forskolin-differentiated WT BeWos versus DMSO-treated BeWo from Esbin, et al. 2023. Only protein-coding genes with a mean RPKM > 10 in Forskolin-treated BeWo cells and targetable by Synthego are displayed (grey points) and selected gene targets for screening are shown in color. Full selection scheme outlined in Fig. S1A. **C)** Example of four-channel raw imaging data from the screen in control (DMSO) and Forskolin-treated wells. White arrows indicate syncytial regions that are positive for both GFP and mScarlet. **D)** Example paired measurements for individual control wells in the screen. Segmented overlays for a single field of view show the segmented GFP+ region (teal), the segmented mScarlet+ region (magenta), overlap between GFP and mScarlet regions (orange), double positive nuclei (yellow) and total fluorescent nuclei (gray). Calculated Fusion Index is the percentage displayed below each image. On bottom, raw hCG ELISA absorbance is shown for individual corresponding wells in the screen. **E)** Raw Fusion Index measurements across one example parental screen plate are shown. Points represent individual measurements for each well across triplicate imaging plates. Bars represent the mean; error bars show standard deviation. **F)** Raw hCG measurements (ELISA absorbance divided by the number of nuclei per well) across one example parental screen plate. Scale bars are 200μm.

Forskolin treatment increased the fusion index from ∼10% to ∼40% and hCG secretion >10-fold, recapitulating syncytiotrophoblast differentiation, and all controls behaved as expected: ERVFRD-1 and GCM1 loss suppressed fusion (TRAC had no effect), whereas only GCM1 loss suppressed hCG, in each case dropping to levels at or below the undifferentiated DMSO control (Fig. 1D). These effects were highly reproducible across technical replicates and parental wells (Fig. 1E,F, Fig. S1E,F); even before normalization, per-well triplicate correlations were high (R² ∼0.9 for fusion, ∼0.7 for hCG; Fig. S2A). Positive and negative controls demonstrated high precision and expected behavior in both fusion and hCG assays, with some variability between plate sets (Fig. S1E,F). To correct for plate-to-plate differences in global differentiation efficiency, each plate was internally normalized using a percent inhibition strategy to its own DMSO (negative control, set to 0%) and FSK/TRAC wells (positive control, set to 100%) and wells with predominantly dead cells were excluded from analysis (<45% fluorescent cells, Fig. S2B-D).

To establish reproducibility in an independent setting and validate the findings from our screen, we repeated the hCG screen in full biological replicate by high-throughput robotic automation at the Innovative Genomics Institute; controls again performed as expected, and replicate correlations were high (R² = 0.68–0.86, Fig. S3A-C). Using the same normalization and Robust Z-score MAD ± 2 hit calling, we observed significant overlap between hCG hits called in the two screens with the 16 overlapping hits including *GCM1*, *OVOL1*, *TFAP2C*, *ZBTB10* and *EGLN1* as major candidates for hCG regulation (Fig. S3D). Together, these data establish a quantitative, reproducible platform for dissecting human syncytiotrophoblast differentiation.

### Paired screening identifies ZBTB10 among separable regulators of trophoblast differentiation

Jointly profiling fusion and hCG across all 412 perturbations showed that, although both are hallmarks of syncytiotrophoblast differentiation, they are genetically separable: while most knockouts clustered near controls, knockout of the *CGB* hormone genes reduced hCG without affecting fusion, *ERVFRD-1* loss reduced fusion without affecting hCG, and hits populated all four quadrants of the fusion–hCG landscape (Fig. 2A). This allowed us to identify genes that can affect each phenotype independently. Applying a Robust |MAD Z-score| ≥ 2 cutoff, we identified 117 potential regulators of STB differentiation, comprising 86 hits for cell-cell fusion, 45 for hCG secretion, and 23 for phenotype Differentiation Score (Fig. 2B). Large-effect knockouts showed concordant fusion changes in individual images, validating our hit calling accuracy (Fig. 2C, S4A). As differentiation entails coordinated morphological changes, we used a Cell Painting approach to analyze hundreds of subcellular features from our live imaging data and project each well onto a principal-component axis separating DMSO from FSK/TRAC controls, allowing us to calculate an unbiased Differentiation Score that reflects phenotypic shifts towards or away from differentiation (Fig. 2D,E, S4C–G). In our screen, known regulators behaved as anticipated, validating the screen: GCM1 scored as a hit for all 3 phenotypes, ERVFRD-1 for fusion alone, and the key transcriptional regulators OVOL1 and TFAP2C strongly for both.^4,40–43^ Integrating all three readouts, we identified six candidates with the strongest effects on differentiation: GCM1, OVOL1, ZBTB10, STK40, EGLN1, and ZNF292 (Fig. 2F). A large gain-of-function phenotype was observed with knockout of the known fusion inhibitor STK40, increasing both fusion and hCG (Fig. 2A). Surprisingly, ZBTB10, a zinc-finger and basic helix-loop-helix transcription factor with no previously known roles in placental development^44^, emerged as the strongest loss-of-function hit for cell-cell fusion, whose loss also reduced hCG to 36% of differentiated controls (Fig. 2A). While GCM1, OVOL1, and STK40 are established regulators of syncytiotrophoblast differentiation, ZBTB10, EGLN1, and ZNF292 have not previously been characterized in this context.

**Figure 2.**
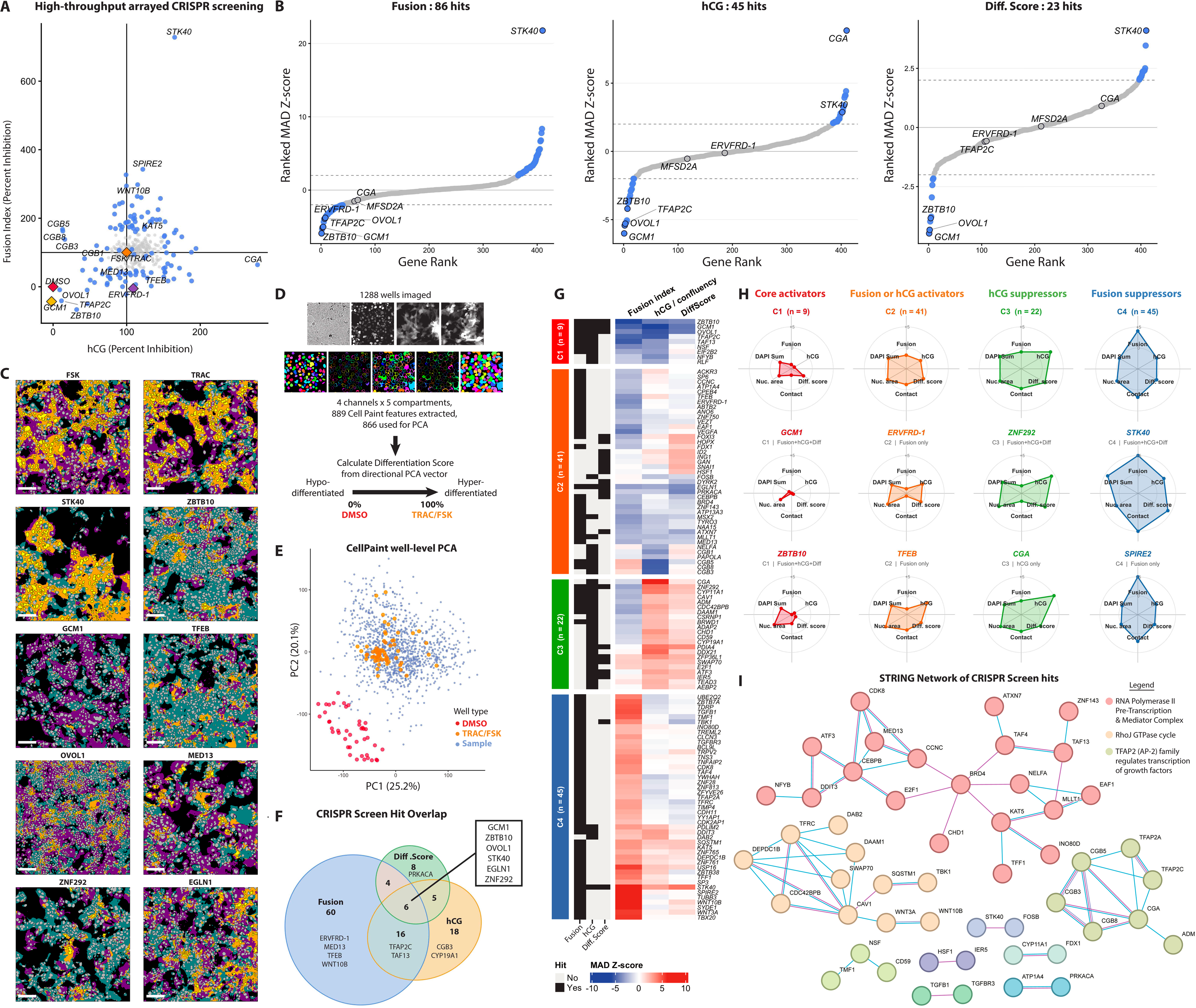
High-throughput arrayed screening reveals regulators of cell-cell fusion, hormone secretion, and differentiation phenotype. **A)** Percent inhibition-normalized per-well measurements of hCG secretion and cell-cell fusion across the screen. Controls included on every plate are indicated in colored diamonds. Hits (+/- 2 MAD Z-Score) called independently for either hCG secretion, cell-cell fusion, or Differentiation Score are shown in blue. **B)** Ranked for Fusion Index, hCG secretion, and Differentiation Score with hits (blue points) and cutoff +/-2 MAD Z-Score (grey dashed lines) shown. **C)** Cell-cell fusion phenotypes for example gene knockouts. Segmented overlays for a single field of view show the segmented GFP+ region (teal), the segmented mScarlet+ region (magenta), overlap between GFP and mScarlet regions (orange), double positive nuclei (yellow) and total fluorescent nuclei (gray). Scale bars are 200μm. **D)** Overview of Cell Paint analysis and Differentiation Score calculation. Four individual channels (top: brightfield, DAPI, GFP, mScarlet) and 5 segmented cell compartments (bottom: nucleus, nuclear ring, cytoplasm, membrane, cell) are used to calculate morphology, texture, and intensity-based parameters for extensive phenotyping. **E)** Principal component analysis of wells annotated with Cell Paint phenotypes, colored by DMSO (red) and FSK/TRAC (orange) controls. **F)** Venn diagram of hits called for each phenotype. **G)** Heatmap of Z-scores (winsorized +/- 10) for hits called across all three phenotypes, grouped using unsupervised clustering. **H)** Radar plots of per-gene Z-scores (winsorized +/- 5) for different phenotypes measured across the screen. The top radar plot shows the average across all genes in each cluster with individual gene examples shown below. DAPI Sum = summed nuclear DAPI intensity, Nuc. Area = Nuclear Area, Contact = ++ nuclei nucleus-nucleus contact area %. **I)** STRING network of all 117 CRISPR hits (displaying interactions based on Experiments and Databases only; confidence interaction score >0.40). Disconnected nodes are not displayed. Related pathways are grouped using K-means clustering (clusters = 9), colored by cluster, with the three main clusters (genes > 3) indicated in the legend.

To organize these regulators by function, we performed unsupervised clustering of the hits across all three phenotypes, which resolved four classes: 1) core activators whose knockout suppresses both hCG secretion and cell-cell fusion, 2) single-phenotype activators affecting one readout, 3) hCG suppressors whose knockout raises hCG, and 4) fusion suppressors whose knockout increases fusion efficiency (Fig. 2G-H). Reassuringly, the classification recovered functions of established trophoblast regulators, TFEB (a fusion-specific Cluster 2 gene) and the known fusion inhibitor STK40 (Cluster 4), and placed CGA among hCG suppressors (cluster 3), consistent with our ELISA detecting the free hCGβ subunit released when its dimerization partner is lost.^45–48^

The core-activator cluster (cluster 1), which shows the strongest defects across syncytiotrophoblast differentiation, includes OVOL1, GCM1, TFAP2C and ZBTB10. Protein-protein interaction analysis linked the screen hits to Pol II and Mediator, the Rho GTPase cycle, and AP-2-regulated steroidogenesis as important modules of regulation in syncytiotrophoblast differentiation yet highlighting ZBTB10 as a factor with few known interaction partners and an unknown placental function (Fig. 2I). These results identify ZBTB10 as a novel regulator of human trophoblast differentiation and given its strong effects on fusion, hCG secretion and morphology, we selected it for in-depth functional analysis.

### ZBTB10 is required for BeWo fusion and hCG secretion

To dissect ZBTB10 function, we first characterized ZBTB10 in BeWo cells. We generated polyclonal knockouts with the three library sgRNAs and confirmed 100% indel efficiency and a knockout score of 93% by TIDE (Fig. 3A). Western blotting confirmed that ZBTB10 protein levels were reduced to <10% of wild-type levels in the ZBTB10 KO BeWo cells (Fig. 3B). Loss of ZBTB10 did not affect cell growth or morphology in undifferentiated BeWo cells (Fig. S5A,B), pointing to a role in differentiation rather than self-renewal. Accordingly, upon Forskolin-induced differentiation, ZBTB10 KO cells secreted significantly less hCG, by both western blot and ELISA assays, and underwent substantially less cell-cell fusion in two-color assays (Fig. 3B-D), reproducing the screen phenotypes. To define the transcriptional basis of these defects, we performed RNA-Sequencing on Forskolin-treated wild-type (WT) versus ZBTB10 KO BeWo cells, revealing ∼2,500 up-regulated and ∼2,500 down-regulated transcripts altered upon differentiation in KO cells (Fig. 3E). Consistent with the loss of hCG secretion at the protein-level, the transcripts for *CGB3*, *CGB5*, and *CGB8* were significantly downregulated compared to WT levels. Strikingly, however, most canonical regulators of trophoblast fusion including *ERVFRD-1*, *GCM1, OVOL1,* and *TFEB,* were unperturbed (Fig. 3E), indicating that ZBTB10 acts largely outside the established GCM1–syncytin axis. Instead, one of the most dysregulated genes was *MFSD2A*, the receptor for Syncytin-2 mediated fusion. Like CGB cluster genes, MFSD2A is typically up-regulated during forskolin induced differentiation but failed to be robustly up-regulated upon ZBTB10 loss (Fig. 3F,G), suggesting that ZBTB10 may be required to transcriptionally activate the receptor in order to mediate fusion. Furthermore, progenitor genes typically repressed during BeWo differentiation, such as *SLC1A5* and *CDH1,* were aberrantly retained (Fig. 3E), and genes that fail to be induced were enriched for fusion-relevant terms such as cell-cell adhesion and cell junction assembly (Fig. S5C), together suggesting ZBTB10 both activates differentiation genes and represses the stem-cell program. Finally, we validated an additional fusion hit MED13 from our primary screen through biological replicate knockouts and observed its loss consistently abolishes cell-cell fusion without perturbing BeWo growth (Fig. S5A-B,D), further supporting the screen’s ability to identify novel regulators of trophoblast differentiation.

**Figure 3.**
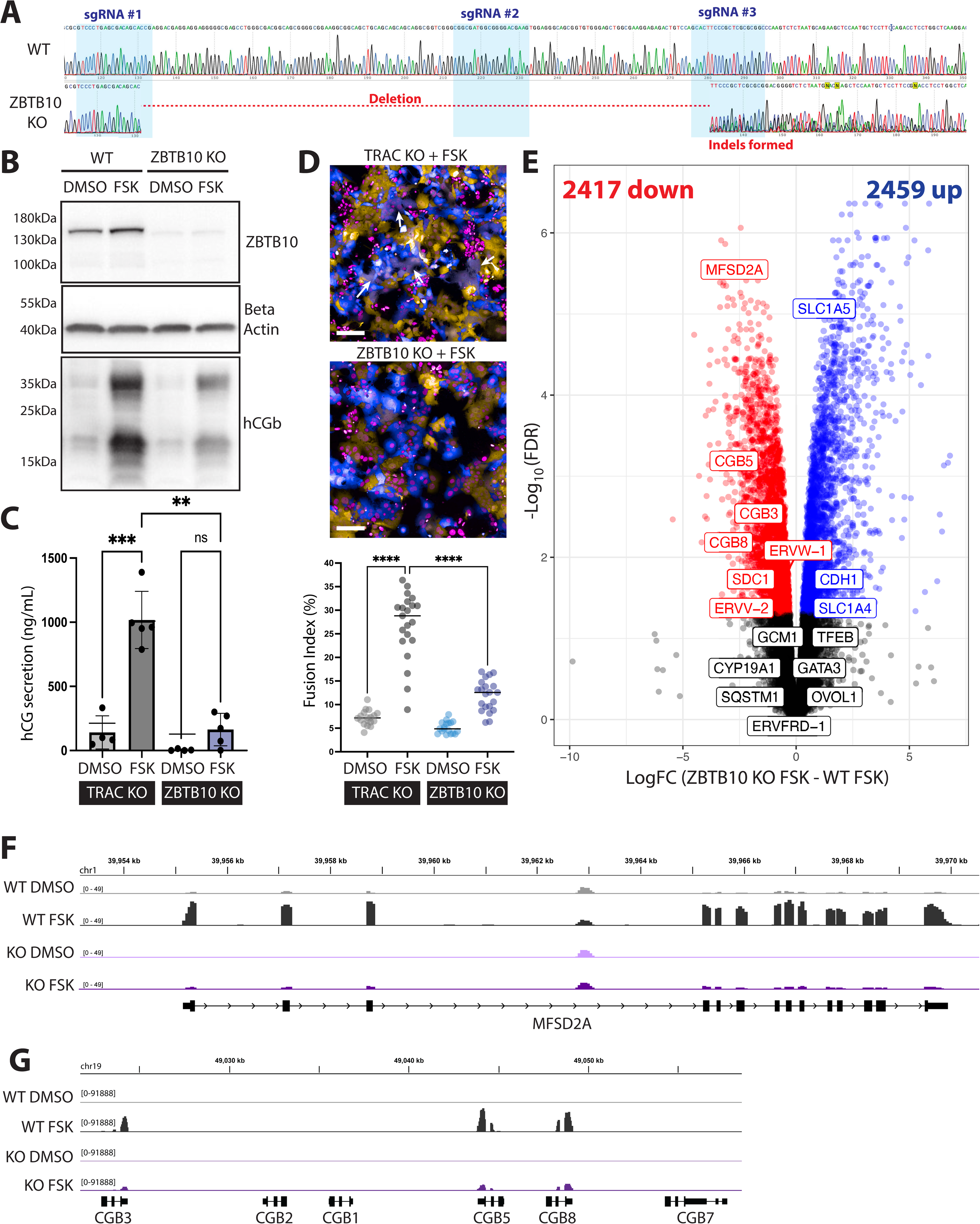
ZBTB10 controls BeWo fusion via expression of MFSD2A, the Syncytin-2 receptor, and hormone cluster genes. **A)** Sanger Sequencing of polyclonal ZBTB10 knockout BeWo cells with deletion and indel formation. **B)** Western blot of ZBTB10 in wild-type (WT) and ZBTB10 KO BeWo cells treated with DMSO or Forskolin for 48 hours (differentiated). **C)** hCG secretion measured by ELISA in supernatant from WT vs. ZBTB10 KO BeWos treated with DMSO or Forskolin for 48 hours (differentiated). Significance bars indicate Brown-Forsythe and Welch ANOVA test with adjusted p-value less than 0.0332 (*), 0.0021 (**), and 0.0002 (***). **D)** Example images of BeWo fusion assays in polyclonal TRAC KO (control) versus ZBTB10 KO mCherry and GFP BeWo co-cultures. GFP BeWos (blue) and mCherry BeWos (yellow) and Hoechst-labeled nuclei (magenta) are shown with white arrows indicating examples of fused regions. Quantified Fusion Index across individual wells from at least 3 independent biological experiments is shown where comparisons indicate a One-way ANOVA with mixed-effects analysis, Geisser-Greenhouse correction, and Dunnett’s multiple comparisons test and **** = adjusted p-value <0.0001. Scale bars are 200μm. **E)** RNA-Seq differential gene expression analysis comparing Forskolin-differentiated ZBTB10 KO BeWos versus Forskolin-differentiated WT BeWos. Significantly up-regulated genes (FDR<0.05, Log Fold-Change >0) are shown in blue, and significantly down-regulated genes (FDR<0.05, Log Fold-change <0) are shown in red. Unchanged genes are in black. **F, G)** Genome browser tracks for **F)** *MFSD2A* and **G)** the CGB cluster genes in BeWo RNA-seq data. Merged (summed) reads from 3 RNA-Seq replicates are shown.

### ZBTB10 facilitates syncytiotrophoblast fusion and is essential for EVT differentiation

To ask whether ZBTB10 regulates trophoblast differentiation in a more physiological setting than the choriocarcinoma model, we turned to human trophoblast stem cells (hTSCs).^49–51^ To exclude off-target effects of the screen guides, we knocked out ZBTB10 in hTSCs with three new sgRNAs targeting exons 4-6 which encompass the DNA-binding domain and predicted nuclear localization signal of ZBTB10. This polyclonal knockout in hTSCs reduced nuclear ZBTB10 by ∼85% relative to a non-targeting control (NEG; Fig. 4A). Loss of ZBTB10 reduced hTSC proliferation but did not abolish self-renewal or alter the expression of canonical TSC markers (Fig. 4B, Fig. S6A,B), indicating that ZBTB10 is dispensable for the stem state itself.

**Figure 4.**
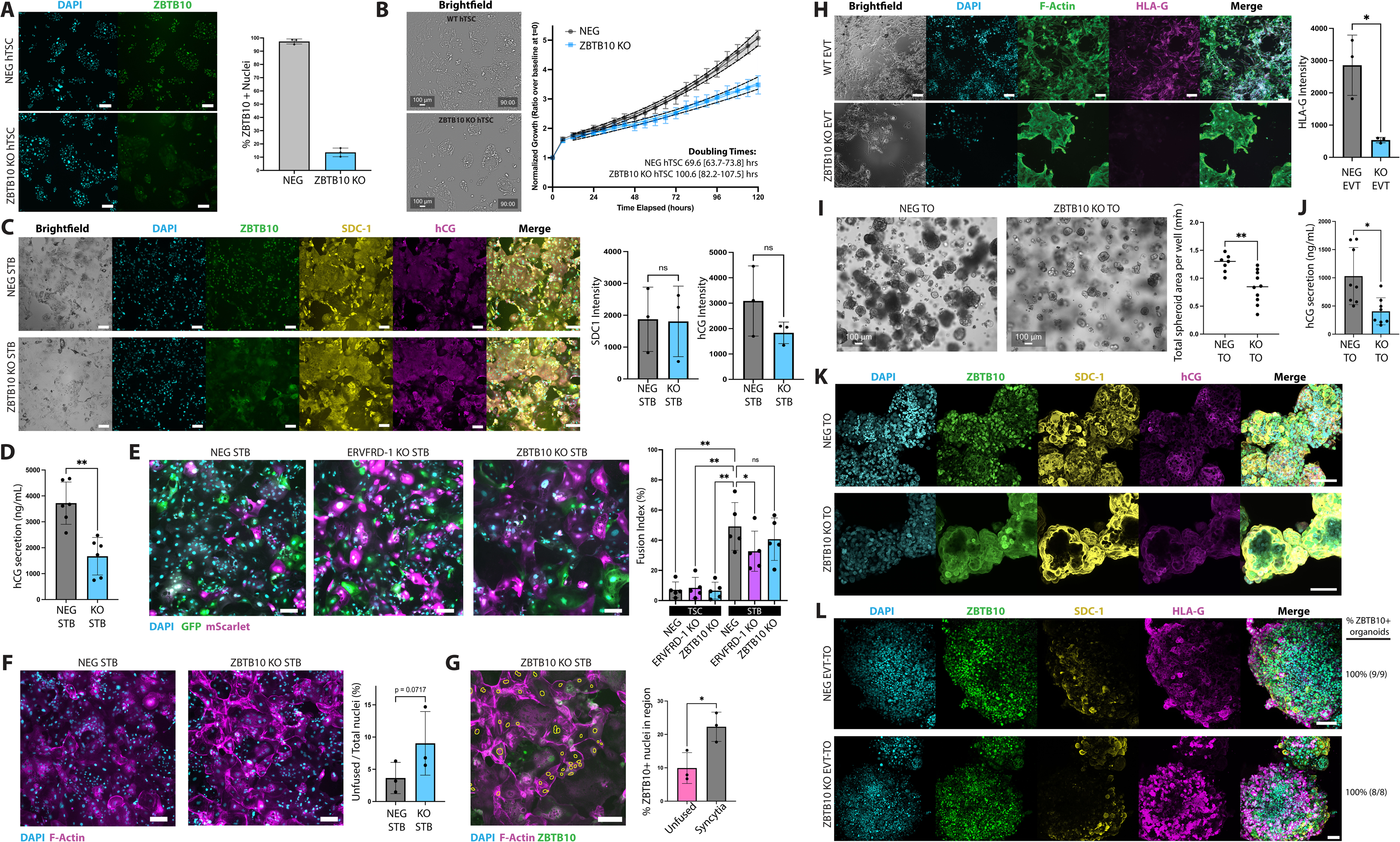
ZBTB10 regulates STB and EVT differentiation in human trophoblast stem cells and organoids. **A)** Immunofluorescence staining of ZBTB10 in negative control (NEG) and polyclonal ZBTB10 knockout hTSCs and quantified percent of nuclei with nuclear ZBTB10 expression shown at right. **B)** Cell growth curves for NEG and ZBTB10 KO hTSCs (6-8 wells per experiment in 3 independent biological experiments). Confluency is normalized to t=0 per well and the 95% confidence interval is shown as dashed lines. Brightfield images of cell morphology and confluency at 90hrs of growth shown at right. **C)** Immunofluorescence of NEG and ZBTB10 KO hTSCs differentiated for 6 days to STB. Quantified immunofluorescent intensity for 3 independent biological replicates and compared by paired t-test. **D)** hCG secretion measured by ELISA in supernatant from NEG vs. ZBTB10 KO differentiated STB compared by Welch’s t-test from six technical replicates across three biological replicates. **E)** Raw images of 2-color fusion assays performed in GFP/mScarlet co-cultured hTSCs treated with 0.1% DMSO (undifferentiated) or differentiated for 6 days into STB. Quantified Fusion Index for five biological replicates shown at right. The statistical comparison indicates a RM one-way ANOVA test with the Geisser-Greenhouse correction and Dunnett’s multiple comparisons test. **F)** F-actin (Phalloidin; magenta) and DAPI (blue) staining of 6-day STB differentiated NEG and ZBTB10 KO hTSCs. The quantified percentage of annotated unfused cells over total nuclei in each condition is shown at right and compared by paired t-test. **G)** Close-up overlay of Phalloidin (magenta) and ZBTB10 (green) immunofluorescence in 6-day STB differentiated ZBTB10 KO STBs. Nuclei within annotated unfused cells are shown with yellow outlines. The percent of ZBTB10+ nuclei in unfused versus syncytialized regions is quantified at right for 3 independent biological replicates compared by paired t-test. **H)** Immunofluorescence of 6-day EVT differentiated NEG control versus ZBTB10 KO hTSCs. Quantified intensity of HLA-G staining is shown at right for 3 independent biological replicates and compared by paired t-test. Scale bars are 200μm in panels A-H. **I)** Brightfield images of Day 7 hTSC trophoblast organoids (TO) in Matrigel domes. The total area occupied by organoids within the imaged domes is quantified at right and compared via Welch’s t-test. **J)** hCG ELISA measurements for supernatant sampled from Day 7 trophoblast organoids formed from NEG versus ZBTB10 KO hTSCs and compared via Welch’s t-test. **K)** Immunofluorescent staining in cleared trophoblast organoids (TOs) formed from NEG versus ZBTB10 KO hTSCs. Each image is a background-subtracted, maximum intensity projection of Z-stack confocal images taken through the organoid. **L)** Immunofluorescent staining of NEG versus ZBTB10 KO EVT-TOs with the percent of organoids demonstrating nuclear ZBTB10 signal quantified at right. Image is a background-subtracted, maximum intensity projection of Z-stack confocal images. Scale bars are 100μm in panel K-L. For all figures, error bars indicate standard deviation and adjusted p-values are indicated by asterisks as less than 0.0332 (*), 0.0021 (**), and 0.0002 (***).

We then examined ZBTB10’s function on formation of the two major trophoblast lineages. Upon 6 days of directed differentiation, ZBTB10 KO hTSCs still formed syncytiotrophoblast (STB) and expressed STB markers SDC1 and hCGβ comparably to control (Fig. 4C) yet produced less flux of hCG secretion as measured by ELISA (Fig. 4D), suggesting ZBTB10’s role is in supporting the secretory output of STB rather than its specification. Cell–cell fusion, by contrast, was only subtly affected by ZBTB10 loss; applying the two-color cell-cell fusion assay system to hTSCs, ZBTB10 knockout produced a slight, but non-significant trend toward lower fusion (Fig. 4E). We did, however, observe altered cellular and cytoskeletal morphology and quantified a small, but notable increase in the proportion of completely unfused cells in the ZBTB10 KO STBs (Fig. 4F). Interestingly, with knockout heterogeneity in the population, ZBTB10⁺ nuclei were significantly enriched in syncytia relative to unfused cells after 6 days of differentiation (Fig. 4G), indicating that ZBTB10-expressing cells are preferentially incorporated into the syncytium. Notably, even knockout of the fusogen ERVFRD-1 itself, although showing a significant decrease in fusion at 6 days, did not reduce it to the near undifferentiated levels seen in BeWo cells (Fig. 4E, Fig. 1D-E), indicating that hTSC fusion is less dependent on any single factor than in BeWo.

In striking contrast to its supporting role in syncytiotrophoblast differentiation, ZBTB10 was absolutely required for the extravillous lineage: after 6 days of EVT differentiation, ZBTB10 KO hTSCs showed dramatically impaired differentiation, with reduced invasive, spiky morphology and near-absent EVT marker HLA-G compared to control (Fig. 4H). These effects were recapitulated in more physiologically relevant 3D trophoblast organoids. ZBTB10 loss reduced organoid growth (Fig. 4I) and while ZBTB10 KO organoids spontaneously differentiated towards STB cells expressing SDC-1 and hCG similar to controls, they had lower overall hCG secretion (Fig. 4J,K). In contrast to negative control hTSCs, directed differentiation of ZBTB10 KO organoids towards the EVT lineage (EVT-TOs) failed to generate KO EVT-TOs, as the KO cells were completely eliminated during differentiation despite starting as the majority of the polyclonal population; all surviving attached EVT-TOs were uniformly ZBTB10⁺, and no ZBTB10-knockout organoids were recovered (Fig. 4L). Because EVT-TO attachment and outgrowth require successful EVT specification, this reinforces that ZBTB10 loss severely impairs commitment to or maintenance of the extravillous lineage.

To define the transcriptional basis of these lineage-specific phenotypes, we profiled control and ZBTB10 KO hTSCs by RNA-sequencing in the undifferentiated TSC, STB, and EVT differentiated states (Fig. S7A). ZBTB10 loss had minimal effects in undifferentiated cells with only ∼400 genes differentially expressed in either direction (Fig. S7B,C) but caused extensive changes upon differentiation, most strongly in EVTs (Fig. 5A,B). In agreement with BeWo data, *MFSD2A* was significantly reduced in ZBTB10 KO STB compared to control, while canonical fusion regulators (*ERVFRD-1, GCM1, TFEB, OVOL1,* etc.) were unchanged (Fig. 5A, Fig. S7F). In EVTs, ZBTB10 loss strongly reduced *HLA-G NOTUM, MMP2, MMP15,* and *ADAM19* expression, key markers of EVT fate and invasion, and overactivated genes associated with the EVT interferon response^52,53^ (Fig. 5B), suggesting a strong transcriptional dysregulation of EVT differentiation. Across both lineages, TSC markers such as KRT7 and GATA3 failed to be as effectively repressed upon differentiation in the ZBTB10 KO cells, suggesting impaired silencing of the progenitor program and a potential repressive role for ZBTB10 during trophoblast differentiation (Fig. 5C, Fig. S7E). GO enrichment analysis was concordant with these phenotypes: ZBTB10 KO STBs aberrantly retained actin filament organization and cell-substrate adhesion while downregulating protein folding and glycolytic pathways, suggesting impaired structural and metabolic maturation; ZBTB10 KO EVTs had elevated viral response and interferon signaling, suggesting incomplete acquisition of the immune-tolerant state required for normal function (Fig. S5D). Together, these data establish ZBTB10 as a transcriptional regulator of both human trophoblast lineages, with roles in activating key EVT genes, supporting STB function, and repressing the trophoblast stem-cell state during differentiation.

**Figure 5.**
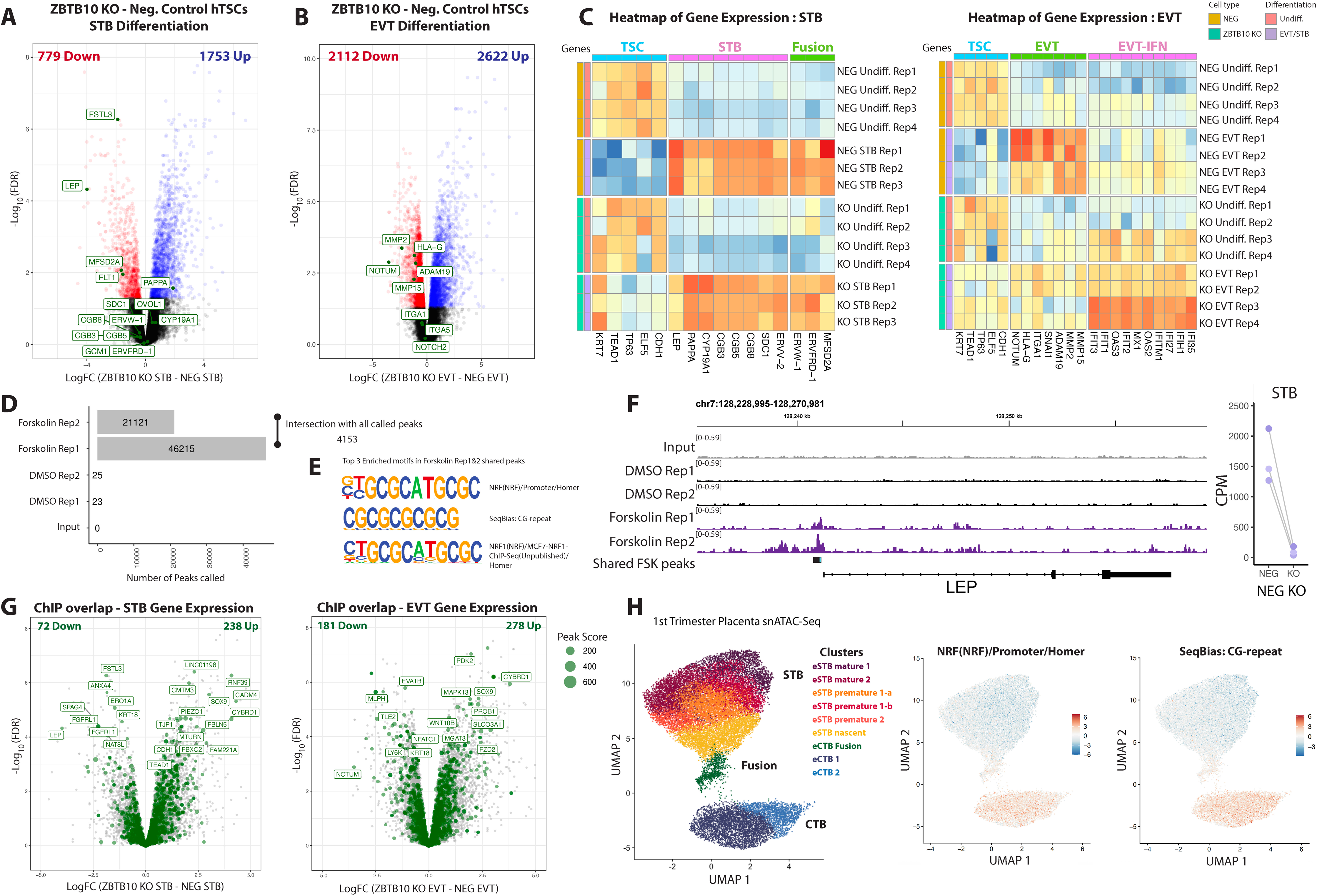
ZBTBlO binds GC-rich motifs to control STB, EVT, and TSC genes. **A, B)** RNA-Seq differential gene expression analysis comparing **A)** day 6 STB-differentiated ZBTB10 KO versus NEG hTSCs or **B)** day 6 EVT-differentiated ZBTBl0 KO versus NEG hTSCs. Significantly up-regulated genes (FDR<0.05, Log Fold-Change >0) are shown in blue, and significantly down-regulated genes (FDR<0.05, Log Fold-change <0) are shown in red. Unchanged genes are in black. **C)** Heatmap of column-scaled log2(CPM) values from individual RNA-Seq replicates, grouped by marker genes for TSC, STB, EVT, and Fusion-related genes, normalized per gene column. **D)** Total number of called MACS2 peaks per replicate and condition for BeWo ZBTBlO ChIP. The number of peaks called in both Forskolin replicates is shown by the “intersection” at right. **E)** Top 3 sequence motifs from HOMER motif emichment of the Forskolin-treated consensus ZBTB10 ChlP peaks. **F)** Genome browser track of ZBTB10 ChlP in BeWo cells at the *LEP* gene. Consensus ZBTBlO peaks identified in both Forskolin replicates are shown with a black bar and the peak center indicated by a teal bar overlay. **G)** Volcano plot showing the overlay of ZBTB10 target genes identified by ChIP (consensus peak within+/- 2000bp of promoter) in BeWo (green) and differentially expressed genes in STB and EVT differentiated hTSCs, as in Figure 5A,B. The number of unique gene targets (by ChlP) that are significantly dysregulated upon ZBTB10 KO in each condition are indicated at the top of the plot. **H)** Annotated cell clusters from snATAC-Seq of first trimester human placenta (Wang, et al. 2024^59^_)_ are plotted by UMAP (left) and major cell groups are indicated (CTB, STB, and Fusion). Right, per-cell ATAC activity (chromVAR deviation z-scores) for the specified ZBTBlO ChlP motifs are overlaid on the snATAC-Seq UMAP.

### ZBTB10 binds GC-rich motifs to control placental gene expression

To identify the genomic targets of ZBTB10 during trophoblast differentiation, we mapped ZBTB10 binding genome-wide by performed ChIP sequencing in BeWo cells stably overexpressing Halo-3xFLAG-ZBTB10 in both undifferentiated (DMSO) and STB differentiated (Forskolin) states. Enrichment of ZBTB10 binding to chromatin was significant over input controls and called peaks localized near transcription start sites in the Forskolin-differentiated samples, but ZBTB10 binding was close to zero in the undifferentiated samples (Fig. 5E, Fig. S8A,B). This suggests ZBTB10 is preferentially recruited to chromatin as cells differentiate. Given the modest signal-to-noise ratio of the ZBTB10 ChIP due to the moderate transcription factor expression and the lack of a ChIP-seq-validated antibody, we adopted a stringent, reproducibility-based definition of binding, focusing on the 4,153 peaks that were shared across Forskolin replicates as high confidence ZBTB10 genomic targets (Fig. 5E). Motif enrichment analysis within consensus ZBTB10 peaks identified GC-rich NRF and CG-repeat motifs as likely ZBTB10 binding sites (Fig. 5F), consistent with prior reports that ZBTB10 binds GC-rich regions in competition with Sp1.^54^ Notably, several of the ZBTB10 ChIP targets connect directly to the knockout phenotypes. A distinct ZBTB10 peak marked the promoter of the *LEP* gene, as well as the gene body of *FSTL3*, two STB-secreted hormones that regulate trophoblast growth, invasion, and metabolism^55–57^ and both *LEP* and *FSTL3* were strongly downregulated in ZBTB10 KO STBs (Fig. 5B, 5G, S8C), nominating *LEP* and *FSTL3* as ZBTB10-activated genes. Slight enrichment of ZBTB10 at the MFSD2A promoter could be seen but was not significant due to low signal (Fig. S8C). *NOTUM*, a WNT pathway modulator required for EVT differentiation, was also downregulated and identified as a target of ZBTB10, suggesting a potential direct link of ZBTB10 to the EVT phenotype (Fig. S8C).^58^ We also detected ZBTB10 binding at the *CDH1* and *TEAD1* promoters, TSC transcripts that are up upregulated upon ZBTB10 loss, indicating that ZBTB10 also acts to repress the progenitor state during differentiation (Fig. S8C). Intersecting the ZBTB10 ChIP-Seq peaks with the differentially expressed transcripts in the KO EVT and STBs indicates that ZBTB10 binds both up- and down-regulated genes, confirming at the genomic level that it acts as both a repressor and activator of gene expression programs in trophoblasts, with a slight bias towards repression (more genes up-regulated upon knockout were ZBTB10-bound; Fig. 5H).

Finally, to infer where ZBTB10 likely acts in vivo across trophoblast states, we examined chromatin accessibility at the ZBTB10 motif (NRF and CG-repeat) in first trimester placental snATAC-seq data.^59^ ZBTB10 motif accessibility is specifically enriched in the cytotrophoblast (CTB) and fusing CTB populations, the cell states poised for the CTB-to-STB transition, reiterating that ZBTB10 likely acts at this stage to both repress stem cell genes and activate differentiation genes, such as *LEP* (Fig. 5I). Moreover, the pattern of ZBTB10 motif accessibility in the CTB and fusing CTB clusters was unique (other early gestation STB-associated TFs such as MITF and CEBPA showed different patterns) and was not due to global changes in chromatin accessibility or read depth within those clusters (Fig. S8D-F), suggesting a specific role for ZBTB10 in facilitating accessibility during the CTB to STB transition. Together, these data suggest ZBTB10 functions as a dual activator/repressor to binds and regulate genes during placental differentiation.

### ZBTB10 is broadly expressed in the placenta and enriched in fusing CTBs and EVTs

Given the functional importance of ZBTB10 in *in vitro* models of placental differentiation, we next sought to investigate the expression of ZBTB10 within human placental tissue across gestation. In snRNA-Seq data from first trimester placentas^59^, *ZBTB10* is most highly expressed in the fusing CTB population, which shows both the highest mean expression and percent of cells expressing *ZBTB10* (Fig. 6A). Substantial expression is also found in a subset of EVT cells with the highest HLA-G expression (Fig. 6A, Fig. S9A). These data match the cell states in which ZBTB10 is functionally required. Single-nucleus multi-omic profiling of the human placenta has identified that STBs follow a bifurcated maturation trajectory, in which a common early population (eSTB nascent) gives rise to two terminal types of nuclei within the syncytium: a PAPPA⁺ growth-hormone type (eSTB mature 1) and an FLT1⁺ transport type oriented toward oxygen response and lipid transport (eSTB mature 2), with both states present in roughly equal proportions during early pregnancy but shifting towards predominantly PAPPA+ STBs at term.^59^ Across all STB subtypes, *ZBTB10* expression declined modestly from early to late gestation and no specific enrichment of *ZBTB10* was identified across cell types from term placentas, suggesting that ZBTB10 plays a more prominent role in early placental development (Fig. S9C). Mapping ZBTB10 expression along the pseudotime trajectory of STB maturation in first trimester placentas revealed enrichment in the early CTB fusion state along with high *ERVFRD-*1 expression, consistent with its supporting role in fusion, but also in the eSTB mature 2 (late pseudotime) arm, which highly expresses *FLT1*, suggesting an additional role in maturation of the FLT1+ STB subtype (Fig. S9D). Gene set enrichment analysis showed that ZBTB10 KO in STB caused a significant and preferential loss of the FLT1+ mature 2 gene signature and a slight enrichment of the PAPPA+ mature 1 gene signature, suggesting ZBTB10 may control the balance or maturation of these transcriptional subtypes within the early STB (Fig. 6B). At the protein level, ZBTB10 was moderately expressed in hTSCs with slightly lower expression in fully differentiated STBs but was most highly expressed in EVTs (Fig. 6C,F). Consistently, in first-trimester placental tissue, immunostaining revealed distinct nuclear ZBTB10 protein throughout the chorionic villus including in CTB and STB nuclei, with particularly strong expression in invading HLA-G+ EVTs (Fig. 6D,F,G). At term, broad ZBTB10 expression persisted in a subset of CTB and STB nuclei (Fig. 6E,G). Our data suggests that ZBTB10 is a broadly expressed placental transcription factor with enriched expression particularly in fusing CTBs and invasive EVTs during early placental development, the populations in which we found it functionally required, consistent with its roles in STB and EVT differentiation.

**Figure 6.**
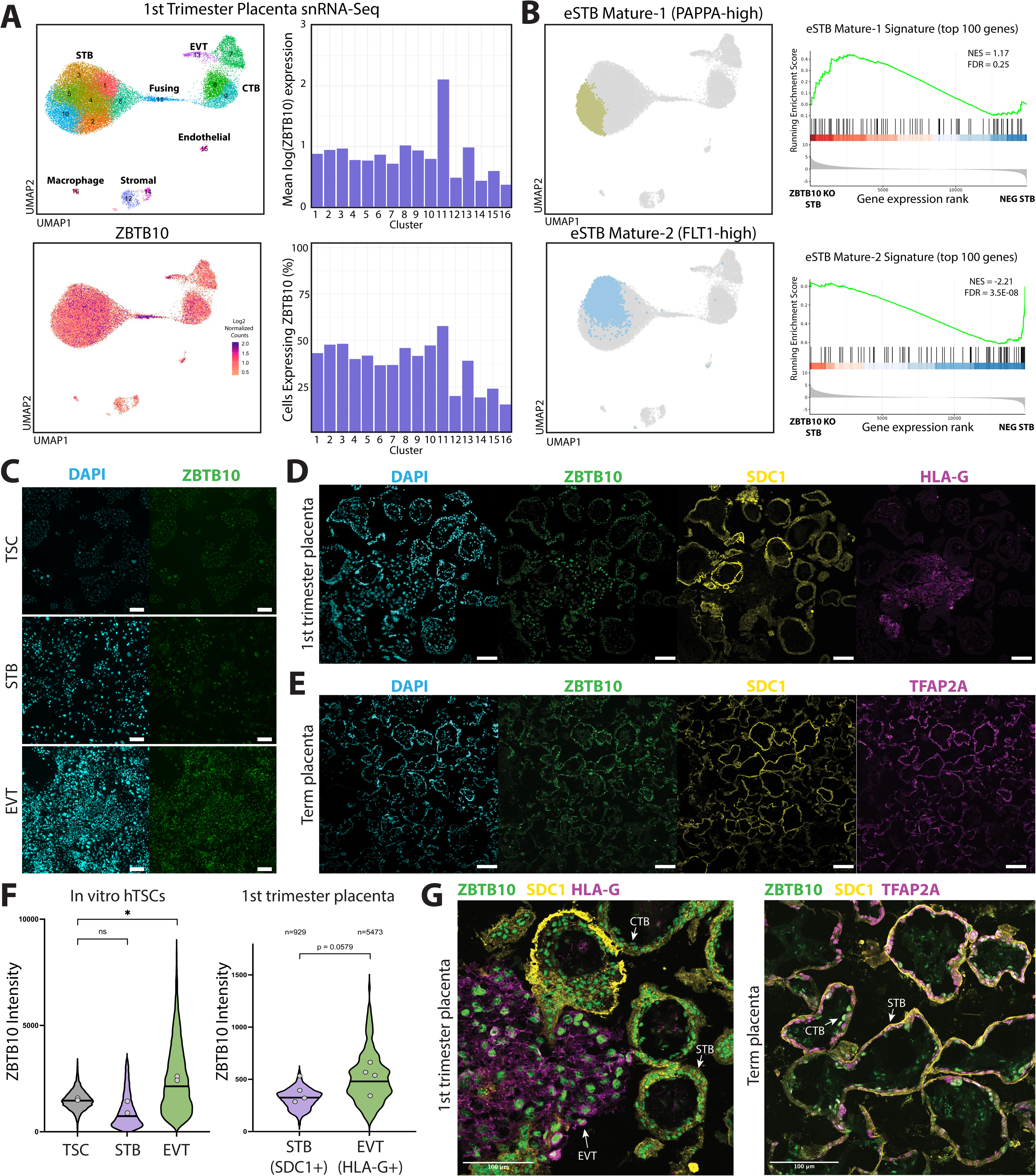
ZBTB10 is expressed in the human placenta and enriched in fusing CTBs and EVTs. **A)** Annotated cell clusters from snRNA-Seq of first trimester human placenta (Wang, et al. 2024^59^) are plotted by UMAP with cluster number indicated (top). Cell groups are labeled by major categories (CTB, STB, and Fusing, etc.). Bottom, ZBTB10 expression is shown on the same UMAP with log2-normalized counts per cell. Right, Bar charts quantify the mean expression and the percent of cells with >0 reads for ZBTB10 in each annotated cell cluster. **B)** The two mature STB clusters overlaid on the UMAP plot of first trimester placenta snRNA-Seq. Right, GSEA plots of the top 100 differentially expressed genes in eSTB mature 1 and mature 2 pathways, with genes ordered from enrichment in ZBTB10 KO STBs (left) to NEG STBs (right). Black bars indicate the presence of each pathway gene in the ordered gene set. NES = normalized enrichment score. **C)** Immunofluorescence of ZBTB10 in undifferentiated hTSCs versus day 6 EVT-differentiated or STB-differentiated hTSCs. Scale bars are 200μm in panel C. **D,E)** Immunofluorescence of **D)** first trimester (weeks 8-12) and **E)** term (weeks 36-40) human placental tissue. **F)** Left, quantification of ZBTB10 intensity in immunofluorescence of in vitro differentiated hTSCs. Comparison indicates a RM one-way ANOVA test with the Geisser-Greenhouse correction and Dunnett’s multiple comparisons test where * = adjusted p-value < 0.0332. Right, quantification of ZBTB10 intensity quantified in the indicated segmented marker-positive cells within first trimester placental tissue. Statistical comparison indicates a paired t-test comparing means between individual donors (n=4). **G)** Close-up overlay images of immunofluorescence staining shown in D,E shown at bottom right with example cell types indicated by white arrows. Scale bars are 100μm in panels D-G.

### Conservation of ZBTB10 and evolution of a placental-specific function

ZBTB10 belongs to the BTB-zinc-finger family (which includes BCL6), whose members are conserved across vertebrates and play important roles in non-placental tissues, notably in hematopoiesis and immunity.^60–63^ In humans, ZBTB10 is broadly expressed across tissues, with the highest expression in the bone marrow and liver, and is highly enriched in the placenta, ranking as the third most highly expressed tissue (The Human Protein Atlas). To elucidate how tissue-specific expression of ZBTB10 might have evolved, we performed amino acid alignment of ZBTB10 and found that while the overall protein sequence diverges with evolutionary time, the two Zn-finger DNA binding domains of ZBTB10 are almost perfectly conserved from humans to fish consistent with an ancestral role in non-placental tissues (Fig. S10A-C). To further investigate recent evolutionary shifts in mammals, we examined six single-nucleus and single-cell placenta sequencing datasets from diverse mammalian species including human (*Homo sapiens*), macaque (*Macaca fascicularis*), mouse (*Mus musculus*), guinea pig (*Cavia porcellus*), the marsupial grey short-tailed opossum (*Monodelphis domestica*), and the Malagasy common tenrec (*Tenrec ecaudatus*).^64^ UMAP analysis confirmed conserved cell populations exist across all species, including trophoblasts, lymphoid, and myeloid cell types among others (Fig. S10E,F). In all five eutherian species, ZBTB10 expression is much higher in trophoblast cell types than in lymphoid or myeloid cell types (Fig. S10D,G), whereas in the marsupial opossum, which has limited placental invasion, ZBTB10 was instead highest in dendritic cells (Fig. S10D), consistent with its ancestral immune role. In the tenrec, the most distantly related eutherian species with an invasive placenta, *ZBTB10* is highly expressed in both hematopoietic lineages such as dendritic cells and B cells as well as in extraplacental trophoblasts (EPTs), the functional equivalent of EVTs (Fig. S10D). Given that tenrec EPTs and human EVTs share expression of the invasion-related genes *ADAM19* and *MMP15*, both of which are downregulated in human ZBTB10 KO EVTs, these data suggest a conserved role for ZBTB10 in regulating extravillous trophoblast invasion across eutherians (Fig. 5B).^64^ Collectively, the enrichment of ZBTB10 in trophoblasts across tenrec, guinea pig, mouse, macaque, and human supports a hypothesis that placenta-specific expression of ZBTB10 was acquired alongside invasive placentation during eutherian evolution (Fig. S10D).

## DISCUSSION

By simultaneously measuring live-cell fusion, hCG secretion, and detailed cellular morphologies in the same wells, our scalable arrayed CRISPR screen identified 117 regulators of trophoblast differentiation. Our unique approach overcomes previous limitations to create a genetic screen compatible with the unique biology of fusion: arrayed screening avoided clonal isolation which is impractical across hundreds of genes, the short differentiation in BeWo cells minimized compensatory adaptations that can obscure loss-of-function phenotypes in longer-term differentiation assays, and paired assays captured both gain and loss-of-function effects across multiple phenotypes. The screen uncovered over a hundred genes with impacts on STB differentiation, recovering established regulators like GCM1 and STK40 which validated its robustness, and nominating three previously uncharacterized regulators as strong effectors of STB differentiation: ZBTB10, EGLN1, and ZNF292.

The joint profiling of fusion and hCG shows the two hallmark functions of the STB are largely separable rather than obligately coupled. Previous studies have hinted at this separability: for example, direct disruption of Syncytin-1 or Syncytin-2 impairs fusion without affecting hCG secretion^19^, while knockdown of hCG signaling alters fusion in some studies but not others.^26,27^ Moreover, regulators such as leukemia inducing factor (LIF) enhances cell-cell fusion efficiency while decreasing hCG, and MAPK14 inhibitors decrease hCG secretion without changing fusion.^28,29^ Our screen provides a phenotype-based quantitative resource for the field that reveals that hCG, the primary serum marker of placental function in vivo, can be regulated independently of syncytial integrity. This dissociation may explain inconsistent performance of hCG as a preeclampsia predictor despite evidence of syncytial dysfunction in preeclampsia and suggests that more selective factors could help discriminate and potentially correct different causes of syncytial dysfunction during pregnancy.^6,7,9,65^

Among the strongest hits, EGLN1 (PHD2), a HIF regulator downstream of GCM1, whose loss causes placental malformation in mice^36,66,67^, and ZNF292, a factor implicated in STB gene regulatory networks^68^, both carry prior placental links but have not yet been functionally tested in STB differentiation. ZBTB10, by contrast has no described placental role so far so we sought to uncover its importance in trophoblast biology. Functional studies in physiologically relevant hTSC systems revealed a broad role for ZBTB10 in placental development, concordant with its expression in human placental tissue. We find that ZBTB10 coordinates multiple trophoblast fates by binding to specific genes to promote EVT and STB function (e.g. *LEP, NOTUM)* as well as to restrain epithelial and progenitor programs (e.g. *CDH1*, *TEAD1*), thereby acting as a dual activator and repressor to dictate trophoblast lineage decisions.

Intriguingly, ZBTB10 loss robustly decreased BeWo fusion and hormone secretion, yet did not perturb the canonical GCM1-syncytin axis, suggesting that ZBTB10 operates through a distinct, parallel program in STB establishment. In hTSC-derived STBs, ZBTB10-KO cells still formed STB but had lower transcript levels of *MFSD2A* and reduced hCG secretion. However, unlike in BeWo, fusion was only partially impaired rather than abolished. In combination with the observation that individual ZBTB10- cells were preferentially unfused suggests that ZBTB10 modulates fusion from the receiving cell by controlling expression of the Syncytin-2 receptor MFSD2A and that the distinct requirement is masked by the existence of unedited cells within the polyclonal KO hTSC population. Furthermore, we found that ZBTB10 directly regulates transcription of two key placental hormones secreted by the STB – leptin (LEP) and the follistatin homolog FSTL3. Pathological elevation of both LEP and FSTL3 have been noted in preeclampsia and fetal growth restriction, and elevated circulating FSTL3 protein is predictive of subsequent PE and FGR, which suggests ZBTB10 could be evaluated as a way to control these targets in pregnancy diseases.^69,70^ More broadly, by comparing our transcriptomics to the two major mature STB states in vivo, we find that ZBTB10 loss biases away from the hypoxia-responsive transport program (FLT1⁺) and towards the growth-hormone-responsive STB state (PAPPA⁺) that is dominant at term. Together, these data indicate that ZBTB10 loss reduces the fusogenic and secretory properties of the STB without losing the lineage entirely but specifically compromises the early FLT1+ transport-type population, in line with the decline of both ZBTB10 and the FLT1+ state across gestation.^59^ This suggests ZBTB10 coordinates multiple important functions of the STB, including fusion and subsequent endocrine maturation, and may govern the clinically relevant balance between STB subtypes in the placenta.

Surprisingly, we find that ZBTB10 is strictly required for formation of a functional EVT lineage. Loss of ZBTB10 impairs functional EVT differentiation, characterized by reduced expression of invasion-associated genes, loss of invasive morphology, and activation of interferon and antiviral response pathways. These results are consistent with recent studies showing that interferon exposure itself abrogates EVT invasion^71,72^, and suggests that ZBTB10 KO causes a failure to establish the transcriptional and functional state required for trophoblast invasion at the maternal–fetal interface. The complete elimination of ZBTB10-KO cells during EVT-TO differentiation further underscores this requirement, identifying ZBTB10 as a potential master regulator of the placental invasion program. The invasive role of ZBTB10 may have deep evolutionary origins. We find ZBTB10 enriched in trophoblasts across eutherians, and in the distantly related tenrec it marks invasive extraplacental trophoblasts that co-express ADAM19 and MMP15, the same invasion genes ZBTB10 regulates in human EVT. Cooption of ZBTB10 expression in the placenta may thus have been an important step in the evolution of invasive placentation. This finding of cooption mirrors the case of GCM1, a master transcriptional regulator of glial cell fate in *Drosophila*, which became evolutionarily placentally-enriched and essential for trophoblast lineage differentiation, while maintaining its ability to regulate glial cell fate in mammals.^39,73–76^ Whether ZBTB10’s ancestral immune role also remains relevant in the placenta and could relate to the interferon program de-repressed in ZBTB10-KO EVTs remains open and worth exploring.

In summary, we identify ZBTB10 as a conserved, cross-lineage regulator of human trophoblast differentiation, required for EVT invasion and STB maturation through coordinated activation and repression of distinct trophoblast gene programs. This work provides a systematic resource and regulatory framework for dissecting how the major fusogenic and secretory functions of the placenta are established and how genetic dysregulation may contribute to placental pathogenesis.

## MATERIALS AND METHODS

### Ethics Statement

The study protocol was approved by the Institutional Review Board at the University of Washington under reference number STUDY00023408. This study involves the use of previously collected de-identified samples from the Washington Pregnancy Biorepository (STUDY00010803) and the Birth Defects Research Laboratory (STUDY00000380). Study subjects provided written informed consent for the use of tissue for research purposes. This study was also reviewed and approved by the Embryonic Stem Cell Research Oversight (ESCRO) Committee at the University of Washington.

### Cell culture

#### BeWo culture

Human placental choriocarcinoma BeWo cells (ATCC CCL-98) were cultured at 37°C and 5% CO2 in F-12K medium (Corning or ATCC) supplemented with 10% fetal bovine serum and 10 U/mL penicillin–streptomycin and 5ug/mL Cefotaxime (Millipore Sigma, C7912-1G) antibiotics. Cells were passaged by trypsinization every 2–4 days and split at a ratio of 1:3–1:5. Media was changed completely every 24–36 h.

#### Naïve-derived human trophoblast stem cells (hTSCs)

Human trophoblast stem cells were established previously from naïve human pluripotent stem cells (RUES2 NIHhESC-09-0013) and cultured in hypoxic conditions (37°C, 5% CO_2_, 5% O_2_) in TSC Media (DMEM/F12 supplemented with 0.1 mM β-mercaptoethanol, 0.2% FBS (Sigma, F4135), 0.5% Penicillin-Streptomycin (P/S; Gibco, 15140122), 0.3% BSA, 1% ITS-X (Gibco, 41400045), 1.5μg/mL L-ascorbic acid, 50ng/mL EGF, 2μM CHIR99021, 0.5μM A83-01, 1μM SB431542, 0.8mM VPA, and 5μM Y-27632).^51^ Media was changed every other day and TSCs were passaged using TrypLE Express (Gibco, 12605010) when 80% confluent (approximately every 4 days) at a ratio of 1:4–6 on Cultrex (R&D)-coated plates. Cells from passage 10-25 were used.

#### Differentiation of hTSCs

TSCs were grown until 80% confluent, then dissociated with TrypLE Express for 8–15 min at 37 °C. For the induction of EVTs, TSCs were seeded at a density of 1.05E4 cells/cm^2^ into Ibidi 4-chamber glass-bottom slides (catalog #: 80426) or CellVis glass-bottom 24-well plates (P24-1.5H-N) pre-coated with Laminin-521 (BioLamina catalog #: LN521-05; diluted in PBS, coated overnight at 4C) for imaging or Cultrex-coated 6-well plastic tissue-culture dishes for RNA-Seq. Cells were allowed to adhere overnight in TSC medium, then differentiation was initiated by switching to EVT1 medium (DMEM/F12, 0.1mM β-mercaptoethanol, 0.5% Pen/Strep, 0.3% BSA, 1% ITS-X, 100ng/mL NRG1, 7.5μM A83-01, 2.5μM Y-27632, and 4% Knockout Serum Replacement) supplemented with 1% Geltrex (Gibco) (Day 1). On day 3, the medium was replaced with EVT2 medium (EVT1 medium without NRG1) supplemented with 0.5% Geltrex. On day 5, the medium was replaced with EVT2 medium. Cells were analyzed on day 7. For the induction of STBs, TSCs were seeded identically as above. Cells were allowed to adhere overnight in TSC medium, then differentiation was initiated by switching to STB medium (DMEM/F12, 0.1mM β-mercaptoethanol, 0.5% Pen/Strep, 0.3% BSA, 1% ITS-X, 2.5μM forskolin, 2.5μM Y-27632, and 4% Knockout Serum Replacement) on Day 1. The medium was replaced on day 4, and cells were analyzed on day 7.

#### Trophoblast organoids

TOs were generated following previously established protocols.^49,77,78^ Briefly, TSCs were grown until they were confluent, then dissociated with TrypLE Express and washed twice in DMEM/F12. A total of 6000 cells were suspended in 25μL Matrigel droplets (Corning, 354277), seeded into 48-well plates and incubated at 37 °C for 15 min before adding trophoblast organoid medium (TOM), which consisted of Advanced DMEM/F12, 1x GlutaMAX, 1× N2 supplement, 1× B27 supplement, 1.25mM N-acetyl-L-cysteine, 50 ng/mL recombinant human EGF, 1.5μM CHIR99021, 80ng/mL recombinant human R-spondin-1 (R&D Systems, 4645-RS), 100ng/mL recombinant human FGF-2, 50ng/mL recombinant human HGF, 500nM A83-01, 2.5μM prostaglandin E2 (Sigma, P0409), 0.1mM 2-mercaptoethanol, and 2μM Y-27632. Medium was refreshed every other day. Organoids were maintained for 7 days and then passaged. For passage, Matrigel droplets were broken up in ice cold DMEM/F12 by firm pipetting then centrifuged at 600g for 4 min, the media and gel supernatant removed, and the organoid pellet resuspended in cold DMEM/F12. This process was repeated 3 times to remove gel and resuspend organoids. To promote correct polarity (STB outside), organoids were then transferred to ultra-low adherence Nunclon Sphera 96-well plates and grown in TOM medium for an additional 2 days.^79,80^ Organoids were then harvested at Day 9 by collecting organoids into a 1.5mL tube pre-coated with Anti-Adherence Solution (Stem Cell), washing once with PBS++, then fixing in 4% paraformaldehyde (PFA) overnight at 4C prior to immunostaining.

EVT-TOs were generated following a previous protocol.^49,77,78^ Briefly, TOs were formed in Matrigel for 7 days and cultured in TOM medium, then passaged as above and seeded onto glass-bottom dishes at a 1:2-1:4 ratio in 10uL Matrigel droplets, and cultured in TOM for another 2-4 days or until reaching a size of about 200μm, then switched to TO-EVT1 medium (consisting of Advanced DMEM/F12, 1x GlutaMAX, 0.1mM β-mercaptoethanol, 0.5% P/S, 0.3% BSA, 1% ITS-X, 100 ng/mL NRG1, 7.5μM A83-01, 2.5μM Y-27632, and 4% KSR). After 5-6 days in EVT1 medium, it was replaced with EVT2 medium (TO-EVT1 medium minus NRG1) for an additional 5-6 days. All organoid media were refreshed every 2 days.

### BeWo CRISPR Screening (Berkeley & IGI)

The lyophilized guide RNA library was provided by Synthego and stored at −80C. The guides were resuspended to 100uM in 1x TE by brief vortexing and incubating at 4C for 72 hours. The 100uM library stocks were then aliquoted into sterile 96-well Nunc v-bottom plates using a Benchsmart Pro 96-well pipette, sealed with aluminum seals, and stored at −80C until use. Screen-ready guide RNA stocks were further diluted to 5uM in 1x TE buffer, sealed with aluminum seals, and stored at −80C until use.

sgRNA-Cas9 RNP complexes were prepared immediately prior to nucleofection. Recombinant Cas9 protein (QB3 Macrolab) was diluted to 20μM in 1× TE buffer. For each 96-well nucleofection plate, a master mix consisting of 1,446.4μL Lonza SG buffer supplemented according to the manufacturer’s instructions and 53.6μL 20μM Cas9 was prepared, yielding a total volume of 1500μL. The 96-well library plate containing 6μL of 5μM sgRNA per well was thawed for 10 minutes at room temperature, centrifuged at 500 × g for 2 min to collect condensation, and then maintained on ice. Cas9/Lonza SG master mix (14μL per well) was added directly to each sgRNA-containing well and gently mixed to form RNP complexes. RNPs were incubated on ice until just prior to use, then incubated at room temperature for 10 minutes to promote complex assembly just prior to nucleofection.

BeWo cells were cultured in standard growth medium, expanded in 15cm tissue culture plates, and prepared for nucleofection by routine passaging with trypsinization. For each 96-well nucleofection, 3E7 total cells (1.5E7 GFP BeWo + 1.5E7 mScarlet BeWo) were collected, pooled, and pelleted by centrifugation. Cells were resuspended in 1.5 mL BeWo medium to achieve a final concentration of 1E5 cells per 5μL. Using a multichannel pipette, 5μL cell suspension was added to each well containing preassembled Cas9 RNPs in nucleofection buffer. Immediately following cell addition, 20μL of the cell-RNP mixture was transferred into a 96-well Lonza nucleocuvette plate. Nucleofection was performed using a Lonza 4D-Nucleofector X Unit with the 96-well Shuttle System and program CA-137. Immediately following nucleofection, 80μL prewarmed BeWo medium was gently added to each well without mixing. Cells were allowed to recover for 10 min at 37°C in a humidified incubator with 5% CO₂. Cells were then resuspended by gentle pipetting, and 60μL from each well was transferred into a 96-well tissue culture plate containing 190μL complete medium per well. Media was changed 24 hours post-nucleofection. At 48 hours post-nucleofection, parental plates were passaged using Accutase and quenched in medium and 30% of each parental plate was transferred into 3x receiving Cell Carrier Ultra 96-well plates (PerkinElmer 6055300). Plates were incubated overnight then media was changed to BeWo medium with 20uM Forskolin to induce differentiation in all wells except for DMSO control wells (wells E1-E3 on every plate which received BeWo media + 0.1% DMSO). Cells were differentiated for 48 hours, with respective DMSO and Forskolin media changed at 24 hours of differentiation. At 48 hours, 25uL samples of supernatant were taken from each well and transferred to 96-well PCR plates and stored at −80C until use. Media was then changed to 4.5g/L DMEM without phenol-red containing 10% FBS, 1× GlutaMAX supplement, and 1× sodium pyruvate supplement plus a final concentration of 2μg/mL Hoechst 33342 (Invitrogen) and incubated for 30 minutes before imaging. Plates of live cells were imaged sequentially; each plate took ∼ 1 hour to image so differentiation was staggered so that all plates were imaged within +/-2 hours of the 48-hour differentiation timing.

At nucleofection, an additional 15μL from each 100uL nucleofection reaction was transferred into a separate 96-well tissue culture plate (backup plate) containing 85μL complete medium per well. At ∼72 hours post nucleofection, the confluent backup plate was frozen down (cells were lifted in 96-well plates using 20uL per well Trypsin, then quenched with 150uL per well 10% DMSO + 90% FBS and slowly frozen in a Styrofoam box at −80C) and later used for extraction of gDNA for TIDE analysis of screen editing efficiencies in select wells.

Imaging was performed on a Perkin Elmer Opera Phenix high-throughput spinning disk confocal with 37C, 5% CO2 incubation using a 10X Air objective (NA = 0.3), and the four channels were acquired (Brightfield, Hoechst, GFP, mScarlet), where Hoechst/Brightfield were collected in the same exposure and separated from mScarlet/GFP, which were collected in a second exposure. The following illumination settings and filter sets were used: Brightfield transmission was imaged with transmitted light and 650–760 nm emission filter; GFP was imaged with 488 nm excitation, 500–550 emission; mScarlet was imaged with 561 nm excitation, 570–630 nm emission; and Hoechst was imaged with 375 nm excitation, 435–480 nm emission. Twelve images (fields of view) were acquired per well, which covered almost the entire well area and measured quantities were well-level averaged over all 12 fields.

The hCGβ ELISA (Catalog #: DY9034-05) was performed according to manufacturer’s protocol, using 96-well Immulon 4HBX Flat Bottom Plate (Catalog #: 3855). Supernatant samples were diluted 1:10,000 in Reagent diluent (1% BSA in PBS). Washes were performed with PBST using a BioRad Plate washer. Imaging of the ELISA assay was completed on the BioRad Microplate Spectrophotometer with absorbances recorded at 450 nm and 540 nm. For screening, relative hCG levels were quantified by normalizing the background-subtracted absorbance (450nm-540nm) to the total nuclei per well (from the matched live cell confocal imaging of each well).

#### High-throughput automated hCG screening at the Innovative Genomics Institute

Automated screening was performed in a fully contained BSL2 robotic enclosure with no human intervention during cell processing. A completely automated cell culture system was employed using automated liquid handling (Tecan Fluent), incubation (Thermofisher Scientific Cytomat 5), robotic arm handling (HighRes Biosolutions A Cell robot arm), and washing-dispensing (Agilent EL 406). On this system, liquid handling can achieve a volume tolerance of 2.0 % coefficient of variation (CV), and dead volume of typically less than 1uL; the washer-dispenser dispensing lines have at most a 2.5 % coefficient of variation (CV) with a dead volume of aspiration typically less than 2uL. Robotic liquid handling procedures for the child plate were calibrated such that the aspiration and dispense occur below the liquid level surface for total volume processing and the height was calibrated so the cells were untouched by the disposable tip and minimally disturbed by liquid flow.

Two protocols were developed for automated cell passaging and automated differentiation with supernatant collection using the HighRes Biosolutions Cellario scheduling software. BeWo cells were manually nucleofected with the arrayed CRISPR library as above and pre-seeded into a 96-well plate and cultured for 2 days prior to onboarding onto the automated system. Upon receipt, cells pre-seeded in one parental 96-well tissue culture plate (Corning, 96-well, clear, flat-bottom, tissue culture treated) were passaged onto three child 96-well format high-content imaging plates (PhenoPlate, 96-well, black, optically clear flat-bottom, tissue culture treated) using robotic liquid handling. With an automated washer-dispenser, media is aspirated fully, cells on the parent plate are washed with 100uL of Phosphate Buffer Saline (PBS), then 50uL of Accutase is dispensed. The cells are incubated for 10 minutes in a controlled environment set to 37 °C temperature, 90% relative humidity, and 21% O2 concentration. After incubation, the plate is shaken quickly for 10 seconds to dissociate cells. Through robotic liquid handling (Tecan Fluent), 220uL of media are aliquoted from a pre-prepared 96-well source trough into each well and the dissociated cells in the parent plate are mixed up and down for 20 cycles with a 96-tip channel robotic arm. After mixing, 80uL of cell solution are aliquoted sequentially from the parent plate into each child plate (pre-prepared with 20uL BeWo media) for a total volume of 100uL per well. Each child plate is incubated for 24 hours before initiating differentiation.

Differentiation is induced by transferring medium from a pre-prepared 96-well source trough (manually prepared to contain Differentiation medium (BeWo medium + 20uM Forskolin) in all wells except the 3 DMSO-only wells, which contain BeWo medium + 0.1% DMSO) to the child plates in such a way that the cells are minimally disturbed by the process. This protocol is processed sequentially per child plate and occurs every 24 hours for a total differentiation time of 48 hours per plate. Each child plate remains in incubation (37 °C temperature, 90 % relative humidity, and 21 % O2 concentration) between protocol execution steps. After 48 hours of differentiation, supernatant is collected for ELISA using a 96-tip channel robotic arm that aspirates 20uL of supernatant from the top of each plate and dispenses into a PCR plate and is then heat-sealed by an automated sealer and then output from the system.

### CRISPR Screen analysis

The BeWo CRISPR screen imaging was performed across 5 biological plate groups (parental sgRNA plates), with 3 technical replicates (A, B, C) per group, totaling 15 plates. Primary image analysis was performed in Perkin Elmer Harmony software. To quantify Fusion Index in high throughput in BeWo cells, the Perkin Elmer Harmony software was used to perform bright-field correction, and basic flat-field correction for each field of view. Nuclei were segmented using the “find nuclei” function and method C and selected for area >50 μm^2^ (to eliminate debris). The GFP+ and mScarlet+ regions in each image were segmented using an intensity threshold and an area threshold of 100 μm^2^. To avoid artificial regions of mScarlet and GFP at nearby cell boundaries, all selected mScarlet and GFP regions were resized by eroding the regions by 5 pixels. The pixel-based percentage overlap of selected nuclei with the resized GFP+ or mScarlet+ regions was calculated using the “cross population” function. Nuclei were then filtered using a Boolean operator, and the Fusion Index was calculated as the proportion of nuclei with at least 90% pixel overlap with both mScarlet and GFP regions (fused nuclei) over the proportion of nuclei with at least 70% pixel overlap with either mScarlet or GFP regions (total fluorescent nuclei). Secondary phenotypes were calculated explicitly in the Harmony Software. For the Cell Paint analysis, we used the segmented nuclei from the cell-cell fusion pipeline, then used “Find Cytoplasm” with Method D, and then multi-parameter features across all channels were extracted using the “Cell Paint” analysis pipeline in the Harmony software with default parameters (Default Regions, Property set = Basic, and SER Scale = 2). In addition, hCG levels normalized to the total nuclei per well were added to the Excel spreadsheets and analyzed alongside imaging features using a custom R script. Replicate correlations were assessed using linear regression (R²) on raw well-level values with no exclusions. Wells were then excluded from downstream analysis if the percent of fluorescent nuclei over total nuclei was <45% (indicating cell death or lack of expression). Plates were excluded from Cell Painting analysis if fewer than 3 negative control wells or fewer than 3 positive control wells passed viability filtering. Fusion index and hCG secretion values were normalized to percent inhibition (PI) independently and internally within each plate-replicate group using the formula:

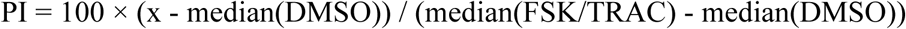

where x represents the raw well measurement, and medians were computed from viable control wells within the same plate. This normalization scaled DMSO (0% differentiation) to 0 and FSK/TRAC (maximal differentiation) to 100. For Cell Paint analysis, from the initial feature matrix (>900 morphological parameters per well), non-biological columns were excluded: metadata fields (row, column, timepoint), instrument environmental parameters (temperature, CO₂), and plate-level identifiers. Features with near-zero variance (NZV) were identified and removed by computing the median absolute deviation (MAD) of each feature across positive control wells within each plate-replicate group and features with MAD < 0.0001 × global median were flagged as uninformative and excluded. This yielded a final feature set of 866 biologically informative morphological parameters. Cell Painting features were normalized to robust Z-scores independently per plate-replicate group, using positive control wells (FSK/TRAC) as the reference population:

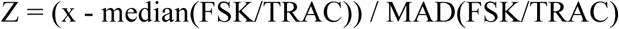

where MAD is the median absolute deviation with the standard 1.4826 scaling factor. Prior to PCA, Z-scores were winsorized to ±10 to limit the influence of extreme outliers. PCA was performed on the normalized feature matrix, retaining principal components (PCs) that cumulatively explained 90% of variance (40 components). A signed differentiation score was computed for each well by projecting its PC coordinates onto the axis connecting the negative control (DMSO) centroid to the positive control (FSK/TRAC) centroid in PC space:

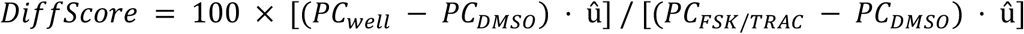

where û is the unit vector along the DMSO→FSK/TRAC axis. This metric quantifies the degree of cellular differentiation relative to controls, with 0 representing DMSO-like state and 100 representing maximal FSK/TRAC-like differentiation. For each gene, median values across all viable sample wells (pooled across technical replicates) were computed for Fusion Index PI, hCG PI, and Differentiation Score. Gene-level MAD Z-scores were calculated for each readout to identify outliers relative to the population distribution. Genes were classified as hits if their |MAD Z-score| ≥ 2.0 for at least one of the three primary readouts (Fusion, hCG, or Differentiation Score). Combinatorial hit categories were assigned based on which readout(s) exceeded threshold (e.g., “Fusion+hCG”, “Diff only”). Hit genes were subjected to unsupervised hierarchical clustering using complete linkage and Euclidean distance on their capped MAD Z-score profiles (±8 maximum) across all three primary readouts. The resulting dendrogram was cut to yield k=4 clusters, grouping genes with similar multi-phenotypic signatures. Phenotypic profiles were displayed as Cartesian-coordinate radar plots with axes representing capped MAD Z-scores (±5) for each readout, enabling visual comparison of multi-dimensional phenotypes across genes, clusters, and hit categories. All analyses were performed in R (version 4.3.2) using the following packages: dplyr (version 1.1.4), tidyr (version 1.3.1), ggplot2 (version 3.5.2) for data manipulation and visualization; ComplexHeatmap (version 2.18.0) for clustering heatmaps; eulerr for set overlap diagrams; and patchwork (version 1.2.0) for multi-panel figure composition.

### TIDE PCR and ICE analysis

Genomic DNA from BeWo cells was purified using the Zymo Genomic DNA miniprep kit (Zymo, D7001). PCR amplification was performed using the primers below using PrimeSTAR HS DNA polymerase (Takara Bio) or KOD One (Sigma Aldrich) PCR master mix, according to manufacturer’s instructions. Sanger sequencing was performed by Elim Biopharm or Genewiz. TIDE Indel % and KO Score were evaluated using Sanger sequencing .ab1 files with Synthego’s ICE Analysis tool (https://ice.editco.bio/#/).

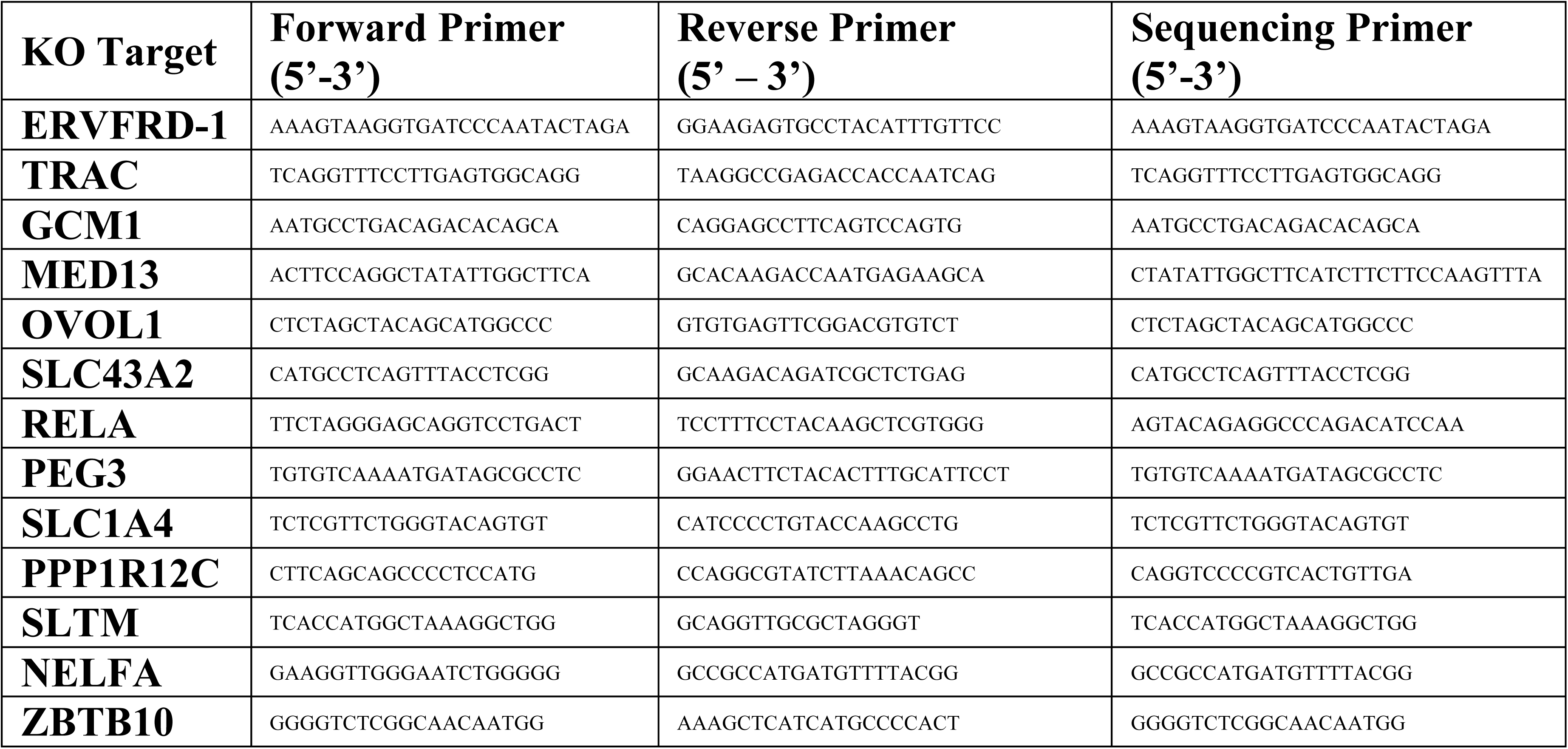

### Polyclonal KO using Cas9 RNPs in BeWo and hTSCs

Guide RNA sequences used:

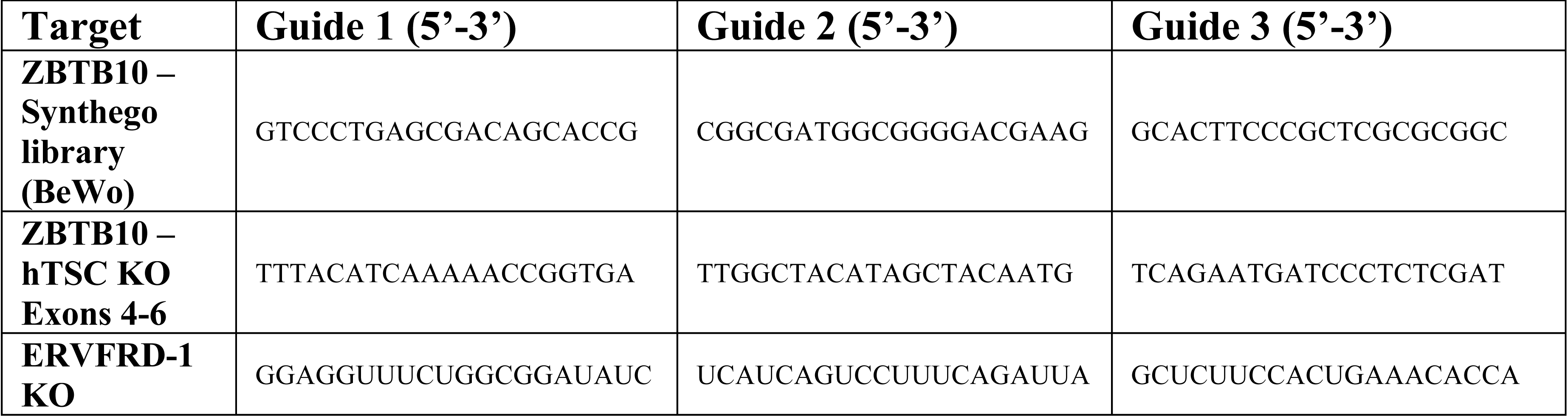

Guide RNAs were ordered from Synthego or IDT and resuspended to 100uM in TE buffer and stored at −80C. Cas9 RNP nucleofections were performed by assembling guide RNAs at a ratio of 3:1 with Cas9. For hTSCs, 15pmoles of RNP (45pmoles sgRNA plus 15pmoles of Cas9 enzyme) were used and for BeWo 10pmoles of RNP (30pmoles sgRNA plus 10pmoles Cas9) were used. Each reaction was mixed up to 20uL total volume with Lonza SG nucleofector solution (pre-mixed with 1x Supplement 1 per manufacturer’s instructions). RNP assembly was performed by incubating at room temperature for 10 minutes before adding cells. For BeWo knockout, 1E5 cells were used per reaction and for hTSCs, 2E5 cells were used per reaction. Cells were resuspended in 5uL cell media and added to the nucleofection/RNP solution then nucleofected with a Lonza 4D nucleofector using the SG buffer setting and code CA-137. Immediately after nucleofection, 80uL media was added and cells were allowed to recover for 10 minutes at 37C, 5% CO2 before transferring to a plate for growth. Knockout hTSCs were used within 1-2 passages of Cas9 RNP nucleofection for downstream assays.

### Cell and organoid growth assays

The confluency of TSCs and size of trophoblast organoids was assessed using CELLCYTE Studio (CYTENA) housed inside a 37 °C, 5% CO_2_ incubator. Organoids were imaged in Matrigel domes using “Spheroid” mode. Confluency area measurements and organoid size were segmented and calculated using the built-in Cellcyte software. The growth of BeWo cells was measured using an Incucyte S3 (Sartorius) housed inside a 37 °C, 5% CO_2_ incubator and confluency calculated using built-in Incucyte software.

### ELISA Assay for hCG**β** quantification

WT and ZBTB10 KO BeWo cells were plated at a density of 3E5 cells/well in a 6-well plate and treated with 0.1% DMSO or 2uM FSK. After 48-hours treatment, 200μL of cell supernatant was collected and stored in low-binding Eppendorf tubes at −80°C until day of ELISA assay. For hTSCs, cells transfected with Cas9 RNPs then allowed to grow and recover for 2-3 days, then plated at a density of 3E4 cells/well in glass-bottom 4-well Ibidi chambers (catalog #: 80426). Cells were then differentiated in STB Media for 6 days (changing media at 3 days) and 200uL of supernatant collected from each well. The hCGβ ELISA (Catalog #: DY9034-05) was performed according to manufacturer’s protocol using 96-well Immulon 4HBX Flat Bottom Plate (Catalog #: 3855). Supernatant samples were diluted 1:2,000-1:10,000 in Reagent diluent (1% BSA in PBS) and the dilution factor was used to calculate the final supernatant concentration. Washes were performed with PBST using a BioRad Plate washer. Imaging of the ELISA assay was completed on the BioRad Microplate Spectrophotometer with absorbances recorded at 450 nm and 540 nm. To quantify interpolated hCG levels, the average of 2 standard curves was used and a Log-Log plot of background-subtracted absorbance vs. hCG concentration was fit using linear regression.

### Cell-cell fusion assays

Two-color BeWo fusion assays were performed in 96-well plates using 10,000 cells in a 50:50 mixture as described previously.^46^ The same plasmids were used to generate GFP and mScarlet lentivirus and two-color hTSC stable cell lines were created by transducing hTSCs in TSC media and selecting with 62.5ug/mL Hygromycin for at least 10 days. Two-color hTSC fusion assays were performed by plating 3E4 cells in a 50:50 mixture into Ibidi glass-bottom 4-well dishes pre-coated with Laminin-521. Cells were then differentiated into STB for 6 days as described above and fixed at Day 7, or undifferentiated and fixed when confluent at Day 3-5. hTSCs were then stained with DAPI for 30 minutes in PBS, washed twice in PBS, and imaged on an ECHO Revolution (BICO) widefield microscope using the 4X Air objective. Fusion Index in BeWo cells was calculated exactly as in CRISPR screening using the Harmony software. For hTSCs, a similar approach was employed using a custom python script with nuclear segmentation using Cellpose version 3.0.10 (model = cyto2, diameter = 30 pixels, flow threshold = 0.4, and cell probability threshold = 0.0), user-based intensity thresholding of mScarlet and GFP channels, and then calculation of the Fusion Index as above. Quantification of unfused nuclei using Phalloidin staining was performed by manual annotation with the annotator blinded to the imaging condition. Only nuclei with complete, contiguous Phalloidin borders were counted as “unfused” and the rest were counted as fused. Total nuclei were segmented with Cellpose, and the number of annotated unfused nuclei was divided by the total detected nuclei.

### RNA-Seq

BeWo cells were plated at a concentration of 3E5 cells/well in a 6-well plate, then treated for 48 hours with 0.1% DMSO or 20µM FSK with media changed every 24 hours. BeWo cells were harvested using 500µL Trizol followed by two sequential chloroform extractions followed by ethanol precipitation per the Trizol user guide (Thermo Fisher MAN0001271). hTSCs were plated at a density of 1E5 cells/well in Cultrex-coated 6-well plates and differentiated into EVT or STB as above and harvested at Day 7, or if undifferentiated were harvested when confluent at Day 4. hTSC RNA was harvested using the RNeasy mini kit (Qiagen, 74104). For BeWo RNA-Seq, total RNA was depleted of rRNA using the Illumina rRNA depletion kit (NEB E6310) and then prepared for Illumina sequencing using the NEBNext Ultra II directional RNA library preparation kit for Illumina (E7760) per the manufacturer’s protocol. Sequencing was performed by MedGenome on an Illumina NovaSeq with paired-end 150 bp reads. Fastq files were assessed by FASTQC and then pseudoaligned to the hg38 transcriptome using Salmon to quantify transcripts in mapping-based mode (salmon quant -i) using decoy-aware mapping with gencode.v39.transcripts.fa. For hTSCs, purified RNA was sent to Plasmidsaurus which uses Illumina NovaSeq sequencing and a 3’ end counting approach. Plasmidsaurus performed read-filtering using FastP v0.24.0: poly-X tail trimming, 3’ quality-based tail trimming, a minimum Phred quality score of 15, and a minimum length requirement of 50 bp then alignment to hg38 reference genome using STAR aligner v2.7.11^81^ with non-canonical splice junction removal and output of unmapped reads. Gene-expression quantification using featureCounts (subread package v2.1.1) with strand-specific counting, multi-mapping read fractional assignment, exons and three prime UTR as the feature identifiers, and grouped by gene_id. Final gene counts were annotated with gene biotype and other metadata extracted from the reference GTF file. For all datasets, differential gene expression analysis was performed with EdgeR^82^ in Jupyter Notebook with filtering for low-expressed genes with edgeR::filterByExpr. Statistical comparisons between individual sample conditions were made using EdgeR makeContrasts and then glmQLFit. The reported P-values and FDR values were corrected with adjust.method = “BH,” and differentially expressed genes are reported with an adjusted FDR of <0.05. For visualization, the STAR program was used to map .fastq reads onto the hg38 genome to generate .bam files for each sample. These .bam files were converted to .bigWig files using bedtools and visualized using IGV Viewer.

### ChIP-Seq

Samples were sent to Active Motif (Carlsbad, CA) for ChIP-Seq. Active Motif prepared chromatin, performed ChIP reactions, generated libraries, sequenced the libraries, and performed basic analysis. In brief, two biological replicates each of human BeWo choriocarcinoma cells stably overexpressing N-terminally tagged Halo-3XFlag-ZBTB10 (integrated via Piggybac under an L30 promoter and Puro selection) were grown to ∼50% confluence then treated for 48 hours with 0.1% DMSO or 20uM Forskolin with media changed at 24 hours. Cells were fixed by adding formaldehyde to a final concentration of 1% in the existing medium and incubating for 15 min and then quenched with 0.125 M glycine. Chromatin was isolated by adding lysis buffer, followed by disruption with a Dounce homogenizer. Lysates were sonicated and the DNA sheared to an average length of 200-500 bp with Active Motif’s PIXUL*®* Multi-Sample Sonicator (catalog 53130). Genomic DNA (Input) was prepared by treating aliquots of chromatin with RNase, proteinase K and heat for de-crosslinking, followed by SPRI beads clean up (Beckman Coulter) and quantitation by Qubit (Thermofisher). Extrapolation to the original chromatin volume allowed determination of the total chromatin yield.

An aliquot of 30ug chromatin was precleared with protein G agarose beads (Invitrogen). Genomic DNA regions of interest were isolated using 4ug of antibody against ZBTB10 (Abcam, cat number ab117786, lot number 1036026-8). Complexes were washed, eluted from the beads with SDS buffer, and subjected to RNase and proteinase K treatment. Crosslinks were reversed by incubation overnight at 65°C, and ChIP DNA was purified by phenol-chloroform extraction and ethanol precipitation.

Illumina sequencing libraries were prepared from the ChIP and Input DNAs by the standard consecutive enzymatic steps of end-polishing, dA-addition, and adaptor ligation. Steps were performed on an automated system (Apollo 342, Takara PrepX DNA Library Kit). After a final PCR amplification step, the resulting DNA libraries were quantified and sequenced on Illumina’s NextSeq 2000 (PE50, 30 million reads).

#### ChIP-Seq analysis

Raw sequencing reads were assessed for quality using FastQC (v0.11.9)^83^ to evaluate per-base quality scores, nucleotide composition, adapter contamination, and overrepresented sequences. Adapter sequences and low-quality bases were trimmed using Trim Galore (v0.6.7)^84^ with default parameters. Trimmed reads were aligned to the reference genome (hg38) using BWA (v0.7.17)^85^, and alignments were converted to BAM format with SAMtools (v1.17).^86^ BAM files were subsequently filtered to remove duplicate reads (Picard, v3.0.0)^87^, non-primary alignments, unmapped reads, and reads mapping to blacklisted regions. Signal-enriched regions were identified using MACS2 (v2.2.7.1)^88^ with a q-value threshold of 0.1, using the corresponding input sample as background. Broad signal-enriched regions were identified using epic2 (v0.0.52)^89^ with a q-value threshold of 0.05, using the corresponding input sample as background. Peaks were annotated to the nearest gene and genomic feature using HOMER (v4.11).^90^ Known and de novo motif enrichment analyses were performed with HOMER (v4.11) on peak sets to identify enriched DNA-binding protein motifs. Fraction of reads in peaks (FRiP) was calculated as a quality control metric. Filtered BAM files were converted to bigWig format using deepTools (v3.5.1)^91^ and normalized as counts per million (CPM) for visualization.

### Western Blotting

Cells were trypsinized and quenched in medium and 1E6 cells were centrifuged at 200g for 2.5 min, resuspended in 150 µL of PBS plus 50 µL of 4x SDS-PAGE loading buffer (200mM Tris at pH 6.8, 400mM DTT, 10% B-ME, 8% SDS, 0.4% bromophenol blue, 40% glycerol), boiled on a heat block for 20 minutes at 100°C, and then centrifuged at 16,000g (Eppendorf 5415) for 3 minutes at 4°C. Supernatant was then transferred to a fresh tube and stored at −20°C. Cell lysate (35μL) was loaded onto 4%–20% Tris-glycine (Bio-Rad) and transferred at 100V for 1 hour onto 0.45μm nitrocellulose. A solution of 5% bovine serum albumin was used to block membranes for 1 h at room temperature prior to blotting and was used to dilute primary and secondary antibodies. Gels were imaged by addition of Western Lightning ECL reagent on a Chemidoc imaging system. Primary and secondary antibodies and concentrations are listed below.

### Antibodies

The antibodies used for western blotting were: hCGβ (1:1000; Abcam ab53087), ZBTB10 (Bethyl, A303-257A, Rabbit), Beta-actin (1:5000, Sigma, A2228-100UL), anti-rabbit HRP secondary (1:5000; Invitrogen 31462), and anti-mouse HRP secondary (1:5000; Invitrogen 31430). The antibodies used for immunofluorescence were: HLA-G (Cell Signaling, 79769, Rabbit), SDC1 (R&D, AF2780, Goat), ZBTB10 (Thermo, PA5-54448, Rabbit), TFAP2A (Santa Cruz, sc-12726, Mouse), KRT7 (Abcam, ab183344, Rabbit), GATA3 (Invitrogen, MA1-028, Mouse), hCG (Abcam, ab53087, Mouse), F-Actin (Phalloidin-Alexa594) (1:500, Invitrogen, A12381), Donkey anti-mouse Alexa-647 (Invitrogen A31571), Donkey anti-rabbit Alexa-488 (Invitrogen A21206), Donkey anti-goat Alexa-594 (Invitrogen A11058).

### Immunofluorescence & quantification

Cells were fixed with 4% paraformaldehyde (PFA) in PBS containing Ca²⁺ and Mg²⁺ (PBS++) for 30 minutes at room temperature. After three washes with PBS++, samples were blocked in Blocking buffer (3% normal donkey serum (Jackson ImmunoResearch #017-000-121) and 0.3% Triton X-100 in PBS) for 30 minutes. Primary antibodies were diluted 1:200 in Blocking buffer and incubated for 1-2 hours at room temperature. Following three 5-minute washes in IF Wash buffer (PBS++ containing 0.3% Triton X-100), samples were incubated with secondary antibodies diluted 1:1000 and DAPI (100 ng/ml; Invitrogen #D1306) in blocking buffer for 30 minutes at room temperature in the dark. After three final washes, samples were mounted using ProLong Antifade mounting medium (R&D Systems #4866-20) with 1.5H glass coverslips and imaged on an ECHO Revolution (BICO) widefield microscope using a 10X Air objective.

Trophoblast organoids were fixed in 4% PFA in PBS++ overnight at 4°C then washed 3 times in PBS for at least 30 minutes each at room temperature. Trophoblast organoids (TO), but not adherent EVT-TOs, were cleared by incubating in CUBIC-L Clearing Solution (TCI, T3740) for 2 days at 37C. Organoids were blocked in Blocking buffer for 3 hours then incubated with primary antibodies overnight at 4C. Samples were washed 3 times for 30min+ each in IF Wash buffer then incubated with secondary antibodies and DAPI (100 ng/ml; Invitrogen #D1306) in blocking buffer overnight at 4C in the dark. Samples were washed again 3 times for 30min+ each in IF Wash buffer then transferred to Ibidi glass-bottom dishes in CUBIC-R+ and imaged using a Zeiss scanning confocal microscope with 20X Air objective.

To quantify the percent of cells with nuclear ZBTB10 signal, multi-channel confocal images were analyzed by segmenting nuclei from the DAPI channel using Cellpose version 3.0.10 (model = cyto2, diameter = 30 pixels, flow threshold = 0.4, and cell probability threshold = 0.0). To quantify cytoplasmic GFP signal for comparison with nuclear signal, a perinuclear ring region was computationally defined for each nucleus. Euclidean distance transform was applied to the binary nuclear mask to generate a distance map from nuclear boundaries. The cytoplasmic ring was defined as the region beginning 3 pixels exterior to each nuclear boundary and extending to 5 pixels in thickness (i.e., 3–8 pixels from the nuclear edge). Ring pixels were assigned to their nearest nucleus based on the distance transform indices. Nuclei lacking valid ring regions (ring area = 0) due to crowding or image edge proximity were excluded from analysis. Mean GFP fluorescence intensity was calculated for both the nuclear region and the corresponding cytoplasmic ring region of each cell. The nuclear-to-cytoplasmic enrichment ratio was computed as the quotient of nuclear mean intensity divided by ring mean intensity. Cells were called as GFP-positive (nuclear-enriched) if nuclear GFP intensity > 1.2-fold cytoplasmic ring intensity. The percent of cells with nuclear ZBTB10+ signal was calculated as the number of GFP-positive (nuclear enriched) nuclei divided by the total nuclei.

To quantify fluorescent intensity in hTSCs, images were analyzed using FIJI (2.16.0) with a custom macro script for semi-automated region-of-interest (ROI) definition and intensity quantification. For STB, hCG was used as the region segmentation channel and for EVT, Phalloidin was used as the region segmentation channel. The segmentation channel was subjected to Gaussian blur filtering (σ = 2) and a binary mask was generated by applying a user-defined intensity threshold to segment marker-positive regions by manual inspection and converted to a binary mask. Positive regions in the binary mask were identified using the “Analyze Particles” function with a minimum size threshold of 100 pixels to exclude small debris. Mean fluorescence intensity was quantified within each individual ROI using ImageJ’s “Measure” function applied to all ROIs in the ROI Manager. Background intensity was then measured outside cells by manually selecting 4 square ROIs throughout the image. The average background was calculated and subtracted from all ROIs and the mean background-subtracted fluorescence intensity was then calculated across each image. For each condition, 4-6 images were obtained and stitched together and averaged. Three independent biological replicates were performed for each imaging experiment.

### Placental Tissue staining

Full thickness biopsies were collected from the placenta and washed in 1x PBS. These biopsies were embedded in Tissue-Tek OCT media (Sakura Finetek), froze in the vapor phase of LN2, and stored at −80°C. Tissue sections of 10μm were cut using a cryostat (Leica CM1850) and mounted onto Superfrost Plus Microscope slides (Fisherbrand), dried, and stored at −80C. Prior to staining slides were thawed to room temperature for 10 minutes, then OCT was dissolved by incubating in PBS++ for 20 minutes at room temperature. Tissue sections were outlined with a barrier pap pen (Thermo Fisher) and blocked in Blocking Buffer (3% normal donkey serum and 0.3% Triton X-100 in PBS) for 1 hour at room temperature. The above IF protocol was slightly modified for tissues: primary antibodies were incubated in a humidified chamber at 4C overnight and secondary antibodies were incubated for 1 hour at room temperature. Wash steps were extended to 15 minutes each. After washing, slides were then rinsed in PBS and treated with TrueBlack Lipofuscin Autofluorescence Quencher (Biotium) for background reduction according to manufacturer’s instructions before mounting with ProLong Gold Antifade and glass coverslips. Imaging of tissue was performed on a Zeiss confocal microscope with 20X Air objective.

All image processing and quantification were performed using Python 3.11.5 with scikit-image (version 0.20.0), NumPy (version 1.24.3), and SciPy (version 1.11.1) libraries. Nuclei were segmented using Cellpose version 3.0.10 (model = cyto2, diameter = 30 pixels, flow threshold = 0.4, and cell probability threshold = 0.0). Binary masks were generated by manually setting intensity thresholds for SDC1, HLA-G, or TFAP2A to accurately capture positive cells over background by visual inspection. These masks were subsequently dilated by 2 pixels using binary morphological operations to account for nuclei at marker boundaries. Nuclei were classified as SDC1-positive or HLAG-positive if ≥20% of its nuclear area overlapped with the SDC1 or HLA-G mask. Mean ZBTB10 fluorescence intensity was then quantified within each respective nucleus and then averaged for each cell type within an individual placenta sample, across 4 independent patient samples.

### Single-nucleus RNA-Seq (snRNA-Seq) analysis

Processed early and late pregnancy placental snRNA-Seq RDS objects were obtained directly from figshare repositories provided in the original paper and as such followed the same analysis outlined in their methods section.^59^ Cells were manually annotated by loading the RDS files into Seurat objects (v5.3.0) and interpreting FeaturePlot distributions of known marker genes for STB, CTB, fusing STB, and EVT in R (version 4.5.2)^92,93^. Plots displaying the annotated clusters were generated via DimPlot in Seurat. The resulting cells with non-zero log-normalized ZBTB10 counts were counted and divided by the total number of cells in each cluster to obtain the percentage of cells expressing ZBTB10 and plotted in ggplot2 (version 4.0.2).^94^ Median expression levels of ZBTB10 were generated by calculating the median values of log-normalized counts for each cluster and plotted via ggplot2.

### Pseudotime analysis

STB differentiation pseudotime was inferred from log-normalized 8-week human placenta snRNA-seq data (Seurat v5.5.0; R version 4.5.3 (2026 03-11)) from Wang et al. 2024.^59^ Six STB clusters were retained from the .rds file for pseudotime analysis: eCTB_fusion (cluster 8; n = 535), eSTB_nascent (cluster 7; n = 1,535), a Mature 1 branch (clusters 0 + 5; n = 7,350), and a Mature 2 branch (clusters 1 + 3; n = 7,233). Genes used for ordering were computed from Seurat FindAllMarkers (Wilcoxon rank-sum test, min. detection 0.1, log-fold-change >= 0.25, positive and negative markers, Benjamini-Hochberg adjusted p < 0.05; n = 4588 genes) between the four cell-type groups. The pseudotime trajectory was built using Monocle2 (v2.38.0) from the log-normalized matrix [log(1 + counts per 10,000)] with Gaussian (uninormal) expression family with estimated size factors, and the trajectory backbone was learned by DDRTree (v0.1.6) with reduceDimension, max_components = 2, norm_method = “none”; K = 136 nodes). Cell-level pseudotime and state were assigned with Monocle2’s standard orderCells over the full cell matrix with the trajectory rooted at the state containing the most eCTB_fusion nuclei. The DDRTree embedding was visualized in ggplot2 (v4.0.3).

### Single-nucleus ATAC-Seq analysis

#### Motif activity scoring

Publicly available 8-week human placental snATAC-seq peak-by-cell counts from Wang et al. 2024^59^ were loaded into a Seurat/Signac object and matched to the published cell metadata, including SnapATAC2 cluster labels, donor information, quality-control metrics, and UMAP coordinates. Motif activity was estimated using chromVAR through Signac. Peaks were converted to genomic ranges and filtered to chromosomes present in the hg38 reference genome. Regional sequence statistics were computed using RegionStats. Human transcription factor position frequency matrices from JASPAR2020 were combined with HOMER motifs identified from the BeWo ZBTB10 ChIP, including NRF-related and sequence-bias motifs. The combined motif set was added to the ATAC assay, and per-cell chromVAR deviation Z-scores were calculated. These chromVAR scores were visualized on the published snATAC-seq UMAP embedding to compare motif activity across trophoblast cell states.

#### Depth-adjusted NRF1 residual analysis

To evaluate whether NRF1 motif activity was independent of sequencing depth, NRF1 chromVAR deviation scores were regressed against per-cell fragment counts using a linear model with log-transformed fragment count as the predictor. The residuals from this model were interpreted as NRF1 activity after adjustment for library size. Residual values were then grouped by the published snATAC-seq cluster labels and visualized as boxplots to compare depth-adjusted NRF1 activity across clusters.

### Multi-species scRNA-Seq comparison

#### Datasets

We analyzed published single-nucleus (human) and single-cell RNA-seq (opossum, mouse, tenrec, guinea pig, macaque) datasets spanning six mammalian species from Stadtmaurer et al. 2025.^64^ Non-human eutherian and marsupial data were from Stadtmaurer et al. (GEO: GSE274701): opossum (*Monodelphis domestica*; 6,761 cells, 26 cell types), tenrec (*Echinops telfairi*; 29,652 cells, 35 cell types), guinea pig (*Cavia porcellus*; 18,131 cells, 23 cell types), and mouse (*Mus musculus*; 11,452 cells, 33 cell types). Human (*Homo sapiens*) data were from the Reproductive Cell Atlas (Arutyunyan et al., 2023; 325,665 cells, 42 cell types). Macaque (*Macaca fascicularis*) data were from Jiang et al. (GEO: GSE180637; MF217404 dataset; 217,404 total cells).^95^ For macaque, annotated embryonic cells were excluded, retaining 165,621 placental and decidual cells across 17 cell types. Cell type labels were used as provided by original authors.

#### Dimensionality reduction

For species with pre-computed embeddings (opossum, tenrec, guinea pig, human), UMAP coordinates were used as-is. For macaque and mouse, UMAPs were computed using Scanpy: raw counts were library-size normalized (target sum = 10,000), log(*x*+1)-transformed, and highly variable genes identified (3,000 genes; Seurat flavor for macaque, Seurat v3 for mouse). PCA retained 50 components; k-nearest neighbor graphs used the top 30 PCs (k = 30 for macaque, k = 15 for mouse). UMAP parameters: min_dist = 0.3 for macaque, default for mouse.

#### ZBTB10 ortholog identification

Orthologs were identified from h5ad feature metadata. Human: direct symbol match. Opossum (ENSMODG00000006584), guinea pig (ENSCPOG00000013700), and mouse (ENSMUSG00000069114; *Zbtb10*): matched via gene_names_full column encoding Ensembl Compara one-to-one orthology assignments. Tenrec (MSTRG.18207): matched via gene_name_refined. Macaque: matched via hsapiens_homolog_associated_gene_name.

#### Expression quantification

For datasets with raw counts, (mouse, opossum, tenrec, guinea pig), expression was library-size normalized (target sum = 10,000) and log(*x*+1)-transformed. For human and macaque, log-normalized values from the original datasheet were used directly. Per cell type, we computed: (1) mean log-normalized expression across all cells; (2) percentage of cells with raw count > 0. Cell types with < 5 cells were excluded.

### Declaration of generative AI and AI-assisted technologies in the manuscript preparation process

During the preparation of this work, the authors used Claude and Phylo Biomni in order to support code writing to analyze screening and genomics data. After using this tool/service, the authors reviewed and edited the content as needed and take full responsibility for the content of the published article.

## DATA AVAILABILITY

The following datasets produced in this publication are available through the NCBI GEO database: ZBTB10 ChIP-Seq (GEO accession: GSE336955), BeWo RNA-Seq (GEO accession: GSE336958), and hTSC RNA-Seq (GEO accession: GSE336960).

## ACKNOWLEDGEMENTS

We thank members of the Tjian-Darzacq, Yang, and Nobu Hamazaki laboratories for helpful discussions. We thank Mary West and Pingping He for help with protocols, automation, and training at the QB3 High-Throughput Screening Facility (HTSF) at University of California, Berkeley, which provided the Opera Phenix microscope, supported by the Office of the Director, NIH, under award number S10OD021828. The content is solely the responsibility of the authors and does not necessarily represent the official views of the National Institutes of Health. The Advanced Translational Genetics lab at the IGI is supported by CRISPR Cures for Cancer Initiative, Li Ka Shing Foundation and Shanahan Family Foundation. We thank the patients who donated their placentas to scientific research. M.Y. is supported by the NICHD Path to Independence Award (4R00HD107219-03) and Director’s New Innovator Award (1DP2HD121941-01). M.N.E. was supported by the H.S. Chau Foundation (2023-2025) and is currently a Fellow of The Jane Coffin Childs Memorial Fund for Medical Research. I.A.G was funded by the NIH under NICHD Grant # R24HD000836. Research was funded by startup funding from UW department of OB/GYN, ISCRM, and the Brotman Baty Institute (BBI). We thank the individual members of the Birth Defects Research Laboratory (BDRL) and their associated affiliations: Ian A. Glass^1^, Kimberly A. Aldinger^1,2^, Dan Doherty^1^, Ian G. Phelps^1^, Jennifer C. Dempsey^1^, Kevin J. Lee^1^, and Lucinda A. Cort^1^; ^1^University of Washington, Department of Pediatrics, ^2^Seattle Children’s Research Institute.

## CONTRIBUTIONS

M.N.E. conceptualized the study, designed the methodology, performed most experiments and formal analysis, visualized results, and wrote the first draft of the manuscript. R.B. supported characterization of ZBTB10 KO BeWo cells including performing TIDE, RNA-seq, and ELISA assays. H.L. provided supervision and project design during automation at the IGI. A.C. analyzed snRNA-Seq data. J.C. performed the snATAC-Seq analysis. T.T. developed and performed automated robotic screening protocols at the IGI. S.A.M., I.A.G., and the BDRL collected and provided placental tissue samples. D.H. and F.U. provided mentorship and supervision during automated CRISPR screening. X.D. and R.T. supervised the experiments in BeWo cells, supported funding of the project, and edited the manuscript. M.Y. provided training and supervised all experiments in trophoblast stem cells, organoids, and placental tissue, supported funding of the project, and edited the manuscript.

## SUPPLEMENTARY MATERIAL

**Figure S1. Library gene selection and screen controls.**

**Figure S2. Screen quality control and normalization.**

**Figure S3. Biological replicate validation of hCG screen results with high-throughput automation at the Innovative Genomics Institute.**

**Figure S4. Segmentation and in-depth phenotyping of screen wells.**

**Figure S5. BeWo growth curves and validation of MED13 impacts on fusion.**

**Figure S6. ZBTB10 knockout does not perturb trophoblast stem cell marker expression.**

**Figure S7. Extended analysis of ZBTB10 KO RNA-Seq in trophoblast stem cells.**

**Figure S8. Extended analysis of BeWo ZBTB10 ChIP and snATAC-Seq analysis of ZBTB10 motifs in human first trimester placenta.**

**Figure S9. Cluster validation and pseudo-time analysis of first trimester placenta.**

**Figure S10. Evolutionary analysis of ZBTB10 coding sequence and cell type-specific expression.**

**Table S1.** Gene_hit_summary.csv. Per-gene raw median Fusion Index and hCG secretion (normalized to per-well confluency), and median percent inhibition normalized values for all primary (fusion, hCG, differentiation score (Diff)) and secondary phenotypes (nuclear area, ++ nuclei nucleus-nucleus contact area % (Contact), summed DAPI intensity (DAPI sum), coefficient of variation in DAPI nuclei (DAPI CV), calculated MAD Z-scores for all six phenotypes, hit calling, and cluster assignment for all 3 primary phenotypes. DoublePos = double positive (fused) nuclei. Median_hCG_Raw = hCG ELISA absorbance divided by total # nuclei per well. Raw values indicate before plate-level percent inhibition normalization to DMSO and TRAC controls.

**Table S2.** CRISPR_all_genes_IGI.csv. Percent inhibition normalized hCG secretion, calculated MAD Z-scores, and hit calling for automated hCG screening performed at the Innovative Genomics Institute.

## REFERENCES

1. Maltepe, E. & Fisher, S. J. Placenta: The Forgotten Organ. 10.1146/annurev-cellbio-100814-125620 31, 523–552 (2015).

2. Burton, G. J. & Fowden, A. L. The placenta: a multifaceted, transient organ. Philosophical Transactions of the Royal Society B: Biological Sciences 370, 20140066 (2015).

3. Turco, M. Y. & Moffett, A. Development of the human placenta. Development (Cambridge) 146, (2019).

4. Renaud, S. J. & Jeyarajah, M. J. How trophoblasts fuse: an in-depth look into placental syncytiotrophoblast formation. Cellular and Molecular Life Sciences vol. 79 Preprint at 10.1007/s00018-022-04475-z (2022).

5. O’Brien, K. & Wang, Y. The Placenta: A Maternofetal Interface. Annu. Rev. Nutr. 43, 301–325 (2023).

6. Vargas, A. et al. Reduced expression of both syncytin 1 and syncytin 2 correlates with severity of preeclampsia. Reproductive Sciences 18, 1085–1091 (2011).

7. Mukherjee, I. et al. Oxidative stress-induced impairment of trophoblast function causes preeclampsia through the unfolded protein response pathway. Scientific Reports 2021 11:1 11, 1–20 (2021).

8. Shao, X. et al. Placental trophoblast syncytialization potentiates macropinocytosis via mTOR signaling to adapt to reduced amino acid supply. Proc. Natl. Acad. Sci. U. S. A. 118, (2021).

9. Ruebner, M. et al. Impaired cell fusion and differentiation in placentae from patients with intrauterine growth restriction correlate with reduced levels of HERV envelope genes. Journal of Molecular Medicine 2010 88:11 88, 1143–1156 (2010).

10. Li, X., Li, Z. H., Wang, Y. X. & Liu, T. H. A comprehensive review of human trophoblast fusion models: recent developments and challenges. Cell Death Discovery vol. 9 Preprint at 10.1038/s41420-023-01670-0 (2023).

11. Brosens, I., Pijnenborg, R., Vercruysse, L. & Romero, R. The “Great Obstetrical Syndromes” are associated with disorders of deep placentation. Am. J. Obstet. Gynecol. 204, 193–201 (2011).

12. Norwitz, E. R. Defective implantation and placentation: laying the blueprint for pregnancy complications. Reprod. Biomed. Online 13, 591–599 (2006).

13. Simpson, R. A., Mayhew, T. M. & Barnes, P. R. From 13 weeks to term, the trophoblast of human placenta grows by the continuous recruitment of new proliferative units: A study of nuclear number using the disector. Placenta 13, 501–512 (1992).

14. Hertig, A. T., Rock, J. & Adams, E. C. A description of 34 human ova within the first 17 days of development. American Journal of Anatomy 98, (1956).

15. Blond, J.-L. et al. An Envelope Glycoprotein of the Human Endogenous Retrovirus HERV-W Is Expressed in the Human Placenta and Fuses Cells Expressing the Type D Mammalian Retrovirus Receptor. J. Virol. 74, 3321–3329 (2000).

16. Mi, S. et al. Syncytin is a captive retroviral envelope protein involved in human placental morphogenesis. Nature 2000 403:6771 403, 785–789 (2000).

17. Blaise, S., De Parseval, N., Bénit, L. & Heidmann, T. Genomewide screening for fusogenic human endogenous retrovirus envelopes identifies syncytin 2, a gene conserved on primate evolution. Proc. Natl. Acad. Sci. U. S. A. 100, 13013–13018 (2003).

18. Esnault, C. et al. A placenta-specific receptor for the fusogenic, endogenous retrovirus-derived, human syncytin-2. Proc. Natl. Acad. Sci. U. S. A. 105, (2008).

19. Oike, A., Shibata, S., Arima, T. & Okae, H. Syncytin-1 Is Responsible for the Fusion Between Human Trophoblasts and Endometrial Stromal Cells. Dev. Growth Differ. 67, 270–278 (2025).

20. Ahmadi, S. M., Perez, M. L. & Guardia, C. M. Secretion of placental peptide hormones: functions and trafficking. Front. Endocrinol. (Lausanne). 16, 1584303 (2025).

21. Gerbaud, P., Taskén, K. & Pidoux, G. Spatiotemporal regulation of cAMP signaling controls the human trophoblast fusion. Front. Pharmacol. 6, (2015).

22. Liang, C. Y. et al. GCM1 Regulation of the Expression of Syncytin 2 and Its Cognate Receptor MFSD2A in Human Placenta. Biol. Reprod. 83, 387–395 (2010).

23. Orendi, K., Gauster, M., Moser, G., Meiri, H. & Huppertz, B. The choriocarcinoma cell line BeWo: syncytial fusion and expression of syncytium-specific proteins. Reproduction 140, 759–766 (2010).

24. Wice, B., Menton, D., Geuze, H. & Schwartz, A. L. Modulators of cyclic AMP metabolism induce syncytiotrophoblast formation in vitro. Exp. Cell Res. 186, 306–316 (1990).

25. Pattillo, R. A. & Gey, G. O. The Establishment of a Cell Line of Human Hormone-synthesizing Trophoblastic Cells in Vitro. Cancer Res. 28, 1231–1236 (1968).

26. Malhotra, S. S., Suman, P. & Gupta, S. K. Alpha or beta human chorionic gonadotropin knockdown decrease BeWo cell fusion by down-regulating PKA and CREB activation. Scientific Reports 2015 5:1 5, 1–15 (2015).

27. Azar, C. et al. RNA-Seq identifies genes whose proteins are upregulated during syncytia development in murine C2C12 myoblasts and human BeWo trophoblasts. Physiol. Rep. 9, e14671 (2021).

28. Leduc, K. et al. Leukemia inhibitory factor regulates differentiation of trophoblastlike BeWo cells through the activation of JAK/STAT and MAPK3/1 MAP kinase-signaling pathways. Biol. Reprod. 86, (2012).

29. Ruebner, M. et al. Regulation of the human endogenous retroviral Syncytin-1 and cell–cell fusion by the nuclear hormone receptors PPARγ/RXRα in placentogenesis. J. Cell. Biochem. 113, 2383–2396 (2012).

30. Santinha, A. J., Strano, A. & Platt, R. J. Methods and applications of in vivo CRISPR screening. Nature Reviews Genetics 2025 26:10 26, 702–718 (2025).

31. Chow, R. D. & Chen, S. Cancer CRISPR Screens In Vivo. Trends in Cancer vol. 4 Preprint at 10.1016/j.trecan.2018.03.002 (2018).

32. Schraivogel, D., Steinmetz, L. M. & Parts, L. Pooled Genome-Scale CRISPR Screens in Single Cells. Annu. Rev. Genet. 57, 223–244 (2023).

33. Datlinger, P. et al. Pooled CRISPR screening with single-cell transcriptome readout. Nature Methods 2017 14:3 14, 297–301 (2017).

34. Yan, X. et al. High-content imaging-based pooled CRISPR screens in mammalian cells. J. Cell Biol. 220, (2021).

35. Kanfer, G. et al. Image-based pooled whole-genome CRISPRi screening for subcellular phenotypes. Journal of Cell Biology 220, (2021).

36. Shimizu, T. et al. CRISPR screening in human trophoblast stem cells reveals both shared and distinct aspects of human and mouse placental development. Proceedings of the National Academy of Sciences 120, e2311372120 (2023).

37. Zhang, H. et al. Development of a split-toxin CRISPR screening platform to systematically identify regulators of human myoblast fusion. Nature Communications 2025 17:1 17, 547-(2026).

38. Chan, C. W. F. et al. High-throughput screening of genetic and cellular drivers of syncytium formation induced by the spike protein of SARS-CoV-2. *Nat*. Biomed. Eng. 8, 291–309 (2024).

39. Baczyk, D. et al. Glial cell missing-1 transcription factor is required for the differentiation of the human trophoblast. Cell Death & Differentiation 2009 16:5 16, 719–727 (2009).

40. Krendl, C. et al. GATA2/3-TFAP2A/C transcription factor network couples human pluripotent stem cell differentiation to trophectoderm with repression of pluripotency. Proc. Natl. Acad. Sci. U. S. A. 114, E9579–E9588 (2017).

41. Papuchova, H. & Latos, P. A. Transcription factor networks in trophoblast development. Cellular and Molecular Life Sciences 2022 79:6 79, 337-(2022).

42. Dong, C. et al. A genome-wide CRISPR-Cas9 knockout screen identifies essential and growth-restricting genes in human trophoblast stem cells. Nature Communications 2022 13:1 13, 1–16 (2022).

43. Renaud, S. J. et al. OVO-like 1 regulates progenitor cell fate in human trophoblast development. Proc. Natl. Acad. Sci. U. S. A. 112, E6175–E6184 (2015).

44. Siggs, O. M. & Beutler, B. The BTB-ZF transcription factors. Cell Cycle 11, 3358 (2012).

45. Li, X. et al. STK40 inhibits trophoblast fusion by mediating COP1 ubiquitination to degrade P57Kip2. J. Transl. Med. 22, 852 (2024).

46. Esbin, M. N. et al. TFEB controls expression of human syncytins during cell–cell fusion. Genes Dev. (2024) doi:10.1101/gad.351633.124.

47. Zheng, W. et al. TFEB safeguards trophoblast syncytialization in humans and mice. Proceedings of the National Academy of Sciences 121, e2404062121 (2024).

48. Cesana, M. et al. TFEB controls syncytiotrophoblast formation and hormone production in placenta. Cell Death & Differentiation 2024 31:11 31, 1439–1451 (2024).

49. Karvas, R. M. et al. Stem-cell-derived trophoblast organoids model human placental development and susceptibility to emerging pathogens. Cell Stem Cell 29, 810–825.e8 (2022).

50. Okae, H. et al. Derivation of Human Trophoblast Stem Cells. Cell Stem Cell 22, 50–63.e6 (2018).

51. Wang, D. et al. Chromosomal instability in human trophoblast stem cells and placentas. Nature Communications 2025 16:1 16, 3918-(2025).

52. Varberg, K. M. et al. Extravillous trophoblast cell lineage development is associated with active remodeling of the chromatin landscape. Nature Communications 2023 14:1 14, 4826-(2023).

53. West, R. C. et al. Dynamics of trophoblast differentiation in peri-implantation–stage human embryos. Proceedings of the National Academy of Sciences 116, 22635–22644 (2019).

54. Tillotson, L. G. RIN ZF, a novel Zinc finger gene, encodes proteins that bind to the CACC element of the gastrin promoter. Journal of Biological Chemistry 274, 8123–8128 (1999).

55. Señarís, R. et al. Synthesis of Leptin in Human Placenta. Endocrinology 138, 4501–4504 (1997).

56. Rumer, K. K., Sehgal, S., Kramer, A., Bogart, K. P. & Winn, V. D. The effects of leptin on human cytotrophoblast invasion are gestational age and dose-dependent. Front. Endocrinol. (Lausanne). 15, 1386309 (2024).

57. Barrientos, G. et al. Leptin promotes HLA-G expression on placental trophoblasts via the MEK/Erk and PI3K signaling pathways. Placenta 36, 419–426 (2015).

58. Shukla, V. et al. NOTUM-mediated WNT silencing drives extravillous trophoblast cell lineage development. Proc. Natl. Acad. Sci. U. S. A. 121, e2403003121 (2024).

59. Wang, M. et al. Single-nucleus multi-omic profiling of human placental syncytiotrophoblasts identifies cellular trajectories during pregnancy. Nature Genetics 2024 56:2 56, 294–305 (2024).

60. Lee, S. U. & Maeda, T. POK/ZBTB proteins: An emerging family of proteins that regulate lymphoid development and function. Immunol. Rev. 247, 107–119 (2012).

61. Maeda, T. Regulation of hematopoietic development by ZBTB transcription factors. Int. J. Hematol. 104, 310 (2016).

62. Liongue, C., Almohaisen, F. L. J. & Ward, A. C. B Cell Lymphoma 6 (BCL6): A Conserved Regulator of Immunity and Beyond. Int. J. Mol. Sci. 25, 10968 (2024).

63. Fernando, T. M. et al. BCL6 evolved to enable stress tolerance in vertebrates and is broadly required by cancer cells to adapt to stress. Cancer Discov. 9, 662–679 (2019).

64. Stadtmauer, D. J. et al. Cell type and cell signalling innovations underlying mammalian pregnancy. Nature Ecology & Evolution 2025 9:8 9, 1469–1486 (2025).

65. Skogler, J. et al. Association between human chorionic gonadotropin (hCG) levels and adverse pregnancy outcomes: A systematic review and meta-analysis. Pregnancy Hypertens. 34, 124–137 (2023).

66. Takeda, K. et al. Placental but Not Heart Defects Are Associated with Elevated Hypoxia-Inducible Factor α Levels in Mice Lacking Prolyl Hydroxylase Domain Protein 2. Mol. Cell. Biol. 26, 8336–8346 (2006).

67. Li, Q., Wei, X., Zhang, Y., Zeng, W. & Lin, Y. Trophoblast adaptation to hypoxia: balance and dysfunction. Cell Commun. Signal. 23, 538 (2025).

68. Keenen, M. M. et al. Comparative analysis of the syncytiotrophoblast in placenta tissue and trophoblast organoids using snRNA sequencing. Elife 13, (2025).

69. Gong, S. et al. The RNA landscape of the human placenta in health and disease. Nature Communications 2021 12:1 12, 2639-(2021).

70. Elgazzaz, M., Brawley, A., Moronge, D. & Faulkner, J. L. Emerging Role of Leptin in Vascular and Placental Dysfunction in Preeclampsia. Arterioscler. Thromb. Vasc. Biol. 45, 585–599 (2025).

71. Simoni, M. K. et al. Type I interferon exposure of an implantation-on-a-chip device alters invasive extravillous trophoblast function. Cell Rep. Med. 6, 101991 (2025).

72. Jantine Van Voorden, A. et al. The pro-inflammatory cytokines IFN-α and TNF-α inhibit organoid-derived extravillous trophoblast invasion. *bioRxiv* 2025.08.15.670497 (2025) doi:10.1101/2025.08.15.670497.

73. Kim, J. et al. Isolation and characterization of mammalian homologs of the Drosophila gene glial cells missing. Proc. Natl. Acad. Sci. U. S. A. 95, 12364–12369 (1998).

74. Bainbridge, S. A. et al. Effects of reduced Gcm1 expression on trophoblast morphology, fetoplacental vascularity, and pregnancy outcomes in mice. Hypertension 59, (2012).

75. Hayashi, Y. et al. Glial cells missing 1 triggers gliosis and angiogenesis after neonatal brain injury. iScience 28, 113860 (2025).

76. Iwasaki, Y. et al. The potential to induce glial differentiation is conserved between Drosophila and mammalian glial cells missing genes. Development 130, 6027–6035 (2003).

77. Turco, M. Y. et al. Trophoblast organoids as a model for maternal–fetal interactions during human placentation. Nature 2018 564:7735 564, 263–267 (2018).

78. Sheridan, M. A. et al. Establishment and differentiation of long-term trophoblast organoid cultures from the human placenta. Nat. Protoc. 15, 3441–3463 (2020).

79. Zhou, J. et al. Development of apical out trophoblast stem cell derived organoids to model early human pregnancy. iScience 28, 112099 (2025).

80. Yang, L., Liang, P., Yang, H. & Coyne, C. B. Trophoblast organoids with physiological polarity model placental structure and function. J. Cell Sci. 137, jcs261528 (2024).

81. Dobin, A. et al. STAR: ultrafast universal RNA-seq aligner. Bioinformatics 29, 15–21 (2013).

82. Robinson, M. D., McCarthy, D. J. & Smyth, G. K. edgeR: a Bioconductor package for differential expression analysis of digital gene expression data. Bioinformatics 26, 139 (2010).

83. Babraham Bioinformatics - FastQC A Quality Control tool for High Throughput Sequence Data. https://www.bioinformatics.babraham.ac.uk/projects/fastqc/.

84. Babraham Bioinformatics - Trim Galore! https://www.bioinformatics.babraham.ac.uk/projects/trim_galore/.

85. Li, H. & Durbin, R. Fast and accurate short read alignment with Burrows–Wheeler transform. Bioinformatics 25, 1754–1760 (2009).

86. Li, H. et al. The Sequence Alignment/Map format and SAMtools. Bioinformatics 25, (2009).

87. Picard Tools - By Broad Institute. https://broadinstitute.github.io/picard/.

88. Zhang, Y. et al. Model-based Analysis of ChIP-Seq (MACS). Genome Biology 2008 9:9 9, R137-(2008).

89. Stovner, E. B. & Sætrom, P. epic2 efficiently finds diffuse domains in ChIP-seq data. Bioinformatics 35, 4392–4393 (2019).

90. Heinz, S. et al. Simple Combinations of Lineage-Determining Transcription Factors Prime cis-Regulatory Elements Required for Macrophage and B Cell Identities. Mol. Cell 38, 576–589 (2010).

91. Ramírez, F. et al. deepTools2: a next generation web server for deep-sequencing data analysis. Nucleic Acids Res. 44, (2016).

92. R: A Language and Environment for Statistical Computing Reference Index The R Core Team. https://www.gnu.org/copyleft/gpl.html.

93. Hao, Y. et al. Dictionary learning for integrative, multimodal and scalable single-cell analysis. Nature Biotechnology 2023 42:2 42, 293–304 (2023).

94. Wickham, H. ggplot2. Wiley Interdiscip. Rev. Comput. Stat. 3, 180–185 (2011).

95. Jiang, X. et al. Identifying a dynamic transcriptomic landscape of the cynomolgus macaque placenta during pregnancy at single-cell resolution. Dev. Cell 58, 806–821.e7 (2023).

