## Supplementary material for "Phenotypic CRISPR screening identifies ZBTB10 as a novel regulator of human trophoblast differentiation": Figure S1

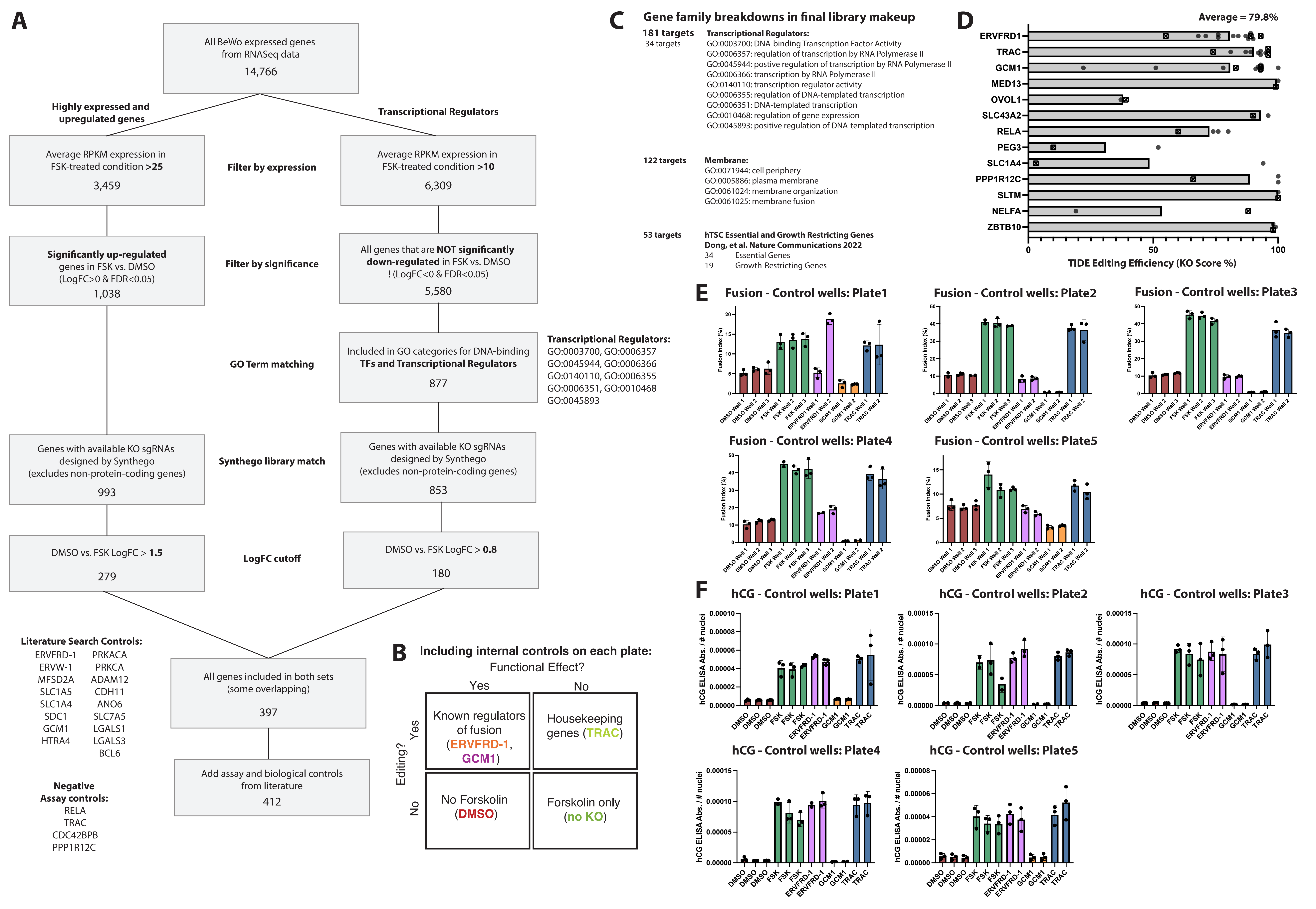

A
All BeWo expressed genes from RNASeq data
14,766
Highly expressed and upregulated genes
Transcriptional Regulators
Average RPKM expression in FSK-treated condition >25
3,459
Filter by expression
Average RPKM expression in FSK-treated condition >10
6,309
Significantly up-regulated genes in FSK vs. DMSO (LogFC>0 & FDR<0.05)
1,038
Filter by significance
All genes that are NOT significantly down-regulated in FSK vs. DMSO ! (LogFC<0 & FDR<0.05)
5,580
GO Term matching
Included in GO categories for DNA-binding TFs and Transcriptional Regulators
877
Transcriptional Regulators:
GO:0003700, GO:0006357
GO:0045944, GO:0006366
GO:0140110, GO:0006355
GO:0006351, GO:0010468
GO:0045893
Genes with available KO sgRNAs designed by Synthego (excludes non-protein-coding genes)
993
Synthego library match
Genes with available KO sgRNAs designed by Synthego (excludes non-protein-coding genes)
853
DMSO vs. FSK LogFC > 1.5
279
LogFC cutoff
DMSO vs. FSK LogFC > 0.8
180
Literature Search Controls:
ERVFRD-1 PRKACA
ERVW-1 PRKCA
MFSD2A ADAM12
SLC1A5 CDH11
SLC1A4 ANO6
SDC1 SLC7A5
GCM1 LGALS1
HTRA4 LGALS3
BCL6
All genes included in both sets (some overlapping)
397
Negative Assay controls:
RELA
TRAC
CDC42BPB
PPP1R12C
Add assay and biological controls from literature
412
B
Including internal controls on each plate:
Functional Effect?
Yes
No
Editing?
Yes
No
Known regulators of fusion (ERVFRD-1, GCM1)
Housekeeping genes (TRAC)
No Forskolin (DMSO)
Forskolin only (no KO)
C
Gene family breakdowns in final library makeup
181 targets
34 targets
Transcriptional Regulators:
GO:0003700: DNA-binding Transcription Factor Activity
GO:0006357: regulation of transcription by RNA Polymerase II
GO:0045944: postive regulation of transcription by RNA Polymerase II
GO:0006366: transcription by RNA Polymerase II
GO:0140110: transcription regulator activity
GO:0006355: regulation of DNA-templated transcription
GO:0006351: DNA-templated transcription
GO:0010468: regulation of gene expression
GO:0045893: positive regulation of DNA-templated transcription
122 targets
Membrane:
GO:0071944: cell periphery
GO:0005886: plasma membrane
GO:0061024: membrane organization
GO:0061025: membrane fusion
53 targets
hTSC Essential and Growth Restricting Genes
Dong, et al. Nature Communications 2022
34 Essential Genes
19 Growth-Restricting Genes
D
Average = 79.8%
ERVFRD1
TRAC
GCM1
MED13
OVOL1
SLC43A2
RELA
PEG3
SLC1A4
PPP1R12C
SLTM
NELFA
ZBTB10
0
50
100
TIDE Editing Efficiency (KO Score %)
E
Fusion - Control wells: Plate1
Fusion - Control wells: Plate2
Fusion - Control wells: Plate3
Fusion - Control wells: Plate4
Fusion - Control wells: Plate5
Fusion Index (%)
DMSO Well 1
DMSO Well 2
DMSO Well 3
FSK Well 1
FSK Well 2
FSK Well 3
ERVFRD1 Well 1
ERVFRD1 Well 2
GCM1 Well 1
GCM1 Well 2
TRAC Well 1
TRAC Well 2
F
hCG - Control wells: Plate1
hCG - Control wells: Plate2
hCG - Control wells: Plate3
hCG - Control wells: Plate4
hCG - Control wells: Plate5
hCG ELISA Abs. / # nuclei
DMSO
FSK
ERVFRD-1
GCM1
TRAC

**Figure S1. Library gene selection and screen controls. A)** Selection of screen target genes based on expression and GO-term filtering in BeWo RNA-Seq data. **B)** Diagram showing classes of controls included in the screen for both CRISPR Cas9 editing and differentiation phenotypes. **C)** GO ontology of major categories within the 412 gene library and overlap with previously identified essential and growth-restricting genes in hTSCs from Dong, et al. 2022. **D)** Cas9 editing efficiencies in a selection of wells assessed by TIDE PCR sequencing. The Knockout Score from TIDE analysis is shown for individual samples (round circles) and as assessed in wells from the Berkeley screen (crossed square). **E,F)** Raw per-plate **E)** Fusion Index and **F)** hCG secretion normalized to # nuclei per well, measured across individual control wells. Points indicate triplicate measurements in separate plates passaged from a parental well. Error bars are standard deviation.
