## Supplementary material for "Phenotypic CRISPR screening identifies ZBTB10 as a novel regulator of human trophoblast differentiation": Figure S2

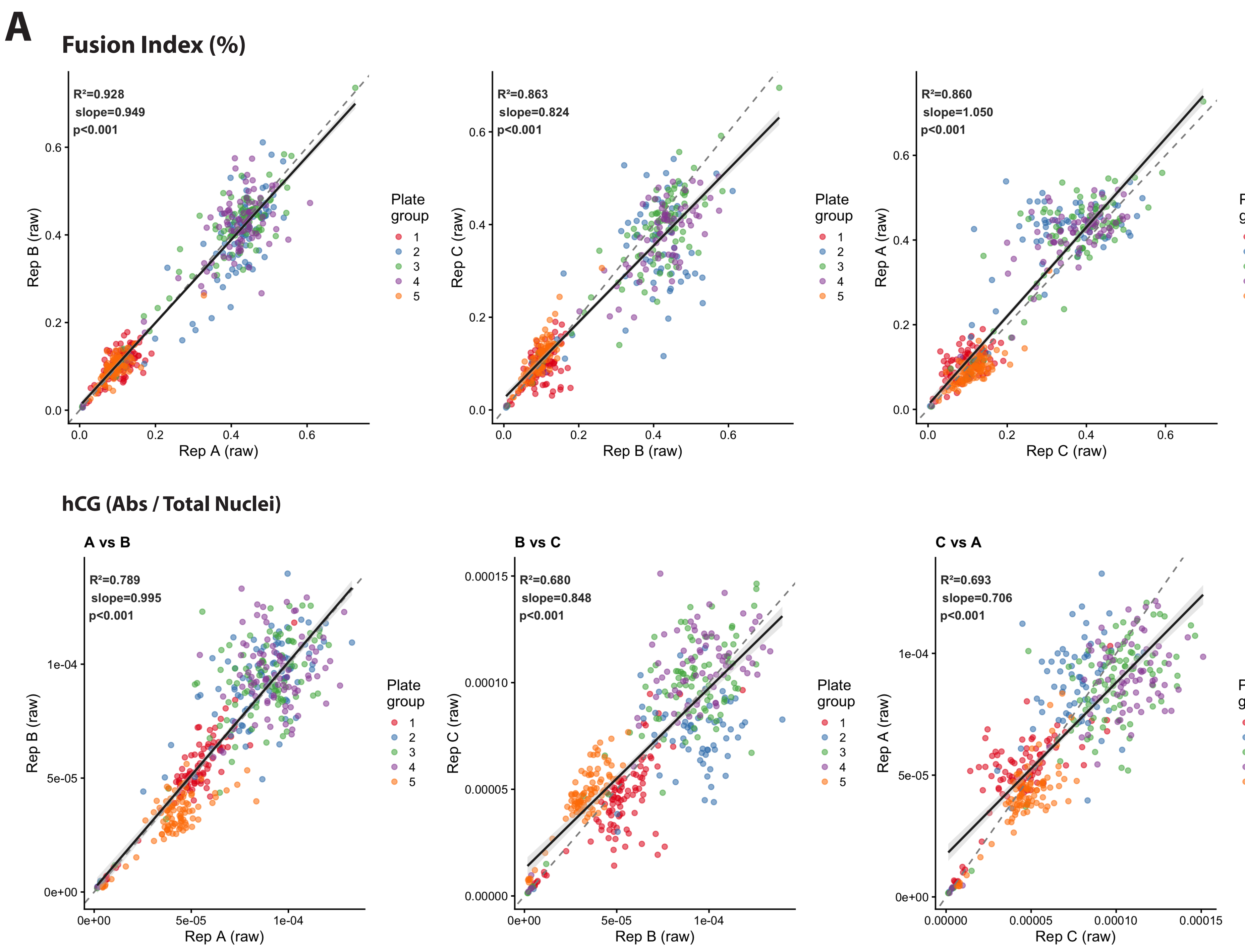


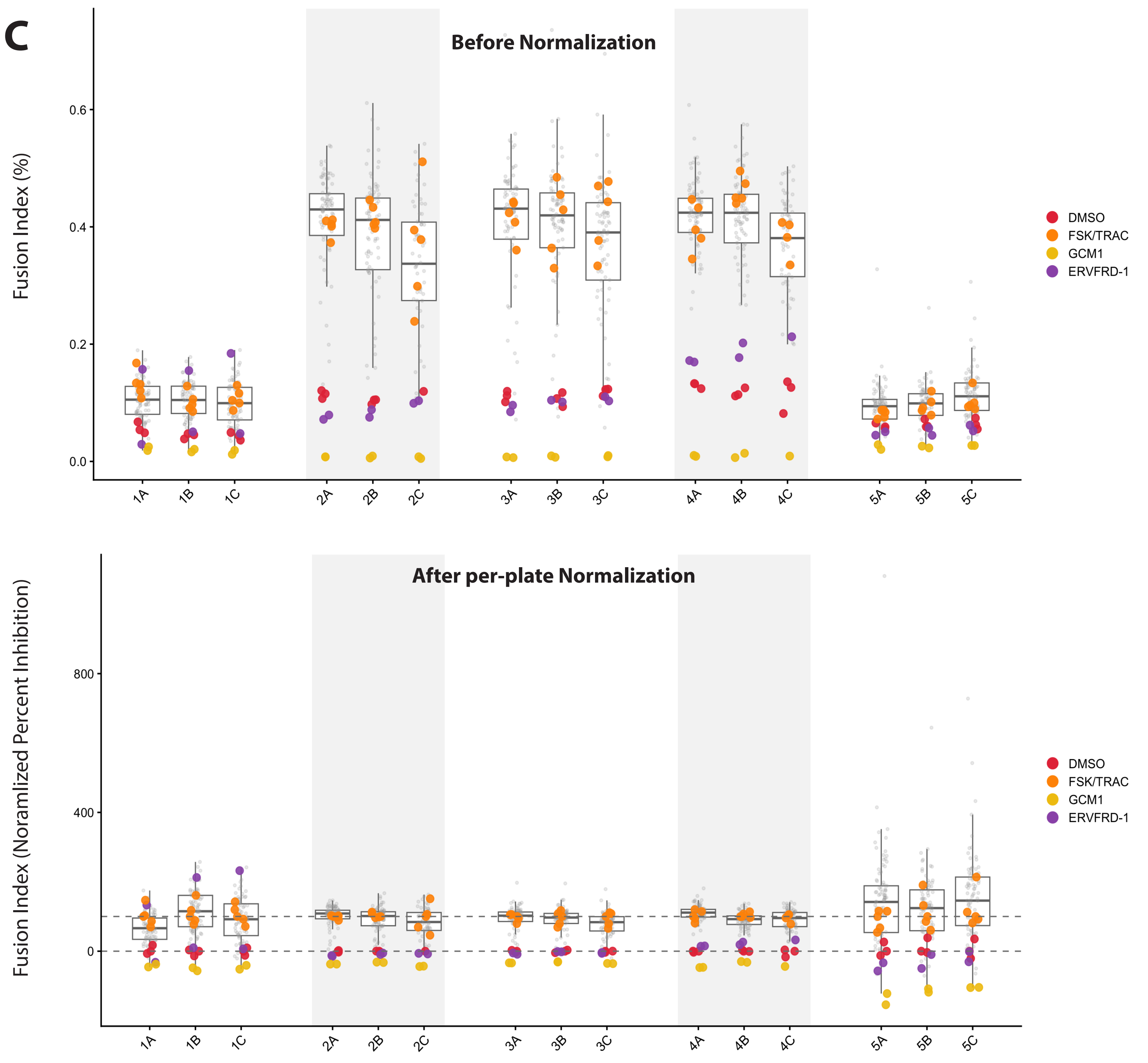


**B**

| Plate | Total wells | Blank wells | Wells Passed | Wells excluded (Viab <45%) | % Plate Excluded | DMSO wells pass | FSK/TRAC wells pass | Included in Cellpaint analysis? |
|---|---|---|---|---|---|---|---|---|
| 1A | 96 | 2 | 94 | 0 | 0.0 | 3 | 5 | Included |
| 1B | 96 | 2 | 94 | 0 | 0.0 | 3 | 5 | Included |
| 1C | 96 | 2 | 94 | 0 | 0.0 | 3 | 5 | Included |
| 2A | 96 | 2 | 94 | 0 | 0.0 | 3 | 5 | Included |
| 2B | 96 | 2 | 94 | 0 | 0.0 | 3 | 5 | Included |
| 2C | 96 | 2 | 68 | 26 | 27.7 | 1 | 5 | Excluded |
| 3A | 96 | 2 | 93 | 1 | 1.1 | 3 | 5 | Included |
| 3B | 96 | 2 | 92 | 2 | 2.1 | 3 | 5 | Included |
| 3C | 96 | 2 | 92 | 2 | 2.1 | 3 | 5 | Included |
| 4A | 96 | 2 | 94 | 0 | 0.0 | 3 | 5 | Included |
| 4B | 96 | 2 | 93 | 1 | 1.1 | 3 | 5 | Included |
| 4C | 96 | 2 | 75 | 19 | 20.2 | 3 | 4 | Included |
| 5A | 96 | 3 | 93 | 0 | 0.0 | 3 | 5 | Included |
| 5B | 96 | 3 | 93 | 0 | 0.0 | 3 | 5 | Included |
| 5C | 96 | 3 | 93 | 0 | 0.0 | 3 | 5 | Included |

Wells were excluded if fluorescent nuclei/total nuclei <45% indicating mostly dead cells.
Plate excluded from Cellpaint analysis if fewer than 3 control wells (DMSO or FSK/TRAC) passed viability filters.

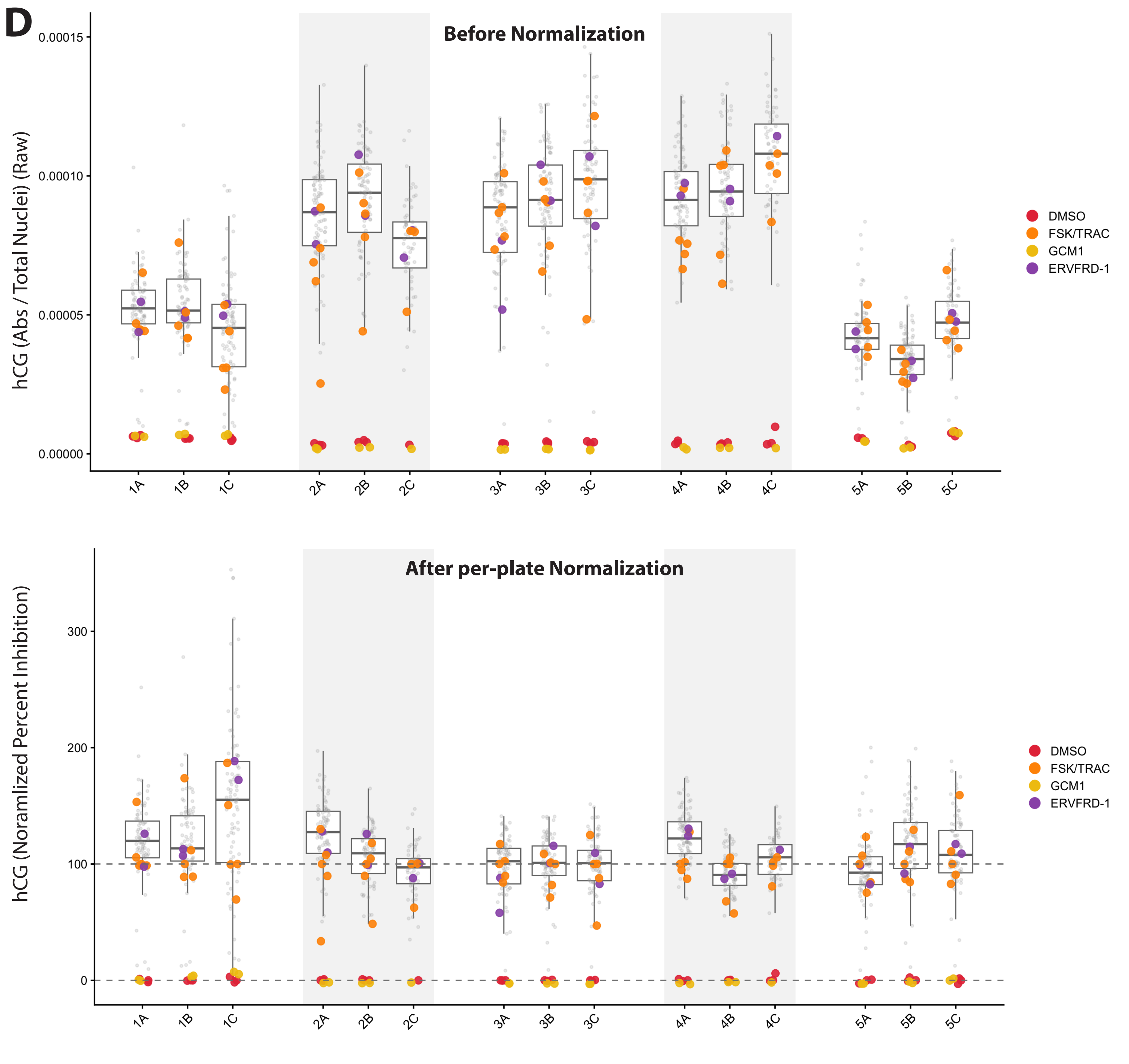

**Figure S2. Screen quality control and normalization. A)** Replicate correlation of raw Fusion Index and hCG secretion across individual replicate plates (Three replicates indicated as A, B, C) prior to normalization. A linear regression model fit (black line with grey shading representing 95% confidence intervals; unit line of slope=1 shown in dashed grey line), $R^2$ correlation values, slope of linear fit, and p-value from t-test indicates all replicate correlations are significantly different from zero. **B)** Table summarizing per-well and per-plate quality control metrics and criteria for exclusion from downstream analysis. **C,D)** Primary phenotypes **C)** Fusion Index and **D)** hCG secretion (normalized to # nuclei per well) per replicate plate across the screen (1A, 1B, 1C indicate 3 triplicate plates passaged from one parental nucleofection plate). Top panel shows raw values and bottom panel shows after per-plate percent inhibition normalization. Dashed lines represent the percent inhibition normalization where the median DMSO wells are set to 0% and median FSK/TRAC wells are set to 100% per plate. Control wells included on every plate (DMSO, FSK/TRAC, GCM1, ERVFRD-1) are shown as large colored dots and all sample wells are shown as smaller grey dots.
