## Supplementary material for "Phenotypic CRISPR screening identifies ZBTB10 as a novel regulator of human trophoblast differentiation": Figure S3

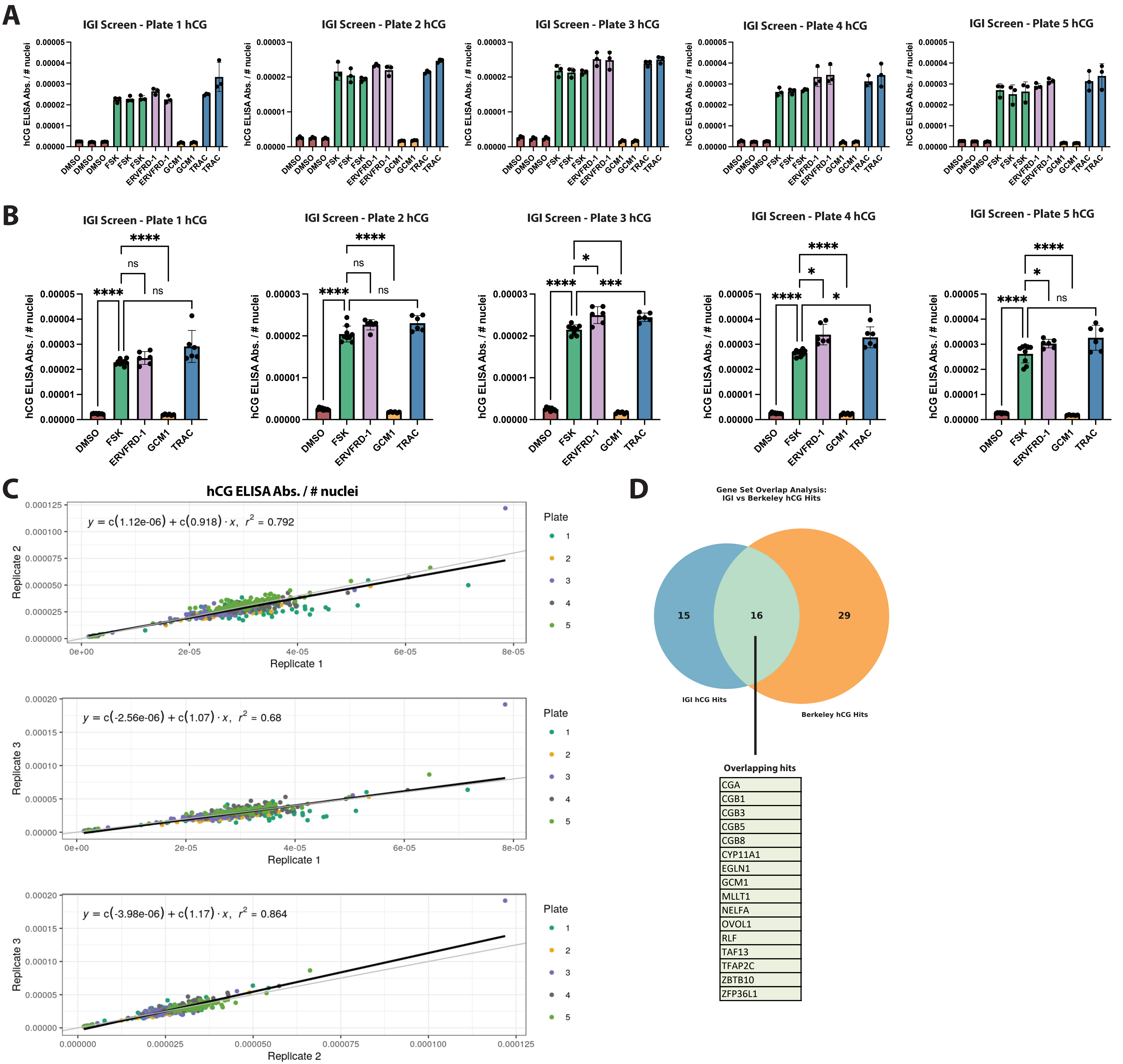
A
IGI Screen - Plate 1 hCG
IGI Screen - Plate 2 hCG
IGI Screen - Plate 3 hCG
IGI Screen - Plate 4 hCG
IGI Screen - Plate 5 hCG
hCG ELISA Abs. / # nuclei
DMSO
FSK
ERVFRD-1
GCM1
TRAC
B
IGI Screen - Plate 1 hCG
IGI Screen - Plate 2 hCG
IGI Screen - Plate 3 hCG
IGI Screen - Plate 4 hCG
IGI Screen - Plate 5 hCG
hCG ELISA Abs. / # nuclei
****
ns
*
***
C
hCG ELISA Abs. / # nuclei
y = c(1.12e-06) + c(0.918) · x, r2 = 0.792
Replicate 2
Replicate 1
Plate
1
2
3
4
5
y = c(-2.56e-06) + c(1.07) · x, r2 = 0.68
Replicate 3
Replicate 1
y = c(-3.98e-06) + c(1.17) · x, r2 = 0.864
Replicate 3
Replicate 2
D
Gene Set Overlap Analysis:
IGI vs Berkeley hCG Hits
15
16
29
IGI hCG Hits
Berkeley hCG Hits
Overlapping hits
CGA
CGB1
CGB3
CGB5
CGB8
CYP11A1
EGLN1
GCM1
MLLT1
NELFA
OVOL1
RLF
TAF13
TFAP2C
ZBTB10
ZFP36L1

**Figure S3. Biological replicate validation of hCG screen results with high-throughput automation at the Innovative Genomics Institute. A)** Raw measurements for individual wells across parental screening plate for hCG absorbance normalized to the number of nuclei per well. Points represent individual measurements for each well across triplicate imaging plates. **B)** Raw measurements for each control category averaged across wells in each plate. Significance bars indicate Brown-Forsythe and Welch ANOVA test with adjusted p-value less than 0.0332 (*), 0.0021 (**), 0.0002 (***), and 0.0001 (****). **C)** Replicate correlation of hCG secretion (normalized to # nuclei per well) across individual replicate plates for all 5 screening plates prior to plate-level normalization. A linear regression model was fit (shown as black line; unit line of slope=1 shown in grey line), $R^2$ correlation values, and equation of linear fit are shown for each comparison. **D)** Venn diagram overlap of gene knockouts called as hits for hCG secretion in either the IGI or Berkeley hCG screens. The 16 overlapping gene hits are shown in the callout table.
