## Supplementary material for "Phenotypic CRISPR screening identifies ZBTB10 as a novel regulator of human trophoblast differentiation": Figure S4

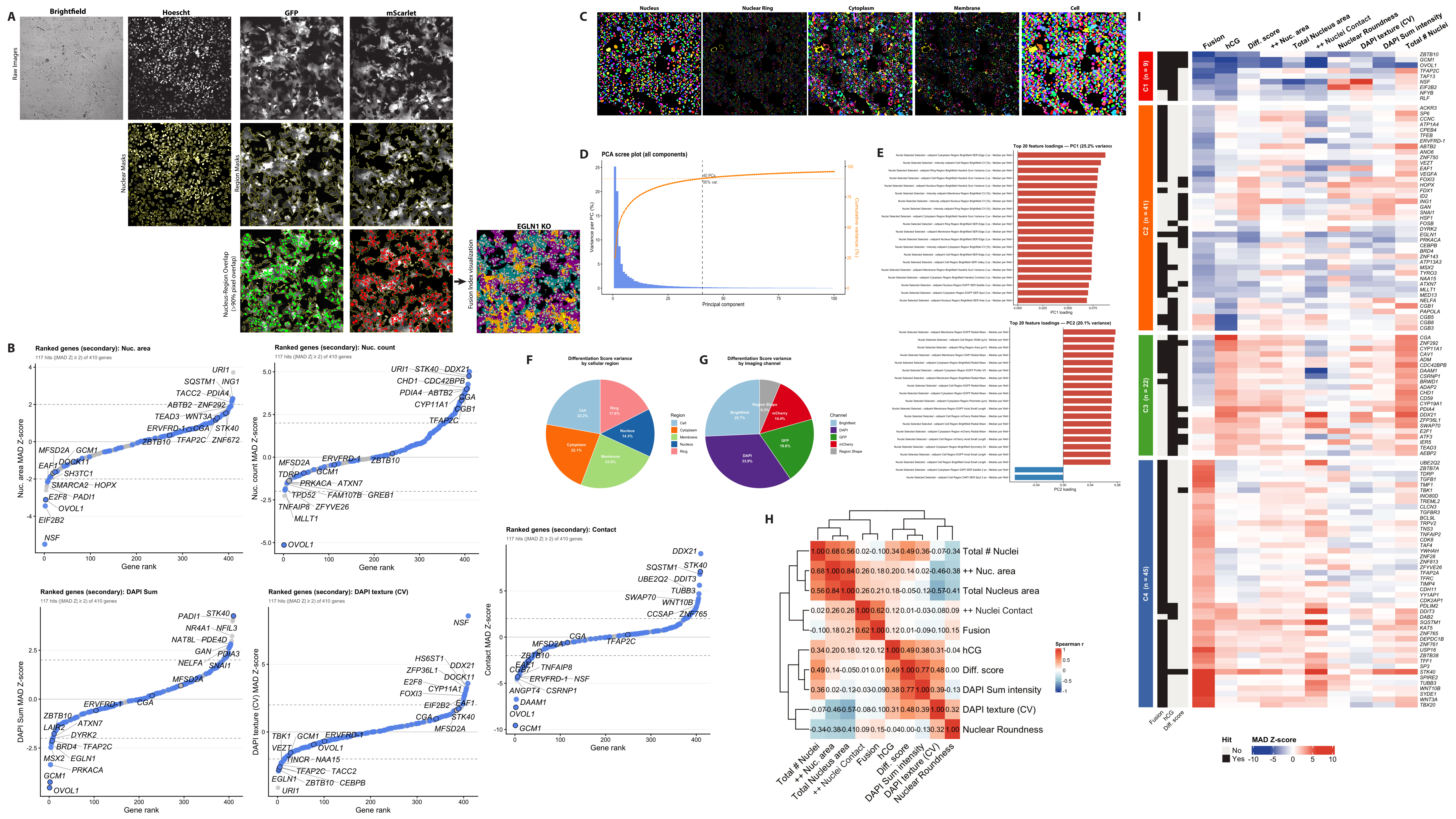

A
Brightfield
Hoescht
GFP
mScarlet
Raw Images
Nuclear Masks
Region Masks
Nucleus-Region Overlap (>90% pixel overlap)
Fusion Index visualization
EGLN1 KO
B
Ranked genes (secondary): Nuc. area
117 hits (|MAD Z| ≥ 2) of 410 genes
Nuc. area MAD Z-score
Gene rank
URI1
SQSTM1
ING1
TACC2
PDIA4
ABTB2
ZNF292
TEAD3
WNT3A
ERVFRD-1
CGA
STK40
ZBTB10
TFAP2C
ZNF672
MFSD2A
GCM1
DOCK11
EAF1
SH3TC1
SMARCA2
HOPX
E2F8
PADI1
OVOL1
EIF2B2
NSF
Ranked genes (secondary): Nuc. count
117 hits (|MAD Z| ≥ 2) of 410 genes
Nuc. count MAD Z-score
Gene rank
URI1
STK40
DDX21
CHD1
CDC42BPB
PDIA4
ABTB2
CGA
CYP11A1
CGB1
TFAP2C
MFSD2A
ERVFRD-1
ZBTB10
TDRP
GCM1
PRKACA
ATXN7
TPD52
FAM107B
GREB1
TNFAIP8
ZFYVE26
MLLT1
OVOL1
Ranked genes (secondary): DAPI Sum
117 hits (|MAD Z| ≥ 2) of 410 genes
DAPI Sum MAD Z-score
Gene rank
PADI1
STK40
NR4A1
NFIL3
NAT8L
PDE4D
GAN
PDIA3
NELFA
SNAI1
MFSD2A
ERVFRD-1
CGA
ZBTB10
ATXN7
LAIR2
DYRK2
BRD4
TFAP2C
MSX2
EGLN1
PRKACA
GCM1
OVOL1
Ranked genes (secondary): DAPI texture (CV)
117 hits (|MAD Z| ≥ 2) of 410 genes
DAPI texture (CV) MAD Z-score
Gene rank
NSF
HS6ST1
DDX21
ZFP36L1
DOCK11
E2F8
CYP11A1
FOXI3
EIF2B2
EAF1
CGA
STK40
MFSD2A
TBK1
GCM1
ERVFRD-1
VEZT
OVOL1
TINCR
NAA15
TFAP2C
TACC2
EGLN1
ZBTB10
CEBPB
URI1
Ranked genes (secondary): Contact
117 hits (|MAD Z| ≥ 2) of 410 genes
Contact MAD Z-score
Gene rank
DDX21
SQSTM1
STK40
UBE2Q2
DDIT3
TUBB3
SWAP70
WNT10B
CCSAP
ZNF765
CGA
TFAP2C
MFSD2A
ZBTB10
EAF1
CGB7
TNFAIP8
ERVFRD-1
NSF
ANGPT4
CSRNP1
DAAM1
OVOL1
GCM1
C
Nucleus
Nuclear Ring
Cytoplasm
Membrane
Cell
D
PCA scree plot (all components)
Variance per PC (%)
Cumulative variance (%)
Principal component
40 PCs
90% var.
E
Top 20 feature loadings — PC1 (25.2% variance)
PC1 loading
Top 20 feature loadings — PC2 (20.1% variance)
PC2 loading
F
Differentiation Score variance by cellular region
Cell 22.2%
Ring 17.5%
Nucleus 14.3%
Membrane 23.9%
Cytoplasm 22.1%
Region
Cell
Cytoplasm
Membrane
Nucleus
Ring
G
Differentiation Score variance by imaging channel
Brightfield 25.7%
Region Shape 6.3%
mCherry 14.4%
GFP 19.8%
DAPI 33.9%
Channel
Brightfield
DAPI
GFP
mCherry
Region Shape
H
Total # Nuclei
++ Nuc. area
Total Nucleus area
++ Nuclei Contact
Fusion
hCG
Diff. score
DAPI Sum intensity
DAPI texture (CV)
Nuclear Roundness
Spearman r
I
C1 (n = 9)
C2 (n = 41)
C3 (n = 22)
C4 (n = 45)
Fusion
hCG
Diff. score
++ Nuc. area
Total Nucleus area
++ Nuclei Contact
Nuclear Roundness
DAPI texture (CV)
DAPI Sum intensity
Total # Nuclei
Hit
No
Yes
MAD Z-score

**Figure S4. Segmentation and in-depth phenotyping of screen wells. A)** Example Fusion Index segmentation pipeline using Perkin Elmer Harmony software. Four channel images are acquired and region masks (segmented nuclei or thresholded GFP and mScarlet regions) are shown as yellow outlines below. Nuclei with >90% pixel overlap with each region are shown in green (GFP+ nuclei) or red (mScarlet+ nuclei). A final merged segmentation overlay as in Fig. 1D for the same field of view. The segmented GFP+ region is displayed in teal, the segmented mScarlet+ region is displayed in magenta, the overlap between GFP and mScarlet regions is shown in orange, double positive nuclei (>90% pixel overlap with both GFP and mScarlet regions) are highlighted in yellow with total fluorescent nuclei are shown in gray. **B)** Ranked gene plots showing called hits for any primary phenotype in blue overlaid on ranked plots for example secondary phenotypes with highlighted genes annotated. **C)** Example region masks annotated in the Cell Paint analysis (shown for DMSO control). **D)** Principal component analysis scree plot showing cumulative variance explained for summed principal components in the CellPaint analysis. **E)** Top 20 feature loadings for the major principal components (PC1, PC2) from the Cell Paint analysis. **F,G)** The percent variance of the Differentiation Score vector explained by the sum of different feature types including **F)** cellular region and **G)** imaging channel. **H)** Spearman correlation between different primary and secondary phenotype readouts. **I)** Expanded heatmap from Fig. 2G, showing robust Z-scores (winsorised at ±10) for primary and secondary phenotypes for all genes that are hits in at least one phenotype with their cluster assignments. Secondary phenotype Z-scores (nuclear morphology, cell contact, DAPI texture, etc.) are displayed for biological comparison but did not influence cluster assignment.
