## Supplementary material for "Phenotypic CRISPR screening identifies ZBTB10 as a novel regulator of human trophoblast differentiation": Figure S5

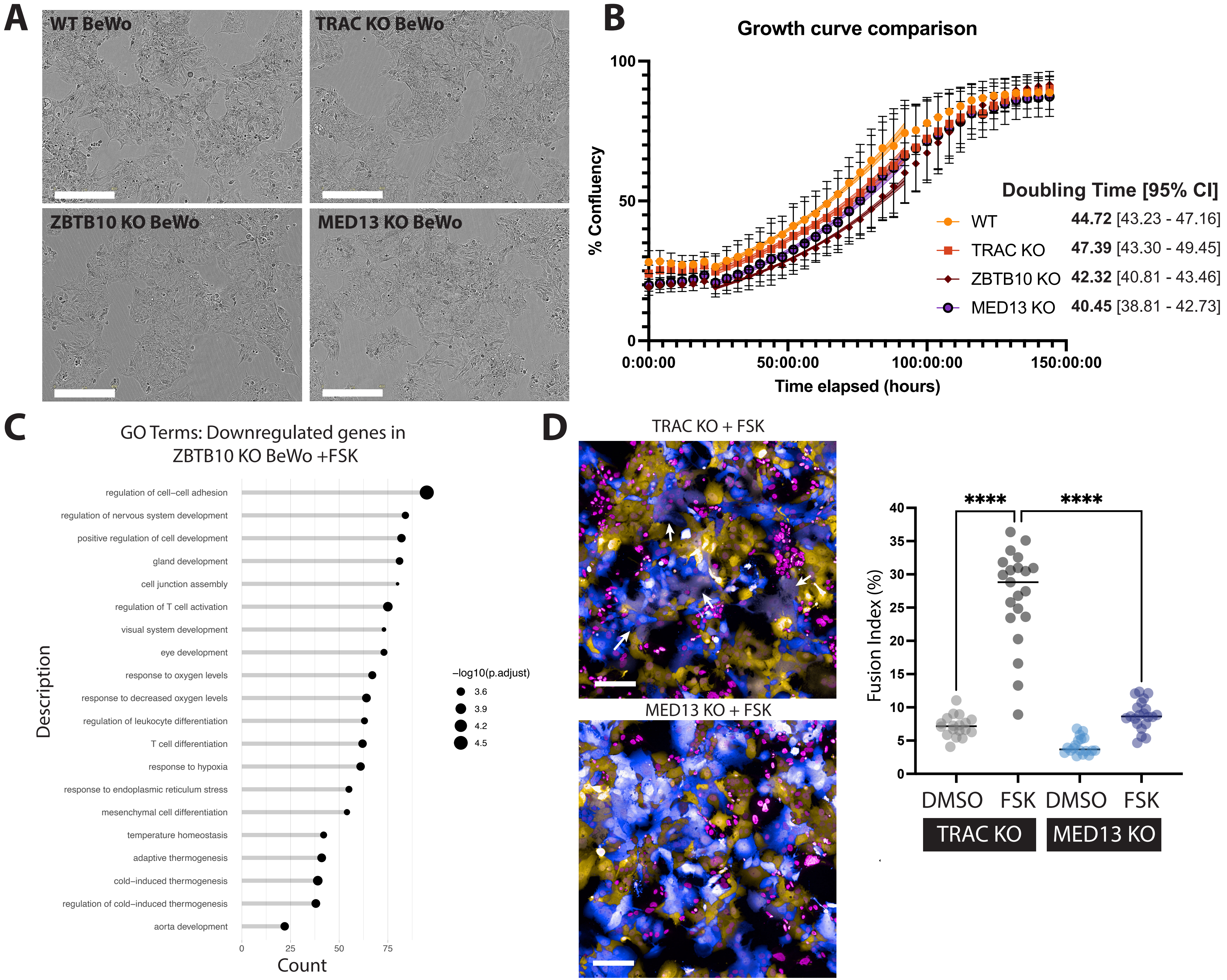

A
WT BeWo
TRAC KO BeWo
ZBTB10 KO BeWo
MED13 KO BeWo
B
Growth curve comparison
% Confluency
Time elapsed (hours)
0:00:00
50:00:00
100:00:00
150:00:00
Doubling Time [95% CI]
WT 44.72 [43.23 - 47.16]
TRAC KO 47.39 [43.30 - 49.45]
ZBTB10 KO 42.32 [40.81 - 43.46]
MED13 KO 40.45 [38.81 - 42.73]
C
GO Terms: Downregulated genes in ZBTB10 KO BeWo +FSK
regulation of cell−cell adhesion
regulation of nervous system development
positive regulation of cell development
gland development
cell junction assembly
regulation of T cell activation
visual system development
eye development
response to oxygen levels
response to decreased oxygen levels
regulation of leukocyte differentiation
T cell differentiation
response to hypoxia
response to endoplasmic reticulum stress
mesenchymal cell differentiation
temperature homeostasis
adaptive thermogenesis
cold−induced thermogenesis
regulation of cold−induced thermogenesis
aorta development
Description
Count
−log10(p.adjust)
3.6
3.9
4.2
4.5
D
TRAC KO + FSK
MED13 KO + FSK
Fusion Index (%)
****
****
DMSO FSK DMSO FSK
TRAC KO
MED13 KO

**Figure S5. BeWo growth curves and validation of MED13 impacts on fusion. A)** Brightfield images of control and polyclonal knockout BeWo cells. Scale bars are 400μm. **B)** Cell growth curves and exponential growth curve fits for polyclonal knockout BeWo cells. The 95% confidence intervals for fit curves (transparent brackets) are shown across 6-8 wells per experiment for at least 3 independent biological experiments. **C)** GO ontology of differential gene expression analysis of down-regulated genes (FDR<0.05, Log Fold-Change <0) for Forskolin-treated ZBTB10 KO BeWo cells versus Forskolin-treated wild-type BeWo cells (genes higher in WT). **D)** Example images of BeWo fusion assays in polyclonal TRAC KO (control) versus MED13 KO mCherry and GFP BeWo co-cultures. GFP BeWos (blue), mCherry BeWos (yellow) and Hoechst-labeled nuclei (magenta) are shown with white arrows indicating examples of fused regions. Quantified Fusion Index across individual wells from at least 3 independent biological experiments is shown where comparisons indicate a One-way ANOVA with mixed-effects analysis, Geisser-Greenhouse correction, and Dunnett's multiple comparisons test and **** = adjusted p-value <0.0001.
