## Supplementary material for "Phenotypic CRISPR screening identifies ZBTB10 as a novel regulator of human trophoblast differentiation": Figure S6

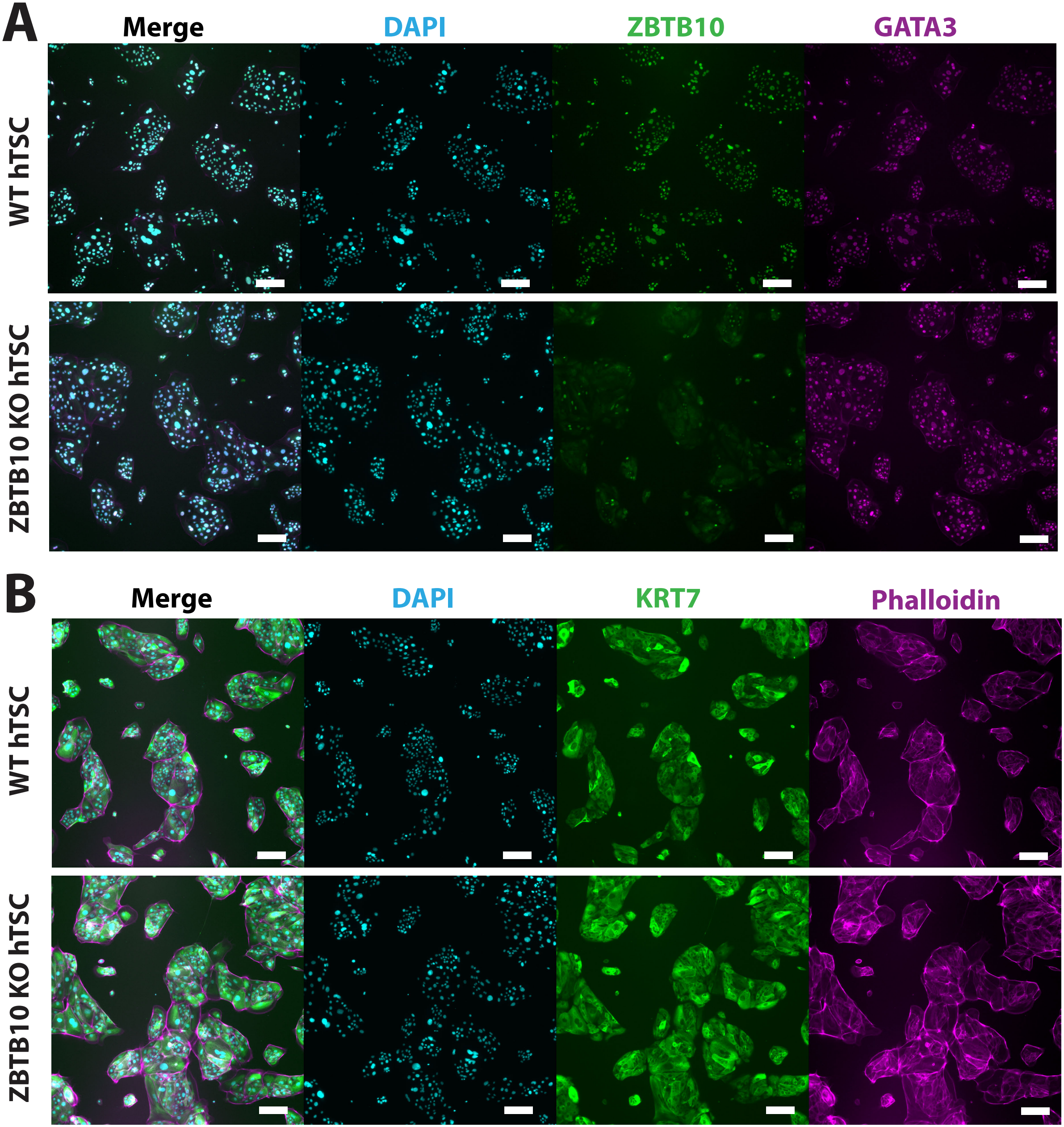
A
Merge
DAPI
ZBTB10
GATA3
WT hTSC
ZBTB10 KO hTSC
B
Merge
DAPI
KRT7
Phalloidin
WT hTSC
ZBTB10 KO hTSC

**Figure S6. ZBTB10 knockout does not perturb trophoblast stem cell marker expression. A-B)** Immunofluorescent staining of ZBTB10, TSC marker genes, and cell morphology (Phalloidin) in undifferentiated negative control (NEG) and ZBTB10 KO hTSCs. Scale bars are 200μm.
