## Supplementary material for "Phenotypic CRISPR screening identifies ZBTB10 as a novel regulator of human trophoblast differentiation": Figure S7

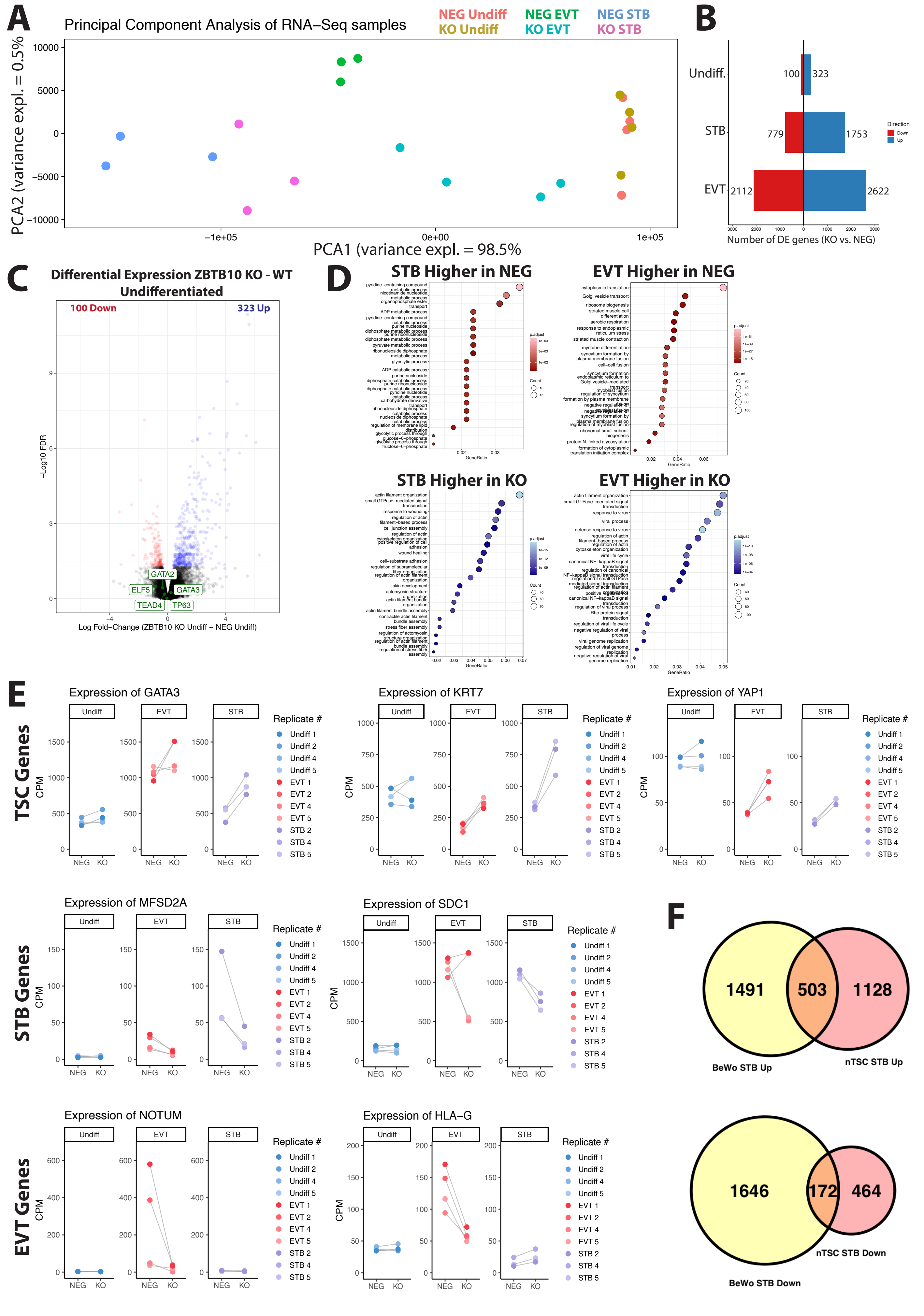

A
Principal Component Analysis of RNA−Seq samples
NEG Undiff
KO Undiff
NEG EVT
KO EVT
NEG STB
KO STB
PCA2 (variance expl. = 0.5%
PCA1 (variance expl. = 98.5%
B
Undiff.
STB
EVT
100
323
779
1753
2112
2622
Direction
Down
Up
Number of DE genes (KO vs. NEG)
C
Differential Expression ZBTB10 KO - WT Undifferentiated
100 Down
323 Up
−Log10 FDR
GATA2
ELF5
GATA3
TEAD4
TP63
Log Fold−Change (ZBTB10 KO Undiff − NEG Undiff)
D
STB Higher in NEG
EVT Higher in NEG
STB Higher in KO
EVT Higher in KO
GeneRatio
p.adjust
Count
E
TSC Genes
STB Genes
EVT Genes
Expression of GATA3
Expression of KRT7
Expression of YAP1
Expression of MFSD2A
Expression of SDC1
Expression of NOTUM
Expression of HLA−G
Undiff
EVT
STB
CPM
NEG
KO
Replicate #
Undiff 1
Undiff 2
Undiff 4
Undiff 5
EVT 1
EVT 2
EVT 4
EVT 5
STB 2
STB 4
STB 5
F
1491
503
1128
BeWo STB Up
nTSC STB Up
1646
172
464
nTSC STB Down
BeWo STB Down

**Figure S7. Extended analysis of ZBTB10 KO RNA-Seq in trophoblast stem cells. A)** Principal component analysis of negative control (NEG) and ZBTB10 KO hTSC RNA-Seq samples. Three to four replicates per condition are shown in corresponding colors. **B)** Differential gene expression analysis comparing the number of differentially expressed genes in NEG control versus ZBTB10 KO hTSCs in each differentiation condition. **C)** RNA-Seq differential gene expression analysis comparing undifferentiated ZBTB10 KO versus NEG hTSCs. Significantly up-regulated genes (FDR<0.05, Log Fold-Change >0) are shown in blue, and significantly down-regulated genes (FDR<0.05, Log Fold-change <0) are shown in red. Unchanged genes are in black. **D)** GO ontology of differential gene expression analysis of up-regulated (FDR<0.05, Log Fold-Change >0) and down-regulated genes (FDR<0.05, Log Fold-Change <0) for STB and EVT differentiated ZBTB10 KO versus NEG hTSCs. **E)** RNA-Seq CPM counts for select marker genes showing individual changes per-replicate between NEG and ZBTB10 KO hTSCs. **F)** Venn diagram of genes regulated by ZBTB10 in both BeWo and hTSCs. BeWo STB: Differentially expressed genes in Forskolin-treated ZBTB10 KO BeWo versus Forskolin-treated WT BeWo with FDR <0.05 (Up: Log Fold-Change>0, Down: Log Fold-Change<0). hTSC STB: Differentially expressed genes in Day 6 STB-differentiated ZBTB10 KO hTSC versus Day 6 STB-differentiated NEG hTSCs with FDR <0.05 (Up: Log Fold-Change>0, Down: Log Fold-Change<0).
