## Supplementary material for "Phenotypic CRISPR screening identifies ZBTB10 as a novel regulator of human trophoblast differentiation": Figure S8

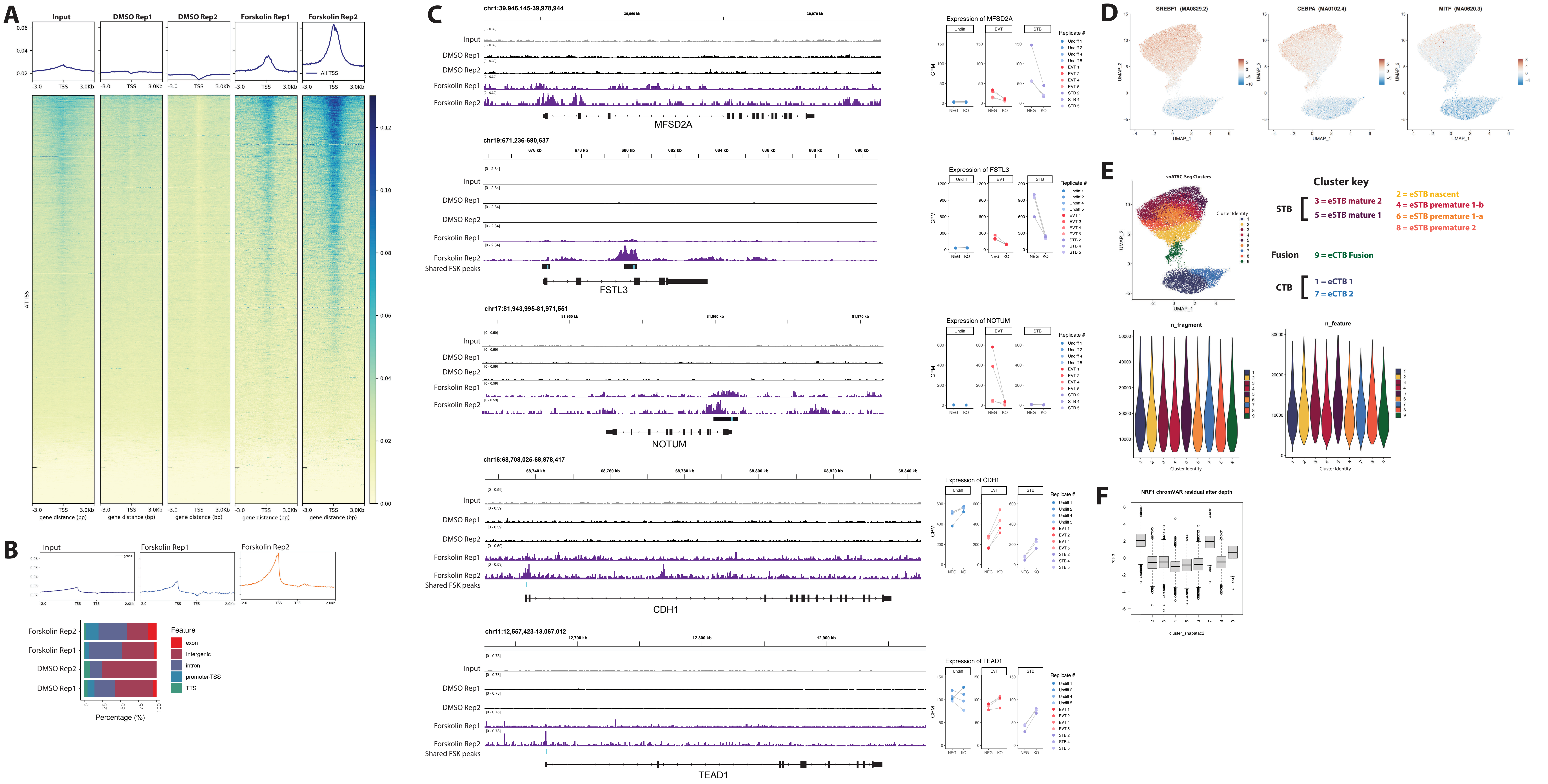

A
Input
DMSO Rep1
DMSO Rep2
Forskolin Rep1
Forskolin Rep2
All TSS
gene distance (bp)
B
Input
Forskolin Rep1
Forskolin Rep2
Forskolin Rep2
Forskolin Rep1
DMSO Rep2
DMSO Rep1
Feature
exon
Intergenic
intron
promoter-TSS
TTS
Percentage (%)
C
chr1:39,946,145-39,978,944
MFSD2A
Expression of MFSD2A
chr19:671,236-690,637
Shared FSK peaks
FSTL3
Expression of FSTL3
chr17:81,943,995-81,971,551
NOTUM
Expression of NOTUM
chr16:68,708,025-68,878,417
CDH1
Expression of CDH1
chr11:12,557,423-13,067,012
TEAD1
Expression of TEAD1
Replicate #
D
SREBF1 (MA0829.2)
CEBPA (MA0102.4)
MITF (MA0620.3)
E
snATAC-Seq Clusters
Cluster Identity
Cluster key
STB
3 = eSTB mature 2
5 = eSTB mature 1
2 = eSTB nascent
4 = eSTB premature 1-b
6 = eSTB premature 1-a
8 = eSTB premature 2
Fusion
9 = eCTB Fusion
CTB
1 = eCTB 1
7 = eCTB 2
n_fragment
n_feature
F
NRF1 chromVAR residual after depth
cluster_snapatac2

**Figure S8. Extended analysis of BeWo ZBTB10 ChIP and snATAC-Seq analysis of ZBTB10 motifs in human first trimester placenta. A)** Heat maps of ZBTB10 enrichment in two replicates each of DMSO-treated and FSK-treated BeWo ChIP-Seq sample genome-wide showing ±3000 bp centered around gene transcription start sites (TSS). **B)** Line plot average of ZBTB10 enrichment aligned my genomic TSS and transcription end sites (TES). Below, the distribution of ZBTB10 peaks mapped to different types of genomic elements shown. **C)** Example genome browser tracks of ZBTB10 ChIP in BeWo cells at STB, EVT, and TSC marker genes. Consensus ZBTB10 peaks identified in both Forskolin replicates are shown with a black bar and the peak center indicated by a teal bar overlay. At right, CPM values from individual replicates of RNA-Seq in WT vs. ZBTB10 KO samples are shown. **D)** Mapping of other indicated HOMER motif activity scores calculated from published snATAC-Seq data from first trimester placenta (Wang et al. 2024[59]). **E)** UMAP of first trimester placenta snATAC-Seq (Wang et al. 2024[59]) colored by cluster annotation. Below, the number of fragments and number of features per cluster are shown. **F)** The calculated chromVAR residual from snATAC-Seq at NRF1 motifs across clusters after normalizing out read depth still shows enrichment for higher activity in CTB and fusing CTB clusters 1,7,9.
