## Supplementary material for "Phenotypic CRISPR screening identifies ZBTB10 as a novel regulator of human trophoblast differentiation": Figure S9

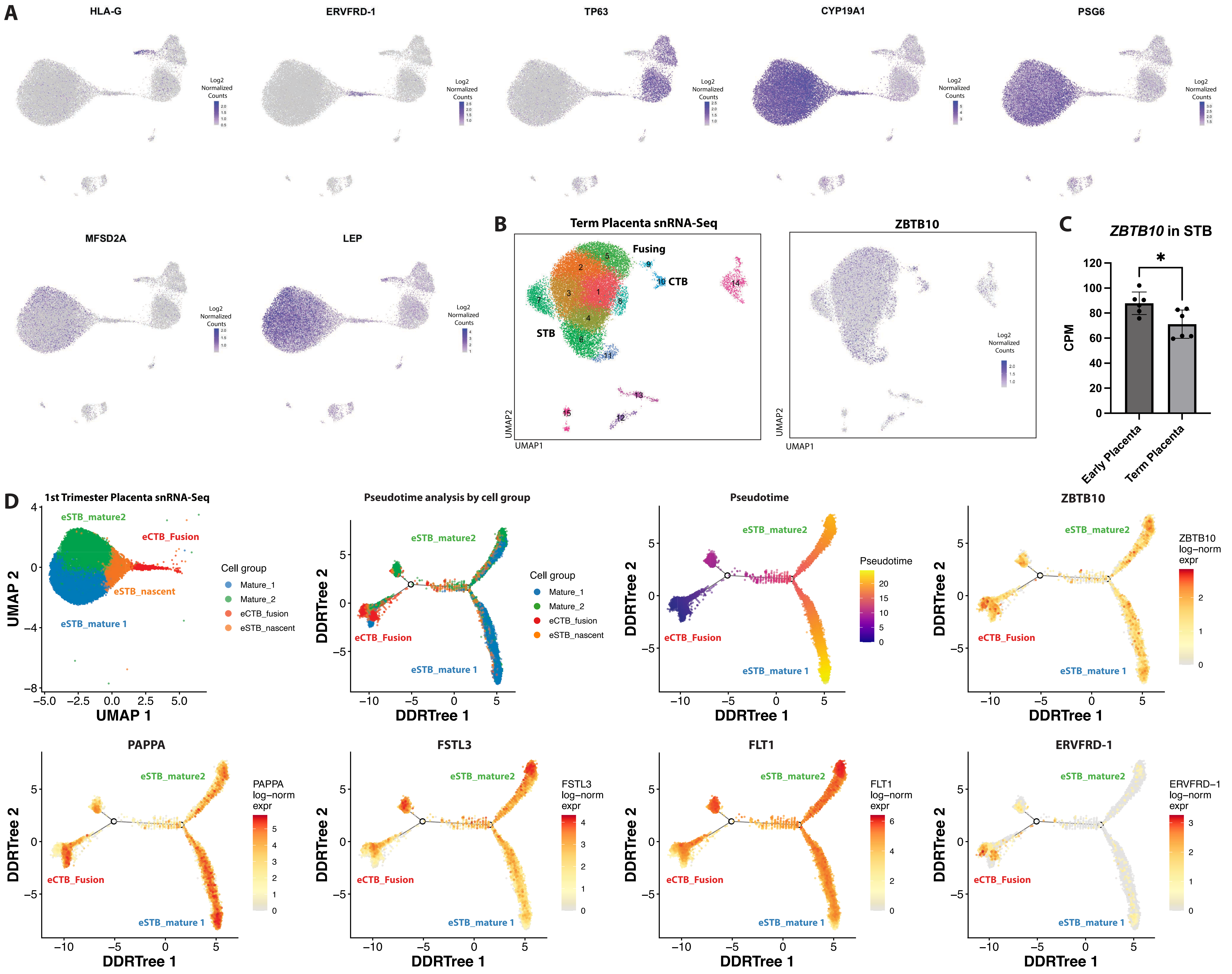
A
HLA-G
ERVFRD-1
TP63
CYP19A1
PSG6
MFSD2A
LEP
Log2 Normalized Counts
B
Term Placenta snRNA-Seq
Fusing
CTB
STB
UMAP1
UMAP2
ZBTB10
C
ZBTB10 in STB
CPM
Early Placenta
Term Placenta
D
1st Trimester Placenta snRNA-Seq
eSTB_mature2
eCTB_Fusion
eSTB_nascent
eSTB_mature 1
UMAP 1
UMAP 2
Cell group
Mature_1
Mature_2
eCTB_fusion
eSTB_nascent
Pseudotime analysis by cell group
Pseudotime
DDRTree 1
DDRTree 2
ZBTB10 log−norm expr
PAPPA
PAPPA log−norm expr
FSTL3
FSTL3 log−norm expr
FLT1
FLT1 log−norm expr
ERVFRD-1
ERVFRD−1 log−norm expr

**Figure S9. Cluster validation and pseudotime analysis of first trimester placenta snRNA-Seq. A)** First trimester human placenta snRNA-Seq (Wang et al. 2024[59]) UMAP overlaid with marker genes for annotated clusters. **B)** UMAP for human term placenta (Wang, et al. 2024[59]) with annotated clusters and overlaid ZBTB10 expression. **C)** Average expression of ZBTB10 in pseudobulk STB (all STB clusters) snRNA-Seq of first trimester versus term placenta from Wang, et al. 2024[59]. Points indicate averages from individual donors. Comparison indicates a Welch's t-test where * corresponds to a p-value <0.032. **D)** Pseudo-time analysis of eCTB fusion, nascent CTB, and two mature STB sub-types. Cell cluster annotations (from Wang, et al. 2024[59]) as well as the calculated per-nucleus pseudotime are overlaid on the DDRTree plot. Log-normalized per-nucleus expression of individual marker genes are shown overlaid on the pseudotime plot.
