## Supplementary material for "Phenotypic CRISPR screening identifies ZBTB10 as a novel regulator of human trophoblast differentiation": Figure S10

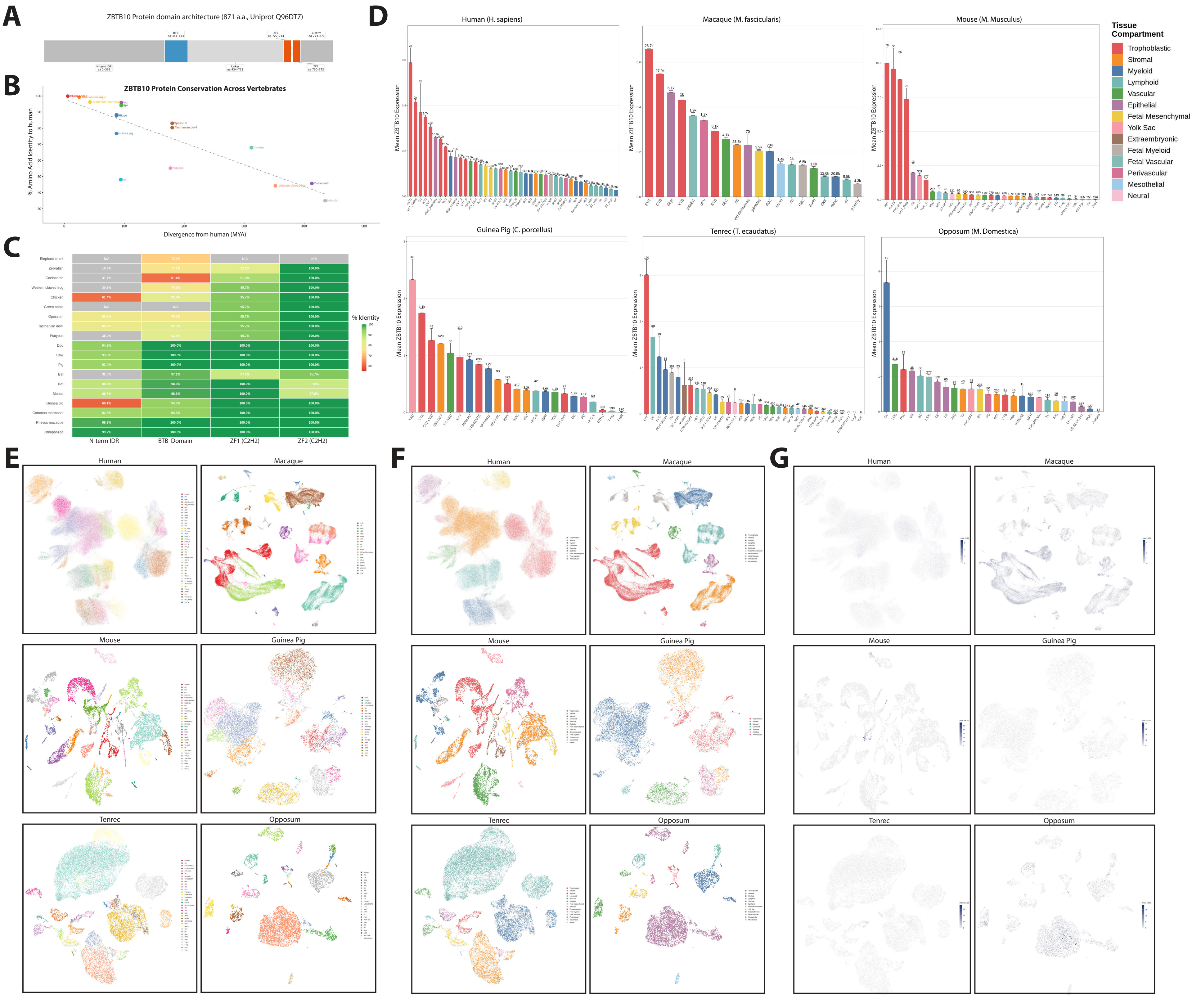
A
ZBTB10 Protein domain architecture (871 a.a., Uniprot Q96DT7)
B
ZBTB10 Protein Conservation Across Vertebrates
% Amino Acid Identity to human
Divergence from human (MYA)
C
% Identity
N-term IDR
BTB Domain
ZF1 (C2H2)
ZF2 (C2H2)
D
Human (H. sapiens)
Macaque (M. fascicularis)
Mouse (M. Musculus)
Guinea Pig (C. porcellus)
Tenrec (T. ecaudatus)
Opposum (M. Domestica)
Mean ZBTB10 Expression
Tissue Compartment
Trophoblastic
Stromal
Myeloid
Lymphoid
Vascular
Epithelial
Fetal Mesenchymal
Yolk Sac
Extraembryonic
Fetal Myeloid
Fetal Vascular
Perivascular
Mesothelial
Neural
E
Human
Macaque
Mouse
Guinea Pig
Tenrec
Opposum
F
Human
Macaque
Mouse
Guinea Pig
Tenrec
Opposum
G
Human
Macaque
Mouse
Guinea Pig
Tenrec
Opposum

**Figure S10. Evolutionary analysis of ZBTB10 coding sequence and cell type-specific expression. A)** Schematic of ZBTB10 annotated protein domains. **B)** Amino acid identity across the entire ZBTB10 protein for annotated ZBTB10 orthologs, colored by clade. **C)** Domain-level identity conservation of ZBTB10 compared to human. **D)** Pseudobulk expression of ZBTB10 transcript levels in annotated cell clusters across six mammalian species (Stadtmauer, et al. 2025), colored by cell group type with trophoblasts indicated in red and myeloid lineages in blue. **E)** Per-species UMAPs of single-cell and single-nucleus sequencing from six mammalian species (Stadtmauer, et al. 2025), showing annotated cell cluster assignments. **F)** Per-species UMAPs colored by broader cell group types (as colored in S10D). **G)** Per-species UMAPs showing overlaid expression levels of ZBTB10.
